# Lateral Septal Circuits Govern Schizophrenia-Like Effects of Ketamine on Social Behavior

**DOI:** 10.1101/2023.08.08.552372

**Authors:** Ruixiang Wang, Zeru Peterson, Nagalakshmi Balasubramanian, Kanza M. Khan, Michael S. Chimenti, Daniel Thedens, Thomas Nickl-Jockschat, Catherine A. Marcinkiewcz

**Affiliations:** Department of Neuroscience and Pharmacology, University of Iowa, Iowa City, IA; Department of Psychiatry, University of Iowa, Iowa City, IA; Department of Psychological Sciences, Daemen University, Amherst, NY; Bioinformatics Division, Iowa Institute of Human Genetics, Iowa City, IA; Iowa Institute for Biomedical Imaging, Iowa City, IA; Iowa Neuroscience Institute, Iowa City, IA

**Author notes:** Please address all correspondence to: Catherine A. Marcinkiewcz, Department of Neuroscience and Pharmacology 2-430 Bowen Science Building, University of Iowa, IA 52242.

## Abstract

Schizophrenia is marked by poor social functioning that can have a severe impact on quality of life and independence, but the underlying neural circuity is not well understood. Here we used a translational model of subanesthetic ketamine in mice to delineate neural pathways in the brain linked to social deficits in schizophrenia. Mice treated with chronic ketamine (30 mg/kg/day for 10 days) exhibit profound social and sensorimotor deficits as previously reported. Using three-dimensional c-Fos immunolabeling and volume imaging (iDISCO), we show that ketamine treatment resulted in hypoactivation of the lateral septum (LS) in response to social stimuli. Chemogenetic activation of the LS rescued social deficits after ketamine treatment, while chemogenetic inhibition of previously active populations in the LS (i.e. social engram neurons) recapitulated social deficits in ketamine-naïve mice. We then examined the translatome of LS social engram neurons and found that ketamine treatment dysregulated genes implicated in neuronal excitability and apoptosis, which may contribute to LS hypoactivation. We also identified 38 differentially expressed genes (DEGs) in common with human schizophrenia, including those involved in mitochondrial function, apoptosis, and neuroinflammatory pathways. Chemogenetic activation of LS social engram neurons induced downstream activity in the ventral part of the basolateral amygdala, subparafascicular nucleus of the thalamus, intercalated amygdalar nucleus, olfactory areas, and dentate gyrus, and it also reduces connectivity of the LS with the piriform cortex and caudate-putamen. In sum, schizophrenia-like social deficits may emerge via changes in the intrinsic excitability of a discrete subpopulation of LS neurons that serve as a central hub to coordinate social behavior via downstream projections to reward, fear extinction, motor and sensory processing regions of the brain.

## Introduction

Schizophrenia is conceptualized as a neurodevelopmental disorder that often manifests in late adolescence to young adulthood[1], with a younger age of onset being associated with a poorer prognosis and an increased incidence of negative symptoms[2, 3]. Negative affective symptoms, which typically emerge before the onset of schizophrenia[4], can persist throughout the disease and are refractory to conventional antipsychotic treatment[5]. The etiology of this disease and its behavioral manifestations are complex and multi-faceted, confounding prior attempts to link neural activity patterns to specific phenotypes. Symptomatic presentation is also diverse and falls into three main categories: positive symptoms (delusions, hallucinations), negative symptoms (blunt affect, poverty of speech) and cognitive symptoms (disorganized thought, memory loss), all of which can vary in intensity and expression[6]. Negative symptoms also typically co-occur with social deficits[7, 8]. Social behavior is an adaptive function that is conserved across mammalian species, and disruptions in social behavior can have substantial personal, vocational and economic costs that make it difficult for affected individuals to fully integrate into society[9]. However, schizophrenia treatment tends to focus on the so-called positive symptoms which are often salient but less debilitating than negative and cognitive symptoms. Many currently approved drugs primarily target dopamine D_2_ and 5-HT_2A_ receptors[10, 11], which influence positive symptoms but are also associated with motor and cognitive side effects. Despite the urgent need, developing effective treatments for the negative symptoms of schizophrenia and other neurodevelopmental disorders remains a challenge given the complexity of the neural circuits governing affective and social behavior.

The goal of the present study was to elucidate the underlying neural circuitry driving social deficits induced by negative symptoms in schizophrenia utilizing a mouse model of chronic subanesthetic ketamine. Ketamine is an NMDA receptor antagonist with psychotomimetic effects in humans that reproduces the sensorimotor and cognitive deficits associated with schizophrenia[12–15], causing disruptions in thalamocortical networks and increased striatal dopamine capacity[16–18]. NMDA receptor hypofunction has also been reported in schizophrenia patients, providing further face validity for the ketamine model[15, 19].

Specifically, marked reductions in NMDA receptor binding and *Grin1* mRNA encoding the NR1 subunit have been reported in the left hippocampus of schizophrenia patients[20, 21]. In mice and rats, subchronic ketamine is associated with deficits in prepulse inhibition (PPI), social interaction, and cognitive function[22–29], all of which have been reported in schizophrenia.

Another advantage of the chronic ketamine model is that it can be easily executed in transgenic mouse lines, enabling cell-type specific manipulations of circuit function *in vivo*. To gain unbiased insight into the neural circuit basis of social deficits in schizophrenia, we used iDISCO to identify neuronal subpopulations that were differentially active during social investigation (i.e. the social engram) in ketamine-treated mice. We then combined chemogenetics with c-Fos-based targeted recombination in transiently active populations (Fos-TRAP) to interrogate the role of social engram neurons in the lateral septum (LS) in driving social deficits. We further investigated gene networks within the LS social engram that were dysregulated after chronic ketamine, and identified differentially expressed genes (DEGs) that overlapped with human schizophrenia and mouse models of neurodevelopmental disorders (e.g. maternal immune activation, valproic acid). Finally, we leveraged chemogenetics with iDISCO and functional magnetic resonance imaging (fMRI) to better understand how neural activity in the LS social engram modulates downstream brain networks, which may provide further insight into the neural circuitry that orchestrates social deficits in schizophrenia.

## Results

### Three-dimensional mapping of neuronal activity during social stimulation in ketamine- treated mice

Chronic treatment with subanesthetic doses of NMDA receptor antagonists like ketamine have been shown to model various aspects of schizophrenia-like behavior in humans and rodents, including social, sensorimotor and cognitive deficits[12–14, 23–26, 29, 30]. We first demonstrated that chronic ketamine (30 mg/kg/day for 10 days) could induce social deficits in male C57BL/6J mice in the three-chambered social interaction test. Here we found a significant reduction in time spent in social interaction (Stranger x Ketamine interaction: F_1,18_=22.05, p<0.001; Saline-stranger vs. Ketamine-stranger post-test with Šídák’s correction: t_36_=4.57, p<0.001; Saline-empty cage vs. Ketamine-empty cage: t_36_=3.02, p<0.01), although the latency to approaching the novel mouse was not significantly altered. Furthermore, the % time spent in social interaction relative to an empty cage investigation was significantly reduced after chronic ketamine (t_18_=4.64, p<0.001) (Figure 1A-F). In agreement with previous studies in mice, we also observed a significant PPI deficit in ketamine-treated mice (Main effect of ketamine: F_1,17_=5.03, p<0.05) and a significant decrease in %PPI at 82 dB (t_17_=2.13, p<0.05) (Supplementary Figure 1A-C). Furthermore, we did not observe an effect of ketamine on locomotor activity (t_18_=0.03, ns), time spent in the center of the open field (t_17_=0.68, ns) or latency to the center of the open field (t_18_=0.78, ns), which is congruent with a previous study reporting no changes in spontaneous locomotor activity after chronic ketamine[28]. However, there are some reports of hyperactivity after chronic ketamine administration suggesting that this phenotype may depend on other experimental conditions [17, 24] (Supplementary Figure 1D-G).

**Figure 1.**
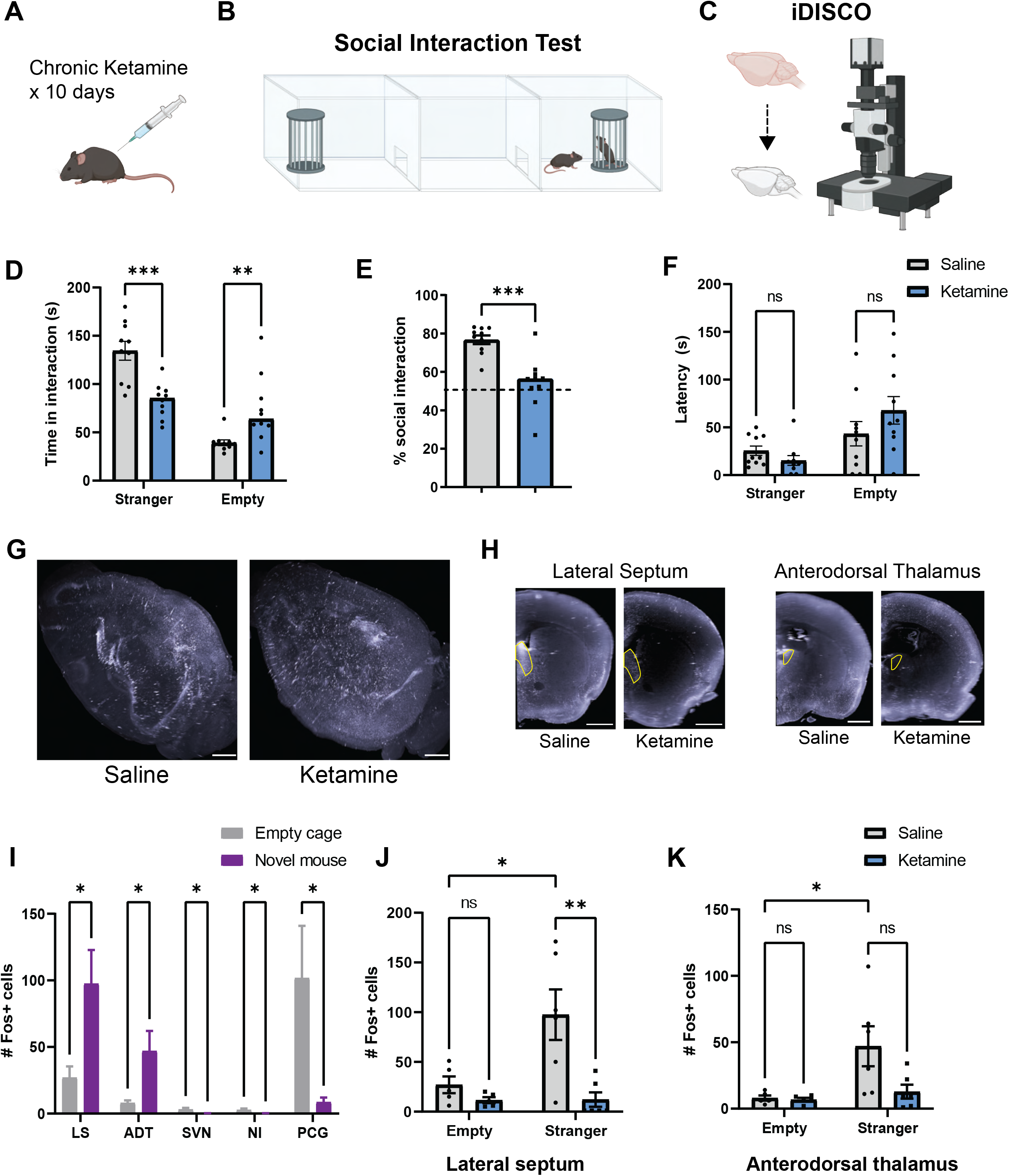
Chronic ketamine leads to reduced social interaction and blunted neural activity in the lateral septum. (A) Experimental schematic of the chronic subanesthetic ketamine model. Mice were administered with ketamine (30 mg/kg/day, i.p.) or 0.9% saline (in the control group) for 10 days. (B) Mice went through a social interaction test in a three-chambered arena. (C) The brains of ketamine-treated mice of a different cohort were cleared and imaged using iDISCO. (D) Histogram showing that the chronic ketamine, versus saline, treatment led to decreased social interaction, displayed by reduced time interacting with stranger mice and increased time investigating the empty cages. (E) The percentage of time engaging in social interaction was also decreased by chronic ketamine exposure. (F) There was no difference in latency to the first social interaction bout between saline and ketamine groups. (G) 3D rendering of c-fos expression in representative iDISCO brains of saline- and ketamine-treated mice (scale bars = 1,000 μm), and (H) coronal views through the lateral septum and anterodorsal thalamus (scale bars = 1,000 μm). (I) Histogram showing that mice exposed to a novel conspecific had higher c-Fos activity in the dorsal part of the lateral septum (LS), the anterodorsal thalamus (ADT), the superior vestibular nucleus (SVN), the nucleus incertus (NI), and the pontine central gray (PCG). (J) Chronic ketamine blunted c-fos activity induced by social interaction in the lateral septum. (K) Chronic ketamine did not lead to a significant reduction in c-fos activity in the anterodorsal thalamus, which was upregulated by social interaction in the saline group. Data are expressed as mean ± SEM. * *p* < 0.05, ** *p* < 0.01, *** *p* < 0.001, ns: non-significant. *Parts of this figure were made using Biorender*.

We utilized iDISCO, a widely-used method for whole-brain c-Fos immunolabeling and volume imaging in solvent-cleared tissues[31], to approximate brain-wide neural activity in response to social stimuli. Here we found that mice exposed to novel social partners had higher c-Fos activity in the dorsal part of the lateral septum (LS), the anterodorsal nucleus of the thalamus (ADT), the superior vestibular nucleus (SVN), nucleus incertus (NI), and pontine central gray (PCG) (Figure 1G-I). Interestingly, activity in the LS was significantly blunted after social interaction in ketamine-treated mice (Stranger x Ketamine Interaction: F_1,18_=5.34, p<0.05, Saline-stranger vs. Ketamine-stranger post-test with Bonferroni correction: t_18_=1.48, p<0.01).

The LS has been previously implicated in social fear and trauma[32–34], but it has not been previously associated with social deficits in schizophrenia. Both ketamine treatment and schizophrenia have been associated with corticothalamic dysconnectivity in humans[16, 18, 19], which prompted us to ask whether activity in the thalamus was altered after chronic ketamine.

Although c-Fos activity was increased in the ADT after social interaction relative to an empty cage in the saline condition (Main effect stranger: F_1,18_= 6.42, p<0.05; Saline-empty vs. Saline- stranger post-test with Bonferroni correction: t_18_=3.10, p<0.05), chronic ketamine did not significantly modify this activity during social interaction (Stranger x Ketamine Interaction: F_1,18_=3.44, ns; Saline-stranger vs. Ketamine-stranger post-test with Bonferroni correction: t_18_=2.85, ns) (Figure 1J-K). However, when saline and ketamine groups in the social interaction condition were analyzed separately, we did see that c-Fos activity was attenuated in the LS and lateral posterior thalamus (LPT) (Supplementary Figure 1H).

### Chemogenetic activation of the LS restores social behavior after chronic ketamine

The LS is a well-established hub that orchestrates complex social behaviors[35], so we decided to interrogate this region as a neural substrate in ketamine-induced social deficits. Here we asked whether inducing neuronal activity in the LS could rescue social deficits after chronic ketamine. Male C57BL/6J mice transduced with a Gq-coupled DREADD (hM3Dq) or a control virus in the LS received daily intraperitoneal injections of saline or ketamine for 10 consecutive days. Mice then underwent clozapine-N-oxide (CNO, 3 mg/kg) injections and behavioral testing in the social interaction test. Here we found that chemogenetic activation of the LS alleviated social deficits at 24 hr after chronic ketamine (Ketamine x DREADD interaction: F_1,33_=51.28, p<0.0001; Saline-mCherry vs Ketamine-mCherry post-test with Bonferroni correction: t_33_=9.98, p<0.0001; Saline-hM3Dq vs. Ketamine-hM3Dq post-test with Bonferroni correction: t_33_=0.549, ns). Ketamine-induced social deficits persisted 12 days later and were relieved by LS activation (Ketamine x DREADD interaction: F_1,16_=20.90, p<0.001; Saline-mCherry vs Ketamine-mCherry post-test with Bonferroni correction: t_16_=7.204, p<0.0001; Saline-hM3Dq vs. Ketamine-hM3Dq post-test with Bonferroni correction: t_16_=0.738, ns). Together, these results suggest that LS activity can restore social behavior in ketamine-treated mice (Figure 2).

**Figure 2.**
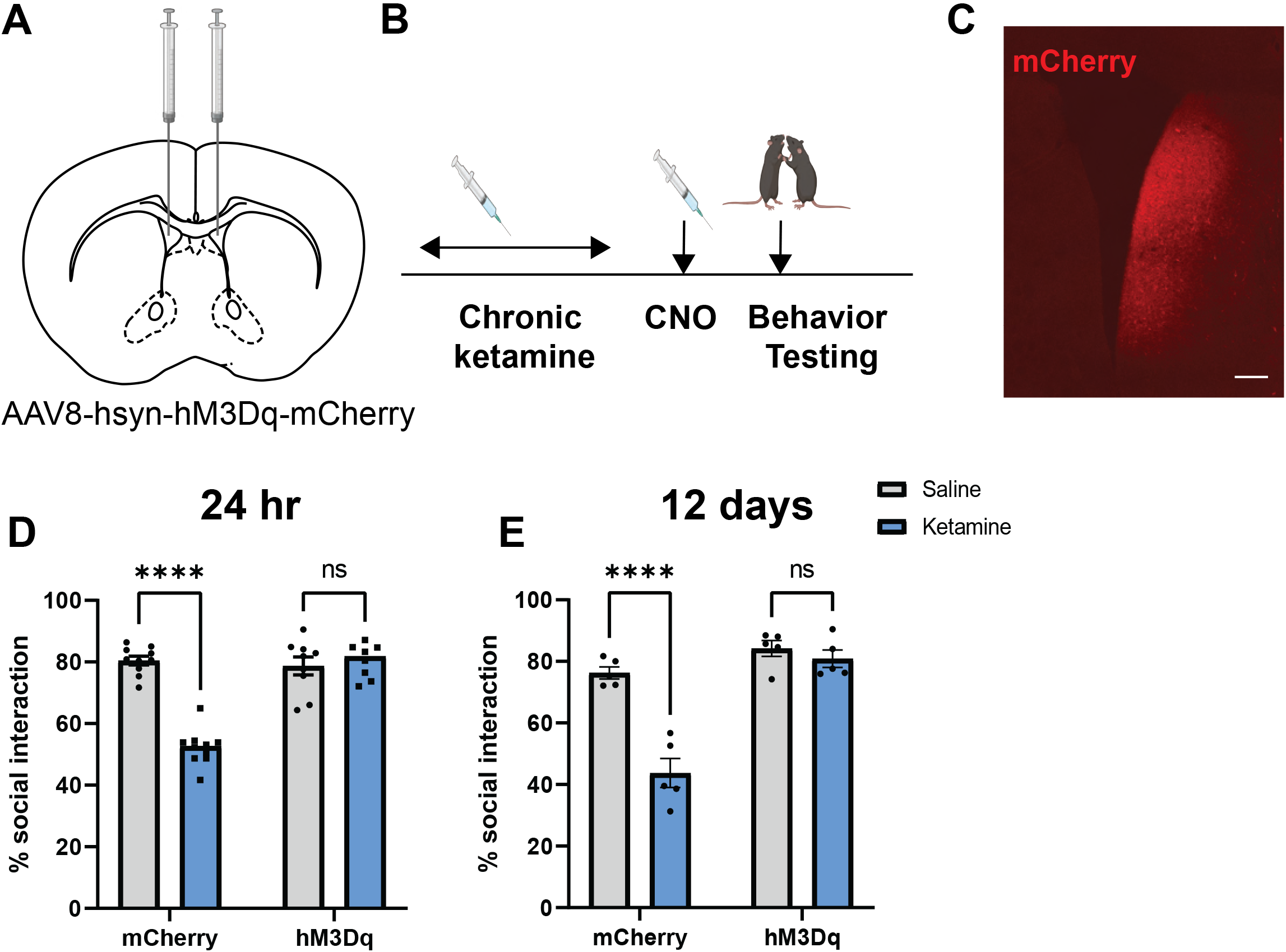
Chemogenetic activation of the lateral septum rescues social deficits induced by chronic ketamine. (A) Experimental schematic of viral targeting of Gq-coupled DREADDs to the lateral septal neurons of C57BL/6J mice. (B) Experimental timeline of the chronic ketamine treatment, chemogenetic activation of lateral septal neurons via clozapine-N-oxide (CNO), and behavioral testing. (C) A representative confocal image of hM3Dq-mCherry expression in the lateral septum (scale bar = 200 μm). (D) Histogram showing that chemogenetic activation of lateral septal neurons reversed social deficits in mice exposed to chronic ketamine 24 hours after the last injections, whereas the deficits were still observed in ketamine mice transduced with only a control (mCherry) virus. (E) Chemogenetic activation of lateral septal neurons still reversed social deficits in ketamine-treated mice 12 days later. Data are expressed as mean ± SEM. **** *p* < 0.0001, ns: non-significant. *Parts of this figure were made using Biorender*.

### A discrete population of social engram neurons in the LS promotes social deficits

LS neurons are a heterogeneous population with a wide range of neurochemical and physiological properties, only a subset of which may be involved in the orchestration of social behavior. Here we genetically tagged LS neurons that were specifically activated by social stimuli using targeted recombination in transiently active populations (TRAP)[36], then chemogenetically inhibited this same neuronal population to interrogate its role in driving social behavior. Fos-iCreER mice were transduced in the LS with Cre-inducible Gi-coupled DREADDs (hM4Di) or a control virus, then exposed to a novel conspecific in the three-chambered social interaction test to induce c-Fos expression in LS social engram neurons. These mice were then injected with 4-OH-tamoxifen (4-OH-TM, 10 mg/kg) to induce Cre-mediated recombination and DREADD expression in the c-Fos-expressing population representing the LS social engram.

Four weeks later, Fos-iCreER::hM4Di^LS^ and Fos-iCreER::mCherry^LS^ mice were injected with CNO and re-assessed in the social interaction test (Figure 3A-C). Here we found that chemogenetic inhibition of LS social engram neurons induced social deficits in male mice (Drug x DREADD Interaction: F_1,13_=96.42, p<0.0001; Baseline-mCherry vs. Baseline-hM4Di post-test with Bonferroni correction: t_26_=0.718, ns; CNO-mCherry vs. CNO-hM4Di: t_26_=10.21, p<0.0001) (Figure 3D). Converging lines of evidence suggest that schizophrenia may arise from different neural pathways in males and females[37], so we asked whether chemogenetic inhibition of the LS social engram could induce social deficits in female mice. In contrast to the males, chemogenetic inhibition of LS social engram neurons did not promote social deficits in females (Drug x DREADD Interaction: F_1,10_=0.862, ns; Baseline-mCherry vs. Baseline-hM4Di post-test with Bonferroni correction: t_20_=0.441, ns; CNO-mCherry vs. CNO-hM4Di: t_20_=2.01, ns) (Figure 3E). We then asked whether chemogenetic inhibition of neurons activated during presentation of two empty cages could elicit the same response. Fos-iCreER::hM4Di^LS^ mice were presented with two empty cages instead of a novel conspecific under one cage during the three-chambered social interaction test prior to injection with 4-OH-TM. Here chemogenetic inhibition of the non-social engram LS neurons did not alter social behavior in these mice (Social stimulus x DREADD interaction: F_1,18_=2.44, ns; Baseline-mCherry vs. Baseline-hM4Di post-test with Bonferroni correction: t_36_=0.94, ns; CNO-mCherry vs. CNO-hM4Di post-test with Bonferroni correction: t_36_=1.10, ns) (Figure 3F). Together, these data suggest that reduced activity of the LS social engram, but not other neuronal populations in the LS, can promote social deficits in a sex-dependent manner.

**Figure 3.**
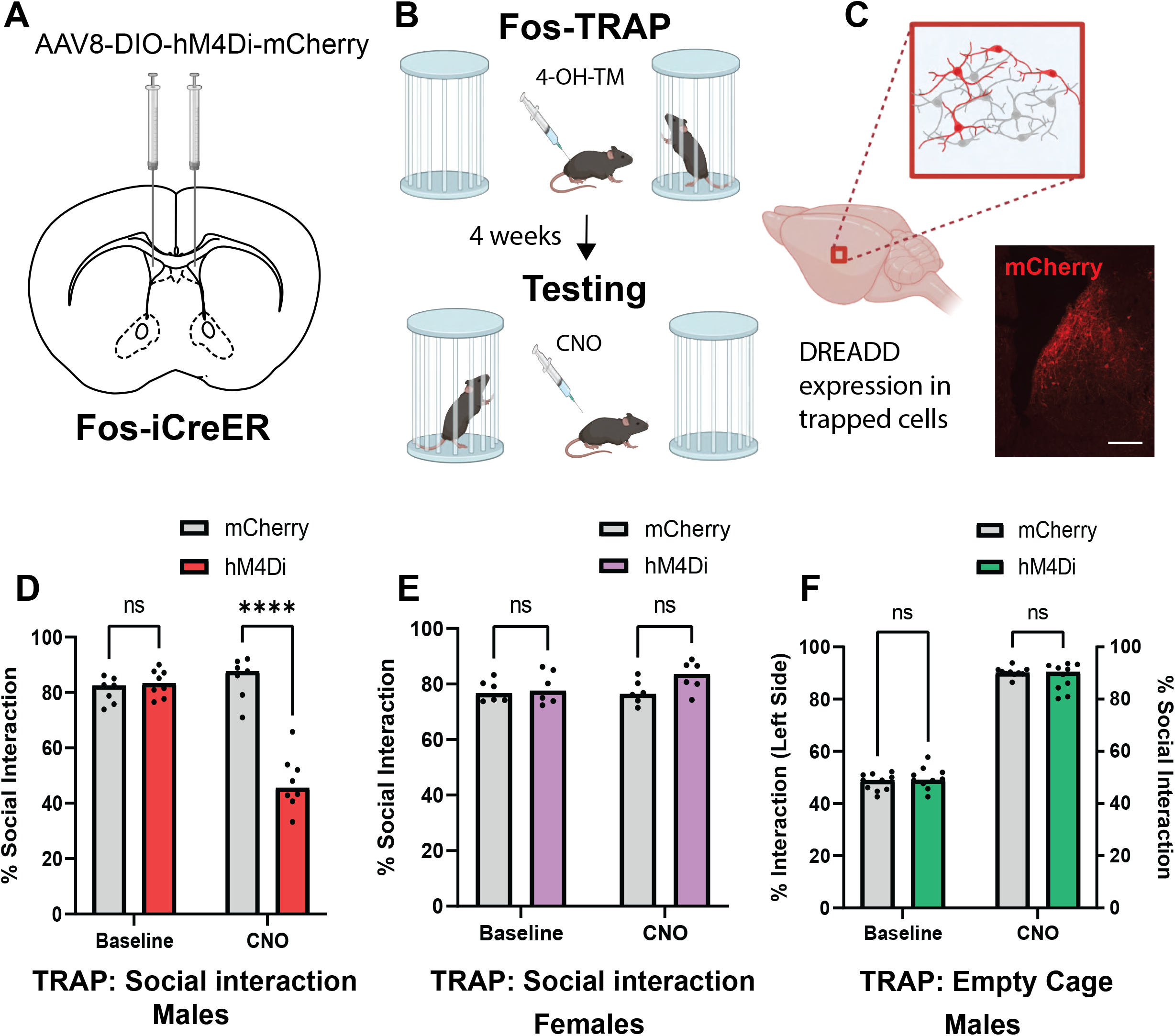
Chemogenetic inhibition of previously activated lateral septal neurons induces social deficits. (A) Experimental schematic of viral targeting of Gi-coupled DREADDs to the lateral septal neurons of Fos-iCreER mice. (B) Experimental timeline of Fos-TRAP (Targeted Recombination in transiently Active Populations) and behavioral testing followed by chemogenetic inhibition (via clozapine-N-oxide/CNO) of TRAPed lateral septal neurons. Fos- TRAP was facilitated by 4-hydroxytamoxifen administration 3 h after exposure to novel conspecifics, which led to selective expression of hM4Di in lateral septal neurons activated by social interaction in Fos-iCreER mice. (C) Illustration and a representative confocal image of hM4Di-mCherry expression in TRAPed lateral septal neurons (scale bar = 200 μm). (D) Histogram showing that chemogenetic inhibition of TRAPed lateral septal neurons led to social deficits in male Fos-iCreER mice. (E) In contrast, chemogenetic inhibition of TRAPed lateral septal neurons did not induce social deficits in female Fos-iCreER mice. (F) Chemogenetic inhibition of TRAPed lateral septal neurons previously activated by empty cage exposure did not affect social interaction behavior in male Fos-iCreER mice. Data are expressed as mean ± SEM. **** *p* < 0.0001. ns: non-significant. *Parts of this figure were made using Biorender*.

### Chronic ketamine dysregulates schizophrenia-associated gene networks in LS social engram neurons

Hypoactivation of the LS during social interaction suggests that chronic ketamine may fundamentally alter social engram neurons to render them less responsive to social stimuli. We next probed the entire translatome of LS social engram neurons to determine which genetic pathways were altered by ketamine treatment, and whether these pathways were shared with human schizophrenia and other mouse models of neurodevelopmental disorders.

Fos-iCreER mice were injected in the LS with AAV-ef1α-FLEX-EGFPL10a, and then administered 4-OH-TM after social interaction to induce expression of this EGFP-tagged ribosomal protein in LS social engram neurons. Fos-iCre::EGFPL10a^LS^ mice then received ketamine or saline injections for 10 consecutive days. Brains were extracted for translating ribosome affinity purification (ribo-TRAP) to isolate RNA that is being actively translated in the LS[38]. EGFP expression (Ct value = 9) was verified by qRT-PCR, confirming successful trapping of c-Fos-positive neurons and expression of the FLEX-EGFPL10a construct in the activated neuronal population. After validating the results from the RT-qPCR experiments, the samples were processed for RNA-seq following established protocols. Differential gene expression analysis was carried out using DESeq2 by setting the default false discovery rate (FDR) adjusted *p*-value < 0.05 and absolute value of log2 fold change > 0.5 as cutoff criteria. The findings showed that 1962 out of 16,144 genes exhibited differential expression in the ketamine vs. saline TRAP fraction, among which the majority of differentially translating transcripts (1653) were upregulated in the ketamine group (Supplementary Table 1). However, a considerable number of genes (309) were downregulated as well (Supplementary Table 2). A volcano plot (Figure 4B), with the criteria of *p*-value < 0.0001 and absolute value of log2 fold change > 1.5, illustrated that the top DEGs after chronic ketamine exposure were primarily associated with neuroinflammation/immune response (*Ighv, Mx1, Nos2, Helz2, & Nlrp3*), risk of developmental disorders (*Itgb4, Hspg2, Rbm47, & Ebf3*), and extracellular matrix remodeling (*Col4a1, Flnb, & Hspg2*; see a list of notable dysregulated genes by category in Supplementary Table 3). These findings corroborate that chronic subanesthetic ketamine is a valid model of schizophrenia, which has been associated with neuroinflammation and neuroimmune response[39–41], as well as extracellular matrix alterations[42]. We also generated a heatmap for hierarchical clustering of the top 50 DEGs (*p*-value < 0.0000001, and absolute value of log2 fold change > 0.5; Figure 4C), which not only include the aforementioned three categories of DEGs, but also contain a group of dysregulated genes related to myelination or oligodendrocyte function: *Plp1, Flnb, Myrf, Mag, Ermn*, etc., which resonates with various lines of evidence implicating myelin and oligodendrocyte dysfunction in schizophrenia[43, 44]. Moreover, among the top 50 DEGs, the epigenetic reader and modifier genes, *Prrc2c* (m6A reader) and *Jmjd1c* (H3K9 demethylases), were upregulated, which could lead to chromatin opening and drive the upregulation of a significant number of genes. In addition, we noticed that a long non-coding RNA (IncRNA) gene, *Meg3*, was upregulated. *Meg3* has various functions, notably in regulating social behavior[45], and has been suggested to serve as a biomarker in schizophrenia[46].

**Figure 4.**
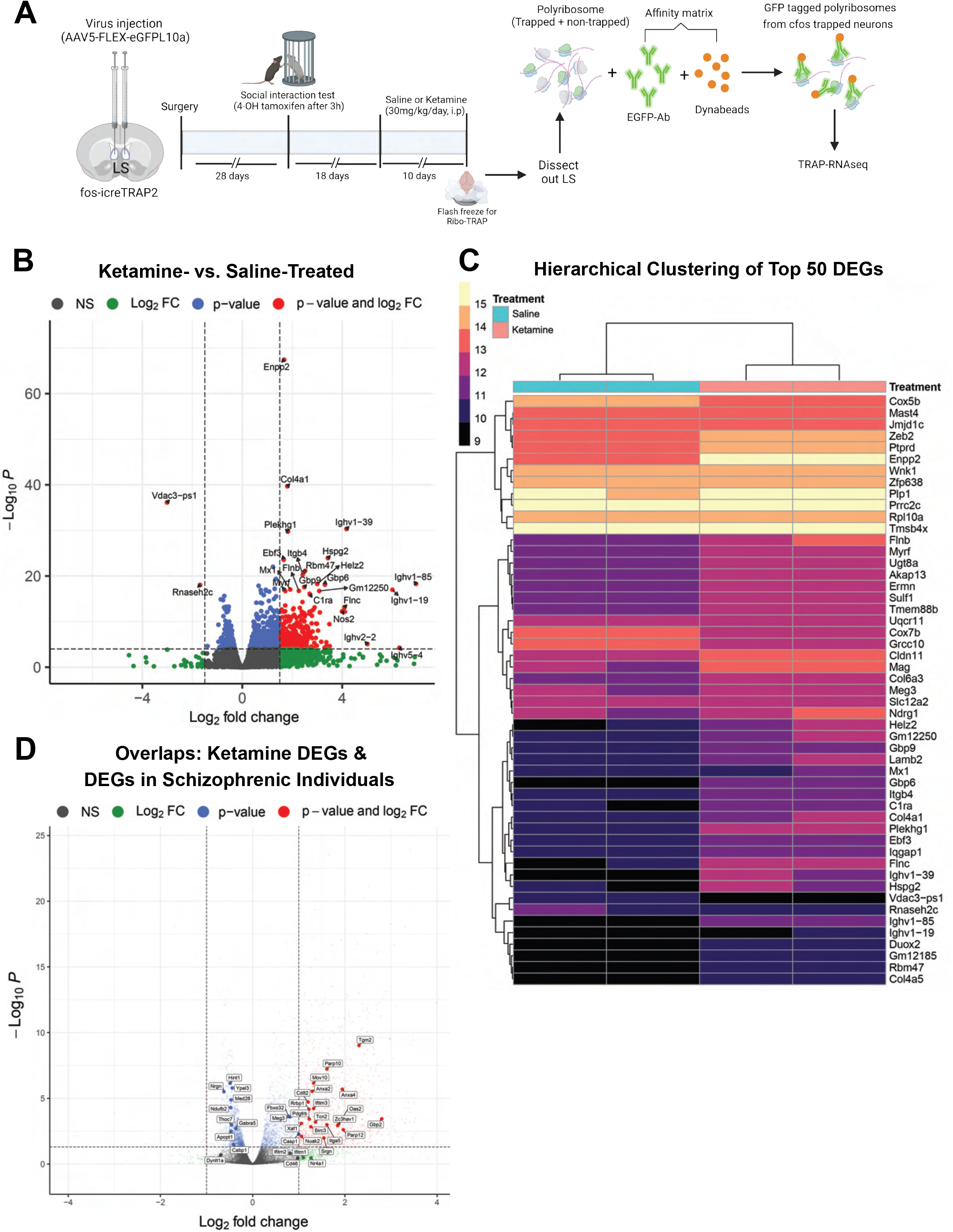
Chronic ketamine-induced transcriptomic changes in lateral septal neurons activated by social interaction. (A) A schematic of the experimental timeline and the procedure of translating ribosome affinity purification (ribo-TRAP) used to reveal transcriptomic changes induced by chronic ketamine in lateral septal (LS) neurons activated by social interaction in Fos-iCre mice. (B) Volcano plot showing chronic ketamine-, versus saline-, induced transcriptional changes in social interaction-activated LS neurons, with false discovery rate (FDR) < 0.05, *p* < 0.0001, and absolute value of log2 fold change > 1.5. Upregulated genes are labeled in red, downregulated in blue, and non-significantly changed in grey. The 20 most differentially expressed genes (DEGs) are named in the plot. (C) Heatmap for hierarchical clustering of top 50 DEGs. Criteria for DEG inclusion: *p* < 0.000001, and absolute value of log2 fold change > 0.5. Two samples within each group are illustrated in two separate columns. Color scheme depicts the mRNA expression levels (lower at 9 and higher at 15). (D) Volcano plot showing the overlaps between chronic ketamine DEGs in the present study and DEGs in individuals with schizophrenia from a meta-analysis by Merikangas et al. (2022). Criteria for DEG inclusion: p < 0.05, or absolute value of log2 fold change > 0.5. *Parts of this figure were made using Biorender*.

We then identified 38 DEGs from the present study that have associations with schizophrenic individuals based on a meta-analysis highlighting 160 crucial genes in schizophrenia pathophysiology [46] (Supplementary Table 4), illustrated in a volcano plot, with *p*-value < 0.05 or absolute value of log2 fold change > 0.5 (Figure 4D). Not surprisingly, among upregulated genes are those involved in neuroinflammation/immune response, i.e., *Nr4a1, Casp1, Anxa4*, and *Anxa2*. Among downregulated genes, some are notably involved in mitochondrial function, such as *Ndufb2* and *Vdac3*. Likewise, in our dataset, several genes encoding ATP synthase subunits (e.g., *Atp5e, Atp5g1, Atp5l*, and *Atp5j*) were downregulated. This is consistent with reports that mitochondrial dysfunction is linked to neuroinflammation underlying schizophrenia pathogenesis[47, 48]. Moreover, among the 38 differentially expressed genes shared by schizophrenic individuals and ketamine-treated mice, some are implicated in apoptotic processes, including *Birc3*, *Ifitm3*, *Xaf1*, and *Anxa2*, which corroborates that apoptosis contributes to schizophrenia pathophysiology[49].

It is noted that four genes encoding potassium inwardly-rectifying (Kir) channels, *Kcnj10*, *Kcnj13, Kcnk6*, and *Kcnk13*, were upregulated. Kir channels have been shown to tightly regulate neuronal excitability[50, 51]. Upregulation of Kir channels could contribute to the reduced LS neuronal activation after chronic ketamine exposure. Regarding neuronal excitability, we also observed upregulation of a few solute transporters, including *Slc44a1* (choline transporter), *Slc4a4* (involved in excitatory synaptic transmission), *Slc7a11* (cysteine/glutamate transporter), and *Slc12a2* (sodium/chloride transporter), possibly affecting neuronal signaling pathways.

In addition, we conducted comprehensive network analyses of the translatome, uncovering noteworthy connections between our ketamine model and other models of neurodevelopmental disorders, including prenatal exposure to valproic acid and material immune activation with Polyinosinic-polycytidylic acid or Poly(I:C). Specifically, we have observed that out of 1660 genes linked to valproic acid exposure, 245 DEGs are associated with our ketamine model (FDR-adjusted *p* = 9.132e-6); out of 26 genes linked to Poly(I:C) exposure, 15 DEGs are associated with our ketamine model (FDR-adjusted *p* = 1.680e-7) (Supplementary Figure 2). These findings further validate the use of our chronic subanesthetic ketamine paradigm for studying schizophrenia.

We then asked whether social engram neurons express unique genetic markers that might distinguish them from the population at large. Among the 50 most abundant genes in social engram neurons in saline-injected mice were *Gad1* and *Gad2* (glutamic acid decarboxylase 1 and 2), which are indicative of GABAergic neurons. This is expected based on previous reports that the LS is enriched in GABAergic neurons[52, 53]. We also found a high abundance of *Camk2a* (calcium/calmodulin dependent protein kinase II alpha) in these neurons, which was unexpected as most LS GABAergic neurons express other calcium binding proteins including calbindin and calretinin[54]. However, while CaMKII is in relatively low abundance in GABAergic interneurons, it may be highly expressed in GABAergic projection neurons[55]. This suggests that GABAergic projection neurons may be enriched in the LS social engram population (Supplementary Table 5).

### Mapping downstream targets that are activated by LS social engram neurons with iDISCO

The LS has extensive projections to parts of the limbic forebrain, thalamus, and pontine central gray[56], making it ideally positioned to orchestrate complex social behavior. We asked whether LS social engram neurons could activate downstream targets and whether these were direct post-synaptic targets of the LS. Fos-iCreER::hM3Dq^LS^ or Fos-iCreER::mCherry^LS^ mice were trapped after social interaction and received CNO injections 4 weeks later to reactivate the LS social engram, followed by brain extraction for c-Fos iDISCO. Here we found that chemogenetic re-activation of LS social engram neurons enhanced neural activity in the dentate gyrus (DG), olfactory areas, intercalated nucleus of the amygdala (IA), subparafascicular thalamic nucleus (SPF), and the ventral part of the basolateral amygdala (BLA) (Figure 5). We then used AAV-mediated transsynaptic anterograde tracing to map brain regions that receive direct inputs from the LS. EGFP-L10a mice were injected with AAV1-hSyn-Cre-hGH in the LS, which drives Cre-dependent expression of transgenes (i.e. L10A-eGFP) in direct post-synaptic neuronal targets[57]. Here we found that the LS had direct inputs to a large proportion (>50%) of neurons in the medial septum, infralimbic prefrontal cortex, and olfactory areas. A high percentage of neurons (30-50%) were also found in the diagonal band of the Broca area (dbB) and the hippocampus (CA1 and CA3), while a smaller proportion (20-30%) were found in the lateral hypothalamus, locus coeruleus, preoptic area of the hypothalamus, and the anterior cingulate cortex. Between 10-20% of neurons in the prelimbic cortex, medial thalamus, orbitofrontal cortex, and dorsal raphe nucleus (DRN) had LS inputs, while the smallest fraction (<10%) was found in the ventral tegmental area, insular cortex, somatosensory cortex, bed nucleus of the stria terminalis, nucleus accumbens, motor cortex, basolateral amygdala (BLA) and laterodorsal tegmental nucleus (Supplementary Figure 3). Of the areas that receive direct efferent projections from the LS, the olfactory area, thalamus and BLA were also activated by LS social engram neurons, suggesting that other regions like the DG may be indirectly modulated by these neurons. It was also interesting that most regions receiving direct input from the LS did not show enhanced activity by chemogenetic re-activation of LS social engram neurons. This may be explained by the fact that LS neurons are primarily GABAergic[52, 53], so these areas may be inhibited rather than activated by chemogenetic stimulation of the LS social engram.

**Figure 5.**
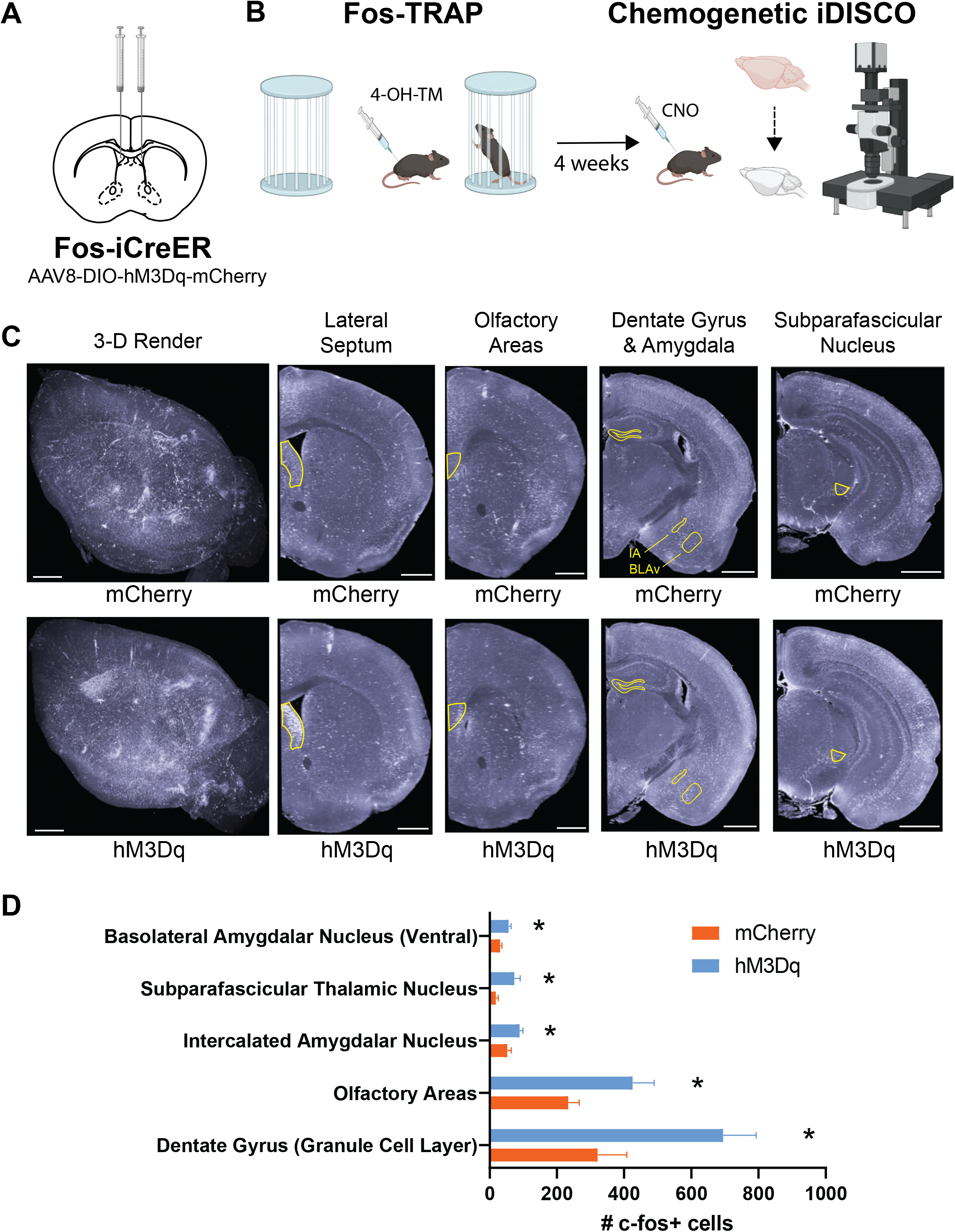
Mapping downstream targets that are activated by LS social engram neurons with iDISCO. (A) Experimental schematic of viral targeting of Gq-coupled DREADDs to the lateral septal neurons of Fos-iCreER mice. (B) Experimental timeline of Fos-TRAP and chemogenetic iDISCO. Fos-TRAP was facilitated by 4-hydroxytamoxifen administration 3 h after exposure to novel conspecifics, leading to selective expression of hM3Dq in lateral septal neurons activated by social interaction in Fos-iCreER mice. 4 weeks later, 2.25 h following chemogenetic activation of TRAPed lateral septal neurons (via clozapine-N-oxide/CNO), mouse brains were fixed, cleared, and imaged using iDISCO. (C) 3D rendering of c-fos expression in representative iDISCO brains from the mCherry and hM3Dq groups (scale bars = 1,000 μm), as well as coronal views through the lateral septum, olfactory areas, dentate gyrus (granule cell layer), intercalated amygdalar nucleus (IA), basolateral amygdalar nucleus (ventral, BLAv), and subparafascicular thalamic nucleus (scale bars = 1,000 μm). (D) Histogram of c-fos-positive cell counts in select brain regions. Data are expressed as mean ± SEM. * *p* < 0.05. *Parts of this figure were made using Biorender*.

The main limitations of the chemogenetic iDISCO approach is that c-Fos is mainly a marker of neuronal activation, so the inhibitory actions of the LS may go undetected, especially in regions of low basal c-Fos activity. We next conducted a small, exploratory study of six animals pairing chemogenetic activation of the LS social engram with functional magnetic resonance imaging (cg-fMRI) to assess whether the large-scale activity patterns emerging from LS social engram activity aligned with our iDISCO results. Four weeks after the Fos-TRAP procedure following social interaction, Fos-iCreER::hM3Dq^LS^ mice underwent baseline fMRI scans on an Agilent Discovery 901 7-Tesla small animal scanner under isoflurane anesthesia. One week later, they were injected with CNO and re-scanned under the same conditions. Seed analysis performed between the LS and all voxels in the rest of the brain before and after injection of CNO showed a significantly decreased connectivity between the LS and the right caudate-putamen as well as the right piriform cortex (TFCE corrected p<0.05) following chemogenetic reactivation of the LS social engram (Figure 6). Reduced connectivity of the LS with the piriform and caudate-putamen may underlie some aspects of social behavior, including behavioral flexibility, reward processing[58], and the processing of olfactory cues.

**Figure 6.**
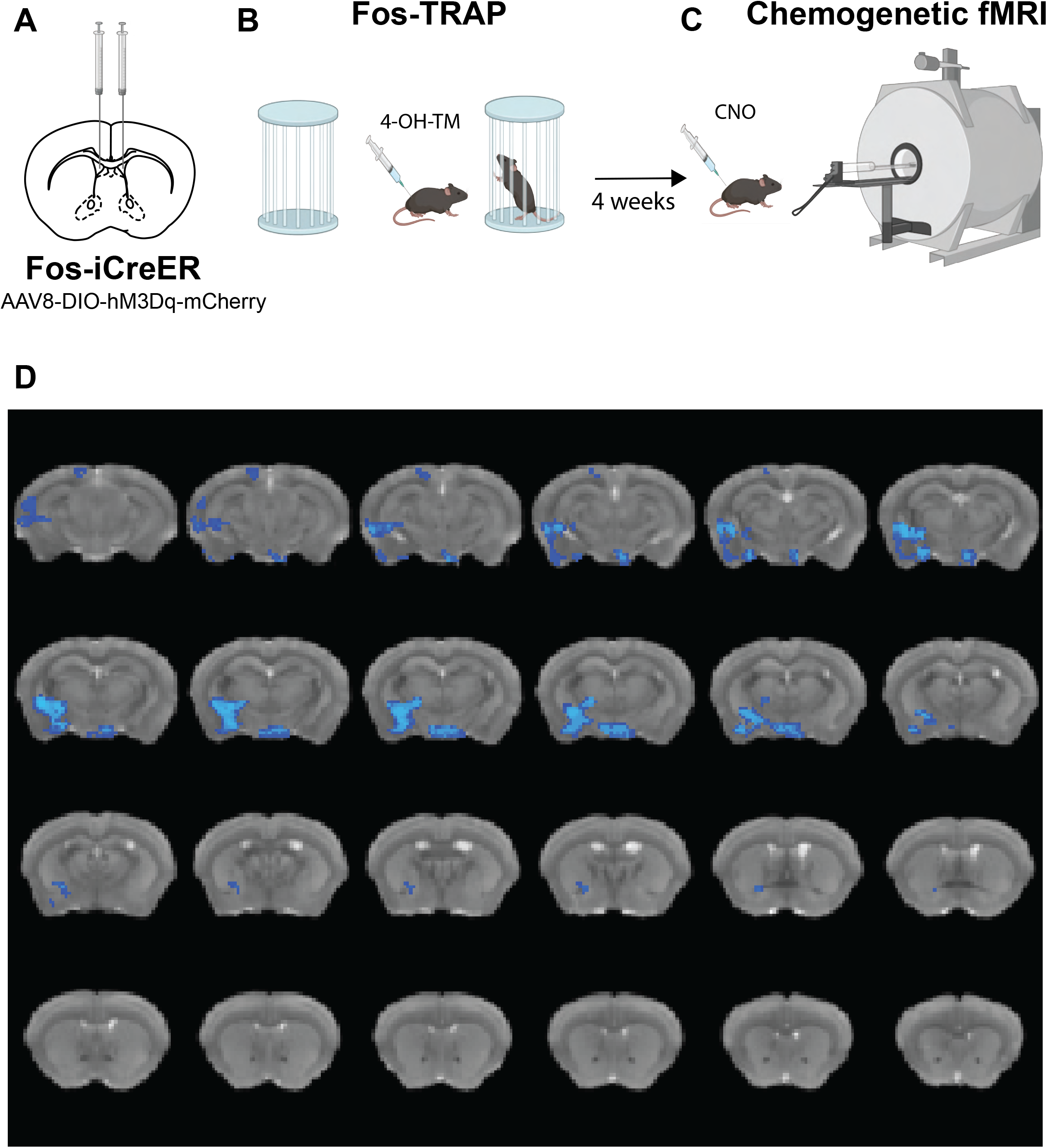
Chemogenetic fMRI mapping of large-scale activity patterns regulated by LS social engram neurons. (A) Experimental schematic of viral targeting of Gq-coupled DREADDs to the lateral septal neurons of Fos-iCreER mice. (B) Experimental timeline of Fos- TRAP and (C) chemogenetic fMRI following CNO injection. (D) Image showing the results for the second level analysis of voxel correlation for the LS seed in blue, where correlation at baseline > correlation following CNO injection, with the p-value set to p<0.05. Underlying image is the Waxholm template that all data are registered to. Each slice is spaced 200 µm apart. *Figure was generated Biorender and FSLeyes*.

### Viral tracing of afferent inputs to the LS that are activated by social stimuli

The chemogenetic iDISCO and fMRI can pinpoint regions that receive efferent input from the LS or one of its projection targets. However, the LS also receives direct afferent projections from the amygdala, hippocampus, thalamus, midbrain and hindbrain which may alter excitability of the LS social engram. To label cellular networks that project to the LS and are responsive to social stimuli, Fos-iCreER mice were injected with a retrograde AAV expressing Cre-dependent tdtomato (AAVrg-hsyn-FLEX-tdtomato) in the LS, then trapped after social interaction or exposure to two empty cages. Here we found that social interaction significantly elevated the percent immunoreactive area in the PCG, the hypothalamus, and dbB (Supplementary Figure 4). The PCG has distinct GABAergic and glutamatergic populations that process the positive and negative valence of sensory stimuli, respectively[59], which may be relevant in orchestrating social approach behaviors. Interestingly, our iDISCO data indicated that activity in the PCG was reduced after social interaction, but this included all neurons in the PCG, whereas here we are considering only those PCG neurons that project to the LS. The PCG consists of distinct populations of neurons that have opposing roles in valence processing, with glutamatergic neurons being activated by aversive stimuli and GABAergic neurons being activated by rewarding stimuli. It would be interesting in a future study to determine which of these two populations is activated during social interaction and whether this activity is modified by ketamine treatment.

## Discussion

Social withdrawal is a common behavioral endpoint for schizophrenia and other neurodevelopmental disorders, but there is limited information regarding how social behavior goes awry in these disorders. The present study advances our knowledge in this area by providing unbiased insight into specific regions that are dysregulated during social behavior in a ketamine mouse model with iDISCO. Using this approach, we show that ketamine results in hypoactivation of the LS during social behavior, which aligns well with the current literature including a recent study in a mouse model of social defeat [33]. We then demonstrate that this hypoactivity in the LS is a primary driver of schizophrenia-like social deficits. Using Fos-iCreER mice to parse out LS neurons that respond to social stimuli (i.e. the LS social engram), we then show that inhibition of specific neuronal ensembles in the LS reproduced social deficits in ketamine-naïve male mice, but not in females, suggesting that the LS may be differentially involved in schizophrenia-like social behavior. This supports the observation in humans that males with schizophrenia are affected earlier and more severely than females and have a higher incidence of negative symptoms and social impairment[37]. Interestingly, oxytocin receptor (Oxtr)-expressing neurons in the LS have been linked to deficits in social interaction in both male and female mice[34, 60]. However, an important distinction here is that Oxtr activation in the LS restores social novelty in male mice treated with valproic acid, whereas in female mice, Oxtrs attenuates conditioned social fear. Valproic acid enhances GABAergic transmission and has been used to model autism in mice[61], so the neural processes disrupted by valproic acid may be similar to those disrupted by ketamine. Another significant finding is that chemogenetic inhibition of neurons that respond to non-social stimuli (e.g. two empty cages) does not induce social deficits, suggesting that social behavior is likely driven by LS social engram neurons and not the LS population at large.

Hypoactivation of the LS social engram may promote social deficits in schizophrenia, but how does ketamine treatment alter the intrinsic excitability of these neurons? By using ribo-TRAP to probe the entire translatome of this social engram cellular network, we observe upregulation of genes encoding potassium inwardly-rectifying channels and solute transporters as well as dysregulation of genes implicated in apoptotic processes, all of which could contribute to the hypoactivation of LS social engram neurons after chronic ketamine exposure. Interestingly, genes involved in extracellular matrix remodeling are among the top upregulated DEGs. It is known that LS neurons are predominantly GABAergic, and extracellular matrix plays a significant role in regulating GABAergic neuronal activity[62, 63]. As such, extracellular matrix alterations are possibly implicated in ketamine-induced LS hypoactivation as well. Extracellular matrix impairments have been observed in several brain regions in postmortem schizophrenic patients as well as rodent models of schizophrenia[64], which warrant further investigations of such abnormalities in schizophrenia pathophysiology. Certain genes regulating neurodevelopment (*Col4a1, Hspg2, Rbm47, Ebf3, Zeb2,* and *Tgm2*) have also been upregulated in the LS social engram neurons, and these genes are closely linked to developmental disorders/schizophrenia [65–70]. In addition, chronic ketamine induces upregulation of neuroinflammation-related genes and downregulation of genes involved in mitochondrial function, which is consistent with observations that schizophrenia is associated with mitochondrial dysfunction and immune system disturbances[71], Notably, positive, negative, and cognitive symptoms of schizophrenia are all linked to inflammation [72].

Moreover, we have identified a group of dysregulated genes associated with myelination or oligodendrocyte function, impairment of which has been implicated in schizophrenia pathophysiology[73]. These congruent findings provide insights into potential schizophrenia pathogenesis and intervention strategies.

In addition to these intrinsic alterations in cellular/molecular function, ketamine may also promote hypoactivity in afferent projections to LS social engram neurons, which would be expected to alter their excitability. We observe that the LS-projecting neurons in the dbB, PCG and hypothalamus are significantly activated by social stimuli in Fos-iCreER::AAVrg-flex- tdTomato^LS^ mice. Interestingly, our iDISCO results indicate that the PCG is inactivated by social stimuli, suggesting that LS-projecting PCG neurons may have a distinct role in valence processing of social stimuli that differs from the PCG population at large. If social stimuli are rewarding, these LS-projecting PCG neurons may represent GABAergic neurons that are activated by rewarding stimuli[59]. Further studies will be required to resolve the genetic identity of these LS-projecting PCG neurons and whether they are modulated by ketamine treatment.

The LS social engram also has a wide range of efferent projections throughout the brain that may participate in orchestrating social behavior. Here we show that chemogenetic activation of the LS social engram promotes c-Fos activity in downstream regions including the DG, olfactory areas, IA, SPF, and the ventral part of the BLA, each of which may be a crucial part of the social circuit that is dysregulated by ketamine. Neurogenesis and expansion of newborn granule neurons in the DG has been found to improve social recognition in a mouse model of autism[74], while inhibiting GABAergic projections from the medial septum to the DG relieves depressive-like behaviors in mice after chronic social defeat[75]. Together, these studies suggest that activity in the DG is important for social motivation and social recognition which may be attenuated by chronic ketamine and the resultant hypoactivation of the LS social engram. Since mice rely heavily on olfaction to navigate their environment, olfactory areas are expected to play an important role in social recognition. The IA consists of clusters of GABAergic neurons surrounding that basolateral nuclear complex (BNC) of the amygdala which are thought to mediate fear extinction by inhibiting amygdalar outputs to the hypothalamic and brainstem circuits[76–78]. These IA neurons receive excitatory inputs from the infralimbic cortex (IL) and BNC that presumably activate this fear extinction circuit[79]. It can be argued that activation of this fear extinction circuit by the LS social engram is necessary for social behavior to occur, and that without it, avoidance of novel social partners will predominate. The SPF projects to somatosensory areas and is part of the posterior intralaminar thalamic nucleus, which can promote social grooming via its projections to the medial preoptic area[80, 81].

Dopaminergic projections from the SPF to the inferior colliculus may also attenuate neuronal responsiveness to unexpected sensory input[82]. In addition, the BLA seems to modulate social behavior in an input-specific manner, with IL➔BLA projections promoting social interaction while prelimbic (PL)➔BLA projections impair social behavior[83]. The IL➔BLA pathway may function similarly to the IL➔ICN pathway by inhibiting social aversion through the fear extinction circuit. Together, these studies provide converging evidence that social behavior may be articulated by efferent projections of LS social engram to these downstream regions. Although the DRN is not significantly activated by the LS social engram, possibly due to c-Fos being a weak marker of neural activity in the brainstem, it is an important hub for coordinating activity between different regions that have been implicated in social behavior, including the LS, habenula, thalamus, hippocampus, cerebellum and olfactory bulb[84]. Importantly, we found that the DRN receives significant inputs from the LS, so reduced activity in the LS after ketamine may dysregulate downstream activity in these interconnected regions via the DRN.

The results of our exploratory cg-fMRI study suggest that there is a significantly reduced connectivity between the LS and the piriform cortex as well as the caudate-putamen after chemogenetic stimulation of the LS social engram. The LS is primarily composed of GABAergic neurons, so chemogenetic activation of these neurons may reduce functional connectivity of the LS with brain regions like the piriform and caudate-putamen that receive these GABAergic inputs. While we cannot say for certain how LS inputs to the piriform shapes social behavior, the piriform cortex has been found to be hyperexcitable following social defeat, which is known to cause deficits in social interaction as well as learning and memory impairments[85]. Other studies suggest that downregulation of Oxtrs in the piriform cortex may contribute to social deficits in reelin haploinsufficient (+/-) mice, which have been used to model neurodevelopmental disorders including autism and schizophrenia[86]. It is also interesting that LS social engram activity may reduce LS connectivity with the caudate-putamen. The LS and the caudate-putamen are activated by atypical antipsychotics, but activation of the LS is thought to alleviate positive symptoms while activity in the caudate-putamen is associated with the motor side effect liability of these medications[87, 88]. Our results suggest that increasing activity in discrete neuronal circuits in the LS can help to overcome social deficits, which may alleviate negative symptoms and simultaneously reduce locomotor side effects of antipsychotic drugs by inhibiting the caudate-putamen. The LS may represent an important pharmacological target for schizophrenia drug development that improves both positive and negative symptoms while minimizing side effect liability. Future studies will focus on deciphering how the LS interacts with these downstream neuronal networks to promote schizophrenia-like social deficits induced by chronic ketamine, providing further insight into potential treatments.

## Acknowledgements

We thank Gabrielle Bierlein De-La-Rosa, Keland Moore, Hannah Stutt, and Emily Herum for excellent technical assistance with iDISCO, light sheet imaging, confocal imaging, stereotaxic injections, behavioral testing in mice, and RNA-seq data analysis. We additionally thank Dr. Shane Heiney at the Iowa Neuroscience Institute Neural Circuits and Behavior Core for technical assistance and training with the La Vision Ultramicroscope II (funded by the Roy J. and Lucille A. Carver Charitable Trust) and the Iowa Institute for Biomedical Imaging for technical assistance with fMRI imaging. RNA-seq data presented herein were generated by the Genomics Division of the Iowa Institute of Human Genetics which is supported, in part, by the University of Iowa Carver College of Medicine.

## Declarations

### Ethics approval

All procedures on mice in this study were approved by the Institutional Care and Use Committee at the University of Iowa.

### Availability of data and materials

All data generated or analyzed during this study are included in this published article and its supplementary information files. TRAP-seq data have been deposited in the NCBI’s GEO database (GSE241234).

---

**Table 1:** Viral constructs.

| **AAV** | **Virus Source** | **Cat No.** | **Titer** |
| --- | --- | --- | --- |
| AAV8-hSyn-mCherry | Addgene | 114472-AAV8 | 1×10<sup>13</sup> vg/ml |
| AAV8-hSyn-hM3D(Gq)-mCherry | Addgene | 50474-AAV8 | 2×10<sup>12</sup> vg/ml |
| AAV8-hSyn-DIO-mCherry | Addgene | 50459-AAV8 | 1 x 10<sup>13</sup> vg/ml |
| AAV8-hSyn-DIO-hM3D(Gq)-mCherry | Addgene | 44361-AAV8 | 4 x 10<sup>12</sup> vg/ml |
| AAV8-hSyn-DIO-hM4D(Gi)-mCherry | Addgene | 44362-AAV8 | 1 x 10<sup>13</sup> vg/ml |
| AAVrg-FLEX-tdTomato | Addgene | 28306-AAVrg | 7 x 10<sup>12</sup> vg/ml |
| AAV5-FLEX-EGFPL10a | Addgene | 98747-AAV5 | 7×10<sup>12</sup> vg/ml |
| AAV1-hSyn-Cre-WPRE-hGH | Addgene | 105553-AAV1 | 7×10<sup>12</sup> vg/ml |

**Table 2:** Antibodies used in immunofluorescence.

| **Stain** | **Antibody** | **Concentration** |
| --- | --- | --- |
| *RFP* | Rabbit anti-RFP<br>(Rockland Immunochemicals; Catalog # 600-401-379) | 1:500 |
|  | Alexa Fluor 555 donkey anti-rabbit<br>(Invitrogen; Catalog # A-31572) | 1:1,000 |
| *NeuN* | Guinea pig anti-NeuN<br>(MilliporeSigma; Catalog # ABN90) | 1:2,000 |
|  | Cy3 donkey anti-guinea pig<br>(Jackson ImmunoResearch; Catalog # 706-165-148) | 1:500 |

