## Supplementary Methods and Figures for "Lateral Septal Circuits Govern Schizophrenia-Like Effects of Ketamine on Social Behavior"

**This PDF file includes:**

**Supplementary Methods**

**Supplementary Figures and Legends S1-S4**

**References**

### Materials and Methods

#### Animals

Adult C57BL/6J mice (strain: 000664) from Jackson Labs (Bar Harbor, ME, USA) and in-house bred $Fos^{2A\text{-}iCreER}$ (TRAP2) (Jackson Labs, strain: 030323) as well as EGFP-L10a (Jackson Labs, strain: 024750) mice were used in the experiments. All mice were kept at a temperature- and humidity-controlled vivarium at the University of Iowa, with *ad libitum* access to food and water, under a 12h/12h light/dark cycle, in conformity with the guidance from AAALAC. All the experimental procedures were approved by the Institutional Animal Care and Use Committee at the University of Iowa.

#### Drugs

In chemogenetic experiments, Clozapine-N-oxide dihydrochloride (CNO; Hello Bio, Princeton, NJ, USA) was dissolved in 0.9% sterile saline at 0.3 mg/ml. Mice were injected at the volume of 10 ml/kg for a dose of 3 mg/kg. In TRAP experiments, 4-hydroxytamoxifen (4-OH-TM; Catalog No. H6278; MilliporeSigma, Burlington, MA, USA) was first dissolved in dimethyl sulfoxide (DMSO) at 40 mg/ml, which was then diluted with saline containing 2% Tween 80 to obtain a 2 mg/ml solution, After a further dilution with saline at 1:1, the final concentration of 4-hydroxytamoxifen was 1 mg/ml. Mice were injected at the volume of 10 ml/kg for a dose of 10 mg/kg.

#### Chronic subanesthetic ketamine administration

Ketamine was administered intraperitoneally at the dose of 30 mg/kg/day for 10 consecutive days. Specifically, solid ketamine hydrochloride (Catalog No. K2753; MilliporeSigma) was dissolved in saline at 3 mg/ml. Mice were injected at the volume of 10 ml/kg for the dose of 30 mg/kg daily between 9:00 and 11:00 AM. The drug was administered in a quiet procedural room, which the mice got acclimated to for one hour before injections. Hyperlocomotion and disorientation were observed in mice shortly after the ketamine administration. The control mice were injected with saline of the same volume. The subanesthetic dose of 30 mg/kg was chosen in consistency with previous studies in rodents [1, 2].

### Stereotaxic surgeries

We performed stereotaxic surgeries for intracranial AAV infusions in the lateral septum/LS (see Table 1 for viral constructs we used). Specifically, under anesthesia with isoflurane (3% v/v induction and 1.5 – 2% maintenance), the mouse was placed in an Angle Two stereotaxic frame (Leica Biosystems, Wetzlar, Germany) on a heating pad. A 1 μl Neuros syringe needle (Model: 7001 KH; Hamilton Company, Reno, NV, USA) was positioned over the LS (ML = ±0.35, AP = +0.85, DV = -3.20, θ = 0°, relative to the Bregma) and the AAV was infused with a Stoelting Quintessential Stereotaxic Injector (Wood Dale, IL, USA) at a rate of 100 nl/min for the total volume of 250 or 300 nl/side. The needle was kept in place for 10 more min to allow for diffusion of the virus before being lifted. The mouse was administered with meloxicam (4 mg/kg/day, s.c.) for 2 – 3 days to relieve post-operative pain and inflammation. Experiments were conducted at least 4 weeks after the surgeries.

### Behavioral tests

#### *Social interaction test*

The social interaction test has been described previously [3, 4]. Briefly, it was performed in a dimly lit (~20 lux in the center) three-chambered Plexiglas container (L × W × H: 64 × 42 × 21 cm) divided by 2 panels with small openings at the bottom for free movement of mice between chambers. Each session started with the test mouse being placed in the center chamber, who was allowed to move freely throughout the 3 chambers for 10 min. The mouse was then briefly confined in the center chamber, when a small holding cage (i.e., a 11-cm-high wire mesh pen cup upside down, open end: 10 cm Ø; closed end: 8 cm Ø) was placed in each outmost chamber, and a conspecific of the same age and sex, strange to the test mouse, was placed inside one holding cage, with the other holding cage remaining empty. The placement of the stranger mouse was alternated between testing sessions to control for side preferences. Afterwards, the openings between chambers were unblocked and the test mouse was allowed to travel freely between chambers for 10 more min. Animal behavior was recorded by an overhead digital camera with Media Recorder software (Noldus, Leesburg, VA, USA). The time spent by the test mouse interacting with the novel conspecific and exploring the empty cage was scored by an experimenter blinded to the experimental conditions.

***Open field test***

The open field test was performed in a dimly lit (~20 lux in the center) opaque Plexiglas arena (L x W x H: 50 x 50 x 25 cm), situated in a custom-made sound-attenuating wood box. At the beginning of a session, the mouse was placed in a corner of the arena, who was allowed to freely roam and explore for 10 min. Animal activity was recorded with an overhead digital camera and analyzed with Ethovision XT14 software (Noldus). The following parameters were scored: latency to the center area, time spent in the center area, and total distance moved. The center area was defined as the central 15% of the arena.

***Prepulse inhibition (PPI) test***

The prepulse inhibition test based on acoustic startle has been described previously [5, 6]. In brief, the test was performed in a small Plexiglas cylinder inside a sound-attenuating box. The cylinder was placed on a piezoelectric transducer to quantify vibrations and transmit the information to a computer. There is a loudspeaker inside the box to deliver acoustic stimuli, and the background noise produced by a fan was kept at 70 dB. A testing session started with a mouse being placed in the cylinder. Following a 5-min habituation period, 42 trials of 7 different types were scheduled, including no-stimulus trials, trials with 40-ms 120 dB acoustic startle stimuli alone, and trials with prepulse stimuli (20 ms; 74, 78, 82, 86, or 90 dB) taking place 100 ms before the onset of the startle stimuli. Trials occurred in blocks of 7. Within each block, different trials were randomized with variable intertrial intervals (mean interval: 15 s: range: 10 – 20 s). For each trial, the startle amplitude was measured within a 65-ms sampling window. For each mouse, the magnitude of prepulse inhibition at each prepulse sound level is computed as follows.

$$prepulseinhibitionataprepulsesoundlevel(\%) = 100 - \frac{startleresponsewithprepulse}{startleresponsewithnoprepulse} \times 100$$

**Chemogenetic manipulations**

To facilitate chemogenetic manipulations, mice were transduced with AAVs to express Gq- or Gi-DREADDs in LS neurons, with controls expressing mCherry only. At least 4 weeks after the viral infusion surgeries, DREADDs were activated with CNO (3 mg/kg, i.p.) 45 min before the behavioral experiment or 2.25 h before perfusion and brain extraction for iDISCO. Dosage and timing of CNO was based on previous publications with this DREADD agonist [4, 7]. For experiments in C57BL/6J mice, DREADDs were expressed in any neurons that AAVs infected. In contrast, for experiments in Fos-iCreER mice, DREADDs were Cre-recombinase dependent, which could only be expressed in AAV-infected TRAPed neurons.

**Targeted recombination in transiently active populations (TRAP)**

To study neurons selectively activated by social interaction, we employed the method of targeted recombination in transiently active populations (TRAP) [8]. In brief, 3 h after initiation of interaction between a Fos-iCreER mouse and a novel conspecific, the Fos-iCreER mouse was administered with 4-OH-TM (10 mg/kg, i.p.), and kept undisturbed in the following several hours. The mouse home cages were changed 24 and 48 h after the 4-OH-TM administration to avoid recycling of tamoxifen due to coprophagia. As such, in the Fos-iCreER mouse, the tamoxifen-inducible Cre-recombinase was expressed from the locus of the immediate early gene *Fos* in a social interaction-dependent fashion, without disrupting endogenous *Fos* expression. Therefore,

TRAP could facilitate selective access to neurons activated solely by social interaction in our experiments.

### Translating ribosome affinity purification (ribo-TRAP)

#### *Ribo-TRAP*

To investigate the impact of ketamine on the translatome of LS-activated neurons during the social interaction test, we injected AAV5-FLEX-EGFPL10a (Addgene 98747-AAV5) into the LS of Fos-iCreER mice to label the social engram neurons by EGFP-tagged L10a ribosomal subunits. To isolate translating mRNAs from these neurons, we performed translating ribosome affinity purification (ribo-TRAP) in fresh LS tissue (Figure 4A) following a previously described protocol [9, 10]. Briefly, after administering 10 days of ketamine/saline injections to the Fos-iCreER mice, the brains were rapidly frozen. The LS tissue was collected in an ice-cold dissection buffer (1X HBSS, 2.5 mM HEPES, 35 mM Glucose, 4 mM NaHCO3, and 100 μg/ml cyclohexamide), and since the expected number of activated neurons was relatively low, we pooled five LS tissue punches into one tube. Consequently, we processed 10 ketamine and 10 saline LS tissue punches to create 2 ketamine and 2 saline pooled samples, respectively, for ribo-TRAP. Next, we homogenized the pooled tissue in a tissue lysis buffer (20 mM HEPES, 10 mM $MgCl_2$, 150 mM KCl, 0.5 mM DTT, 100 μg/ml cyclohexamide, RNase inhibitors and EDTA-free protease inhibitors) using a prechilled micropestle, and the homogenate was centrifuged to obtain the supernatant containing polyribosomes. To pull down the EGFP-tagged polyribosomes, we used an affinity matrix containing Dynabeads (Thermo Fisher Scientific, Waltham, MA, USA), Biotinylated protein L (Genscript, Piscataway, NJ, USA), and GFP antibodies (HtzGFP-19F7 and HtzGFP-19C8, Heintz Lab, Rockefeller University, New York, NY, USA). The supernatant with EGFP-tagged polyribosomes was incubated with this affinity matrix, and after incubation, the beads (now with EGFP-tagged polyribosomes) were washed in high salt buffer (20 mM HEPES, 10 mM MgCl2, 350 mM KCl, 1% NP-40, 0.5 mM DTT and 100 μg/ml cyclohexamide). RNA from the complex was isolated using the Agilent Absolutely RNA Nanoprep kit (Agilent, Santa Clara, CA, USA) following the manufacturer's instructions. Further, before sequencing the RNA was quantified in Drop Sense (Perkin Elmer, MA, USA) and Qubit HS RNA (Thermo Fisher Scientific) and checked for quality in 2100 Bioanalyzer (Agilent).

#### *RNA sequencing*

We used total RNA with a RNA Integrity Number (RIN) greater than 7 to prepare sequencing libraries from 50 ng TRAP samples of saline (n = 2 samples) and ketamine (n = 2 samples) at the Iowa Institute of Human Genetics (IIHG), Genomics Division. The Illumina Stranded Total RNA Prep, Ligation with Ribo-Zero Plus sample preparation kit (Illumina, San Diego, CA, USA) was used for library preparation. Library concentrations were measured, and the average size was determined using the Qubit dsDNA HS kit (Thermo Fisher Scientific) and the 2100 Bioanalyzer (Agilent). The four libraries were then combined equally based on molar concentrations, and the pool's concentration was measured using the KAPA Illumina Library Quantification Kit (KAPA Biosystems, Wilmington, MA, USA). Finally, the pool was sequenced on the Illumina NovaSeq 6000 sequencer with 100bp Paired-End SBS (Sequencing by Synthesis) chemistry at the IIHG Core Facility.

#### *TRAP-seq analysis*

Barcoded samples were pooled and sequenced, to obtain an average of 223 million paired-end 100 bp reads per sample. Reads were converted from the native Illumina BCL format to fastq format with ‘bcl2fastq’ (v2.20.0.422). Data was processed using nf-core/rnaseq v3.10 (doi: https://doi.org/10.5281/zenodo.1400710) of the nf-core collection of workflows [11].

The pipeline was executed with Nextflow v22.10.1 [12] on the University of Iowa's Argon HPC cluster with the following command:

```
nextflow run nf-core/rnaseq -r 3.10 -profile argon --igenomes_base
/Users/mchiment/igenomes/references --input samplesheet.csv --outdir nf --genome GRCm38 --email
 --save_merged_fastq FALSE --skip_markduplicates TRUE --skip_preseq
TRUE --skip_dupradar TRUE --skip_stringtie TRUE
```

Alignments were stored as indexed BAM files. QC was performed with MultiQC v3.9.5 [13]. All samples passed bioinformatic QC thresholds and were retained for exploratory analysis. Salmon-derived gene-level expression values were imported into an R Studio environment. Variance-stabilized gene-level counts were used for exploratory analysis via principal components analysis and sample-to-sample distance calculations. Non-normalized gene-level counts were used for differential expression analysis with DESeq2 [14] following best practices as described in the DESeq2 vignette (http://bioconductor.org/packages/devel/bioc/vignettes/DESeq2/inst/doc/DESeq2.html). DESeq2 data object was created and conditioned solely on 'treatment' for DE testing. Pathway analysis was carried out using iPathwayGuide software (https://advaitabio.com/ipathwayguide) to perform proprietary "impact analysis" to generate reports describing enriched pathways, GO terms, upstream regulatory genes, and gene networks. These analyses consider both the direction and type of all signals on a pathway along with the position, role and type of each gene [15–17]. To determine statistical significance, Fisher's method was employed to compute the initial *p*-value, which was further corrected for multiple comparisons using the false discovery rate (FDR) method. All the TRAP-seq data were deposited into the National Center for Biotechnology Information – Gene Expression Omnibus (NCBI-GEO), accession number: GSE241234.

**Immunofluorescence and Imaging**

Mice were deeply anesthetized with tribromoethanol (250 – 300 mg/kg, i.p.) and then transcardially perfused with 0.01 M phosphate-buffered saline (PBS) and then 4% paraformaldehyde (PFA). After post-fixation overnight, the brains were cryopreserved and then cryosectioned at 30 µm using a Leica SM2010R microtome. For free-floating immunofluorescence, after 3X washes in PBS, brain slices were permeabilized in PBS containing 0.5% Triton X-100 for 30 min, and then blocked in a blocking buffer (PBS containing 10% normal donkey serum/NDS and 0.1% Triton X-100) for 1 h. Afterwards, slices were incubated in the blocking buffer containing primary antibodies (Table 2) with agitation at 4°C overnight. Then, after 3X washes in PBS, slices were incubated in PBS containing fluorescent dye-conjugated secondary antibodies (Table 2) for 2 h. Following the incubation and 3X washes in PBS, slices were mounted with Vectashield antifade mounting medium (Catalog No. H-1000-10; Vector Laboratories, Newark, CA, USA) on Superfrost microscope slides (Thermo Fisher Scientific) and the coverslips were sealed with nail polish. Fluorescent images were obtained using an Olympus FV3000 confocal microscope (Tokyo, Japan) or an Olympus SLIDEVIEW VS200 digital slide scanner. For LS anterograde tracing, NeuN stain was performed to mark mature neurons, EGFP- and NeuN-expressing cells in specific brain regions were counted using QuPath (version 0.4.3) software, and percent neurons receiving afferent inputs from the LS were computed. For LS retrograde tracing, RFP stain was performed, and percent immunoreactive area in each brain region with tdTomato expression was quantified using ImageJ Fiji.

**Immunolabeling-enabled three-dimensional imaging of solvent-cleared organs (iDISCO)**

The method of iDISCO has been documented previously [18]. In brief, PFA-fixed brain samples were stored in PBS containing 0.02% (w/v) $NaN_3$ at 4ºC until the iDISCO protocol (available at idisco.info) was performed. Brain hemispheres were kept individually in 2 ml amber Eppendorf tubes (Hamburg, Germany) during the process, and gentle agitation was applied at each step unless specified otherwise.

***Pretreatment***

Brain samples were dehydrated serially in a methanol/dd$H_2O$ series: 20%, 40%, 60%, 80%, 100%, and 100% for 1 h each. Chilled samples were then incubated in a mixture of 66% dichloromethane (DCM) and 33% methanol overnight. Afterwards, following 2X washes in 100% methanol, samples were bleached in fresh 5% $H_2O_2$ in methanol at 4ºC overnight. The next day, following rehydration with a methanol/dd$H_2O$ series: 80%, 60%, 40%, 20%, and PBS for 1 h each, samples were washed 2X in PBS containing 0.2% Triton X-100 (PTx.2).

***Immunolabeling***

Brain samples were permeabilized in PTx.2 containing 2.3% (w/v) glycine and 20% DMSO for 2 days at 37ºC and then blocked in PTx.2 containing 6% NDS and 10% DMSO for 2 days at 37ºC. Afterwards, samples were incubated in PBS containing 0.2% Tween 20 and 1% (w/v) heparin (PTwH) with added 5% DMSO, 3% NDS, and the primary antibody: rabbit anti-c-fos antibody (1:200 dilution; Catalog No. 226003; Synaptic Systems, Göttingen, Germany) for 5 days at 37ºC. Then, following 5X washes in PTwH overnight, samples were incubated in PTwH with 3% NDS and the secondary antibody: Alexa Fluor 647 donkey anti-rabbit antibody (1:200 dilution; Catalog No. 711-605-152; Jackson ImmunoResearch, West Grove, PA, USA) for 5 days at 37ºC. Finally, samples were washed 5X in PTwH overnight.

***Clearing***

After dehydration in a methanol/dd$H_2O$ series: 20%, 40%, 60%, 80%, and 100% for 1 h each, brain samples were incubated in the 66% DCM / 33% methanol solution for 3 h, and then washed 2X in 100% DCM for 15 min each. Finally, samples were incubated in dibenzyl ether (DBE) without agitation until they were imaged. Bloxygen was applied to prevent sample oxidation.

**Light sheet microscopy and ClearMap analysis**

Light sheet microscopy was used for imaging the DBE-cleared iDISCO brain samples, as described in detail previously [19]. Basically, a LaVision UltraMicroscope II light sheet microscope (LaVision BioTec, Bielefeld, Germany) was employed, along with an Olympus MVPLAPO 2X objective and a dipping cup with a 6-mm working distance. The brain hemispheres were imaged at 1.6X in the sagittal orientation (with the medial side on the bottom) with 3-µm z-steps. The continuous light sheet scanning method was employed, with the contrast blending algorithm for the 640nm channel (15 or 20 acquisitions per plane), and without horizontal scanning for the 480nm channel to acquire images of autofluorescence. The open-source ClearMap 1.0 software (https://github.com/ChristophKirst/ClearMap) was used to quantify c-fos positive cells, which were detected with the cell size threshold set as 20 voxels. Images were aligned to the Allen Brain Atlas (http://alleninstitute.org/) to identify brain regions.

**Chemogenetic fMRI (cg-fMRI)**

Fos-iCreER mice were stereotaxically injected with AAV8-hSyn-DIO-hM3Dq-mCherry in the LS, and went through the 4-OH-TM-enabled Fos-TRAP procedure following social interaction, which led to Cre-dependent DREADD expression solely in LS neurons activated by social interaction, i.e., the LS social engram. Four weeks later, they underwent baseline fMRI scans on an Agilent Discovery 901 7-Tesla small animal scanner under isoflurane anesthesia. FMRI time series were used to identify changes in the blood-oxygen-level dependent (BOLD) signal, which served as a proxy marker for neural activity [20]. To minimize movement artifacts, animals were anesthetized with isoflurane. Previous research has shown that resting-state networks in mouse models remain stable even under anesthesia with different narcotic agents [21]. One week after the baseline scan, they were injected with a DREADD agonist (CNO; 3 mg/kg, i.p.) to chemogenetically re-activate the LS social engram. Forty-five min later, they underwent fMRI scans to measure changes in neural activity induced by chemogenetic activation of the LS social engram neurons.

Processing of the fMRI data was performed using tools from AFNI, ANTs, and FSL [22–24]. Collected fMRI images had a starting resolution of 156 x 156 x 600 µm acquired every 1.5 seconds over 30 min, while anatomical information was taken from a fast-imaging steady-state acquisition (FIESTA) sequence with resolution of 78 x 78 x 200 µm. We applied bias field and motion correction, using the FIESTA image as the input, registered all images into Waxholm space [25] so they could be analyzed in a common reference space. Registered fMRI data were resampled to 200 µm and smoothed using an 800 µm smoothing kernel and a seed was set in the LS of each subject and correlation was calculated between the seed and all other voxels in the brain. The results of these correlations were then fed into a second-level analysis using Randomise from FSL [26], corrected for family wise error (n = 500 permutations), and had threshold free cluster enhancement applied [27].

**Statistical analysis**

All the statistical analyses were performed using GraphPad Prism 9 software (Dotmatics, Boston, MA, USA) unless indicated otherwise. Independent-samples *t* tests were used for comparisons of two groups with one independent variable. Two-way ANOVAs were employed for comparisons with two independent variables, followed by *post hoc* Bonferroni or Šidák tests when needed. Welch's correction was applied in the case of unequal variances. Statistical significance was set at $p < 0.05$ unless otherwise stated. Data are presented as mean ± SEM unless specified otherwise.

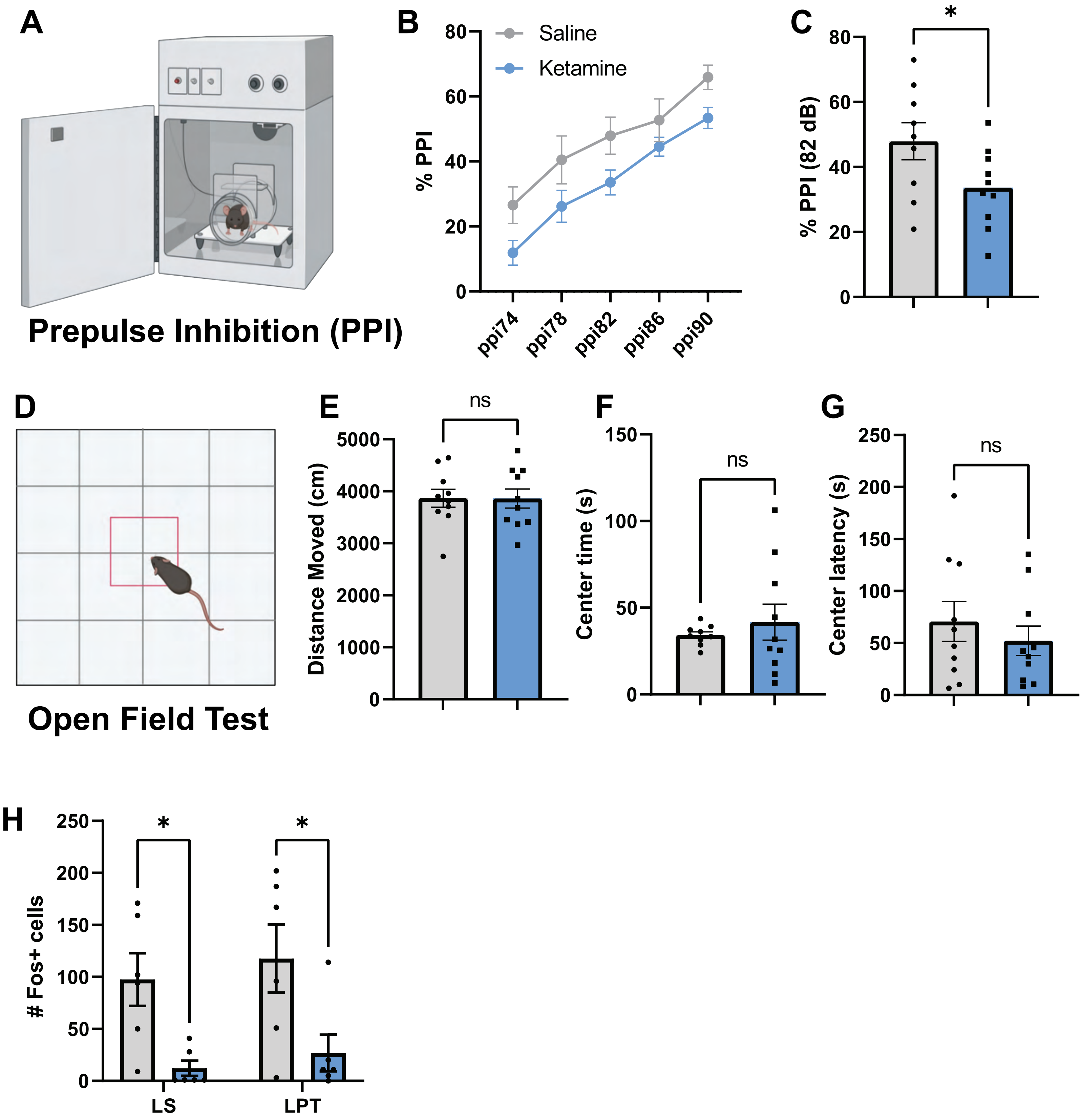


**Supplementary Figure 1: Chronic ketamine induces sensorimotor deficits and reduced thalamic activation during social interaction.**

(A) Schematic depicting operant boxes for measurement of prepulse inhibition (PPI), (B) histogram of % PPI across a range of 74-90 decibels (db) and (C) % PPI at 82 dB. (D) Schematic depicting the open field test. (E) Distance moved, (F) cumulative time in center, and (G) latency to enter the center of the open field over a 10-minute interval. (H) Histogram depicting the number of cfos-positive cells in the lateral septum (LS) and lateral posterior thalamus (LPT) during social interaction in saline and ketamine-treated mice. *p<0.05. *Parts of this figure were made using Biorender.*

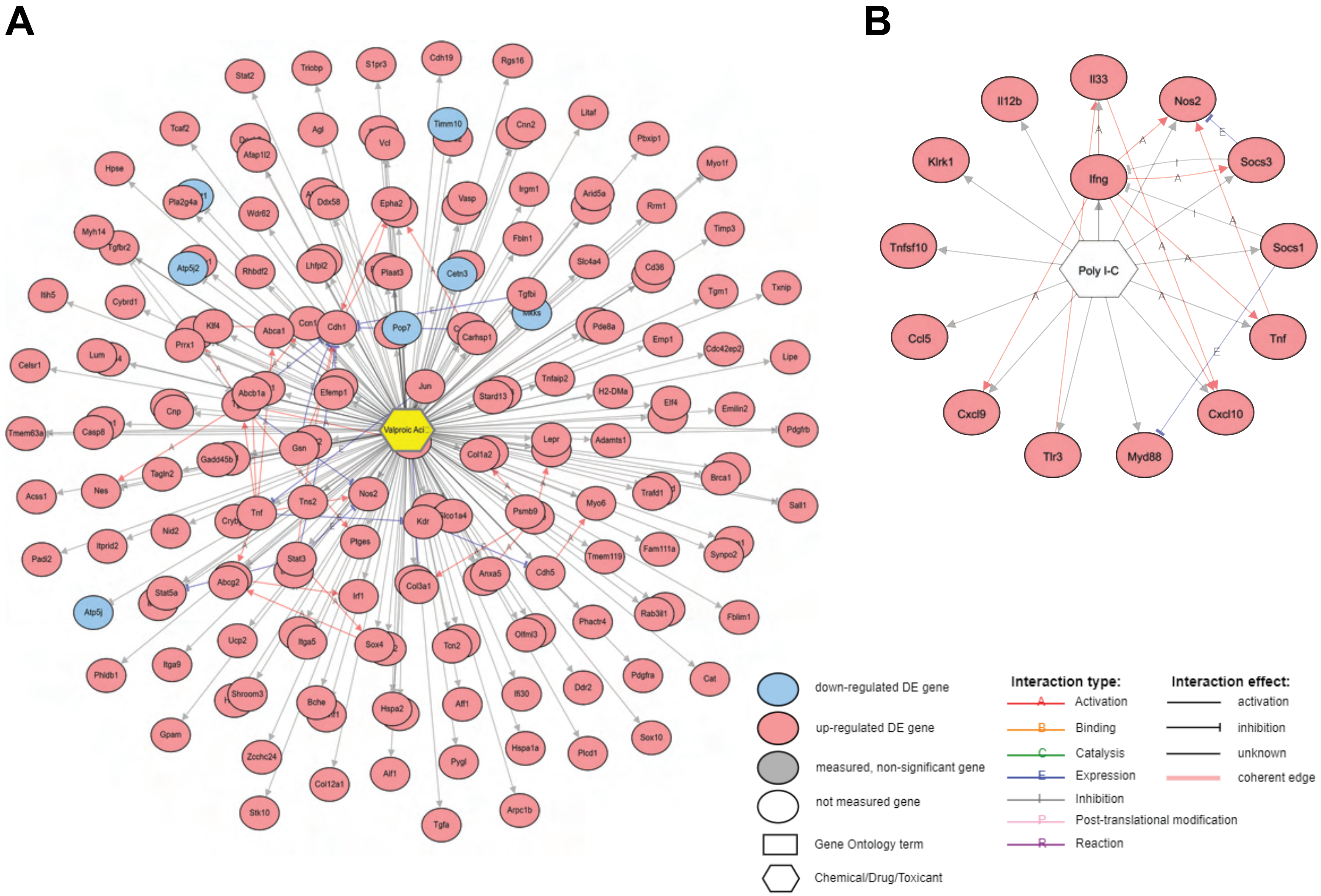


**Supplementary Figure 2. Network analysis of differentially expressed genes (DEGs) relevant to (A) valproic acid and (B) poly-IC, created via IPathwayGuide.**

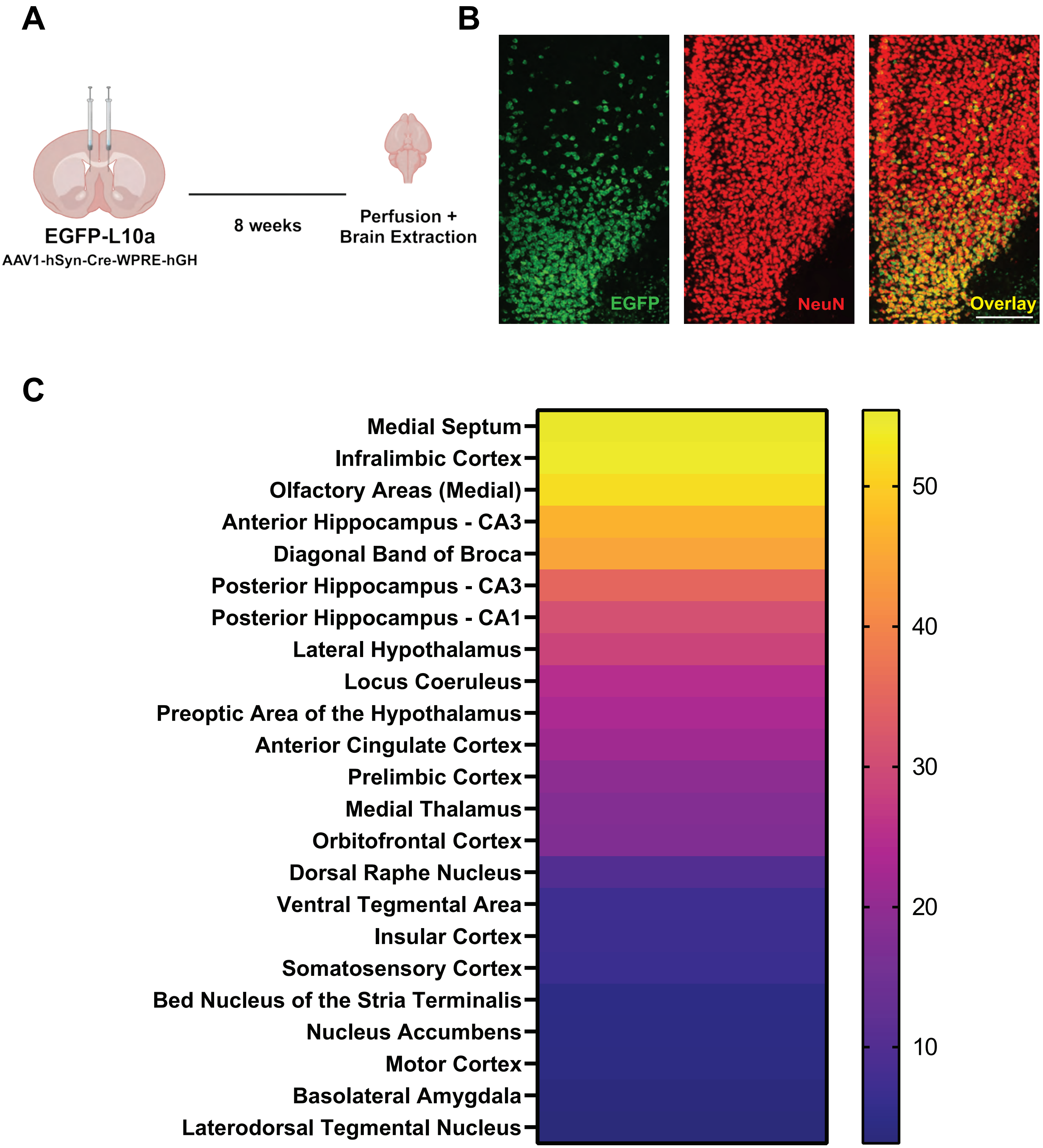


**Supplementary Figure 3. Brain regions receiving inputs from the lateral septum.**

(A) Experimental schematic of infusing AAV1-hSyn-Cre-WPRE-hGH in the lateral septum of EGFP-L10a mice, and timeline of the anterograde tracing experiment. (B) Representative confocal images of EGFP expression in the medial prefrontal cortex and NeuN stain used to quantify percentages of neurons receiving lateral septal inputs (scale bar = 200 μm). (C) Heatmap showing percent neurons receiving lateral septal inputs in 23 different brain regions. *Parts of this figure were made using Biorender.*

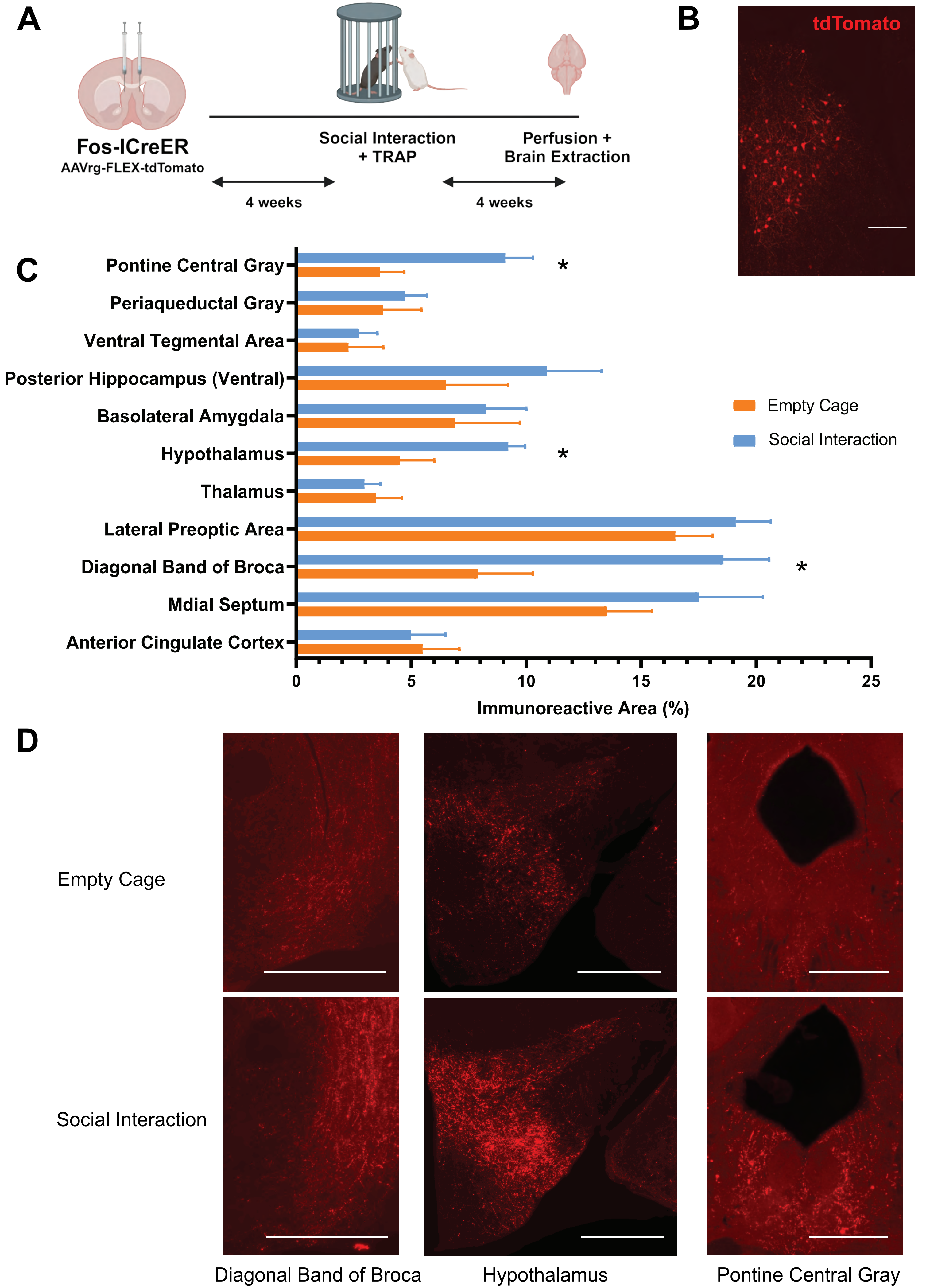
A
Fos-ICreER
AAVrg-FLEX-tdTomato
Social Interaction
+ TRAP
Perfusion +
Brain Extraction
4 weeks
4 weeks
B
tdTomato
C
Pontine Central Gray
Periaqueductal Gray
Ventral Tegmental Area
Posterior Hippocampus (Ventral)
Basolateral Amygdala
Hypothalamus
Thalamus
Lateral Preoptic Area
Diagonal Band of Broca
Mdial Septum
Anterior Cingulate Cortex
*
*
*
0
5
10
15
20
25
Immunoreactive Area (%)
Empty Cage
Social Interaction
D
Empty Cage
Social Interaction
Diagonal Band of Broca
Hypothalamus
Pontine Central Gray

**Supplementary Figure 4. Social interaction-activated afferent inputs to the lateral septum.**

(A) Experimental schematic of viral targeting of AAVrg-FLEX-tdTomato to the lateral septal neurons of Fos-iCreER mice, and timeline of the retrograde tracing experiment. (B) Representative confocal image of tdTomato expression in the right lateral septum (medial side on the left; scale bar = 200 µm). (C) Histogram showing that social interaction, versus exploration of two empty cages, led to enhanced afferent inputs from the diagonal band of Broca, hypothalamus, and pontine central gray. (D) Representative confocal images of three brain regions showing augmented afferent inputs to the lateral septum in social interaction, relative to empty cage exploration (scale bar = 800 µm). * $p < 0.05$. *Parts of this figure were made using Biorender.*

16. Khatri P, Drăghici S. Ontological analysis of gene expression data: current tools,

limitations, and open problems. Bioinformatics. 2005;21:3587–3595.
