## Supplementary Table 1 for "Lateral Septal Circuits Govern Schizophrenia-Like Effects of Ketamine on Social Behavior"

**Supplementary Table 1 - Upregulated Translatome**

| symbol | entrez | logfc | adjpv | |
|---|---|---|---|---|
| Gm26444 | 1.15E+08 | 10 | 0.000001 | 1 |
| Igkv9-123 | 628144 | 10 | 0.010657 | 2 |
| Igkv3-4 | 626347 | 10 | 0.015162 | 3 |
| Ighv14-2 | 668421 | 9.218229 | 0.02157 | 4 |
| Ighv5-16 | 633568 | 8.371297 | 2.66E-05 | 5 |
| Ighv9-3 | 780825 | 8.296644 | 0.001484 | 6 |
| Gm18853 | 1E+08 | 8.257836 | 2.45E-05 | 7 |
| Ighv9-1 | 195180 | 8.236317 | 0.000186 | 8 |
| Ighv1-78 | 213570 | 7.988883 | 5.24E-06 | 9 |
| Gm3194 | 1.01E+08 | 7.696275 | 1.60E-05 | 10 |
| Ighv1-76 | 1.01E+08 | 7.530187 | 0.001183 | 11 |
| Ighv1-85 | 668597 | 6.95361 | 0.000001 | 12 |
| Pou4f1 | 18996 | 6.903037 | 0.00611 | 13 |
| Ighv5-4 | 777688 | 6.285668 | 5.49E-05 | 14 |
| Igkv9-120 | 434025 | 6.205804 | 0.033101 | 15 |
| Arhgap30 | 226652 | 6.078839 | 0.008161 | 16 |
| Ighv1-19 | 382692 | 6.007543 | 0.000001 | 17 |
| Ighv8-12 | 780960 | 5.774182 | 0.038595 | 18 |
| Igkv3-7 | 108005 | 5.576149 | 0.005208 | 19 |
| Ighv1-12 | 629860 | 5.525492 | 0.043921 | 20 |
| Ighv1-15 | 629865 | 5.507948 | 0.001695 | 21 |
| Ighv2-2 | 777686 | 5.008214 | 8.70E-06 | 22 |
| Bcl2l14 | 66813 | 4.829711 | 0.005238 | 23 |
| Ltf | 17002 | 4.82613 | 0.007724 | 24 |
| Ighv1-75 | 622728 | 4.68795 | 0.001369 | 25 |
| Ighv4-1 | 780802 | 4.499996 | 0.013913 | 26 |
| Irx2 | 16372 | 4.358301 | 0.010188 | 27 |
| Ighv1-39 | 780891 | 4.167919 | 0.000001 | 28 |
| Ighv1-64 | 380823 | 4.104666 | 0.000001 | 29 |
| Flnc | 68794 | 4.083767 | 0.000001 | 30 |
| Mefv | 54483 | 4.041774 | 0.006801 | 31 |
| Gbp2b | 14468 | 4.038356 | 0.001561 | 32 |
| Nos2 | 18126 | 3.992887 | 0.000001 | 33 |
| Cobll1 | 319876 | 3.938688 | 0.030987 | 34 |
| Gm4117 | 1E+08 | 3.827226 | 0.039235 | 35 |
| Klrc1 | 16641 | 3.522934 | 0.000329 | 36 |
| Ighd | 380797 | 3.522449 | 0.004648 | 37 |
| Sirpb1c | 1E+08 | 3.501301 | 2.39E-05 | 38 |
| Pydc3 | 1E+08 | 3.485688 | 0.000001 | 39 |
| Iglv3 | 404743 | 3.471378 | 0.012624 | 40 |
| Gm9917 | 1E+08 | 3.460918 | 0.008752 | 41 |
| Hspg2 | 15530 | 3.421629 | 0.000001 | 42 |
| Parp14 | 547253 | 3.412003 | 0.000607 | 43 |
| Mki67 | 17345 | 3.411789 | 0.00012 | 44 |
| Kif18b | 70218 | 3.374608 | 2.17E-05 | 45 |

| | | | | |
|---|---|---|---|---|
| Tgm1 | 21816 | 3.368196 | 5.69E-05 | 46 |
| Dlgap5 | 218977 | 3.362604 | 3.34E-06 | 47 |
| Themis | 210757 | 3.331611 | 0.000348 | 48 |
| Kmo | 98256 | 3.318773 | 0.000714 | 49 |
| Itgax | 16411 | 3.309001 | 0.000763 | 50 |
| Gbp6 | 100702 | 3.305756 | 0.000001 | 51 |
| Gpr141 | 353346 | 3.301403 | 0.046678 | 52 |
| Cfb | 14962 | 3.293253 | 5.20E-05 | 53 |
| Neil3 | 234258 | 3.239535 | 0.031394 | 54 |
| Ighv1-55 | 780932 | 3.23044 | 0.00708 | 55 |
| Adam8 | 11501 | 3.230243 | 0.000403 | 56 |
| Igf2bp3 | 140488 | 3.158431 | 0.00872 | 57 |
| Ifi205 | 226695 | 3.156529 | 2.65E-06 | 58 |
| Ifng | 15978 | 3.134495 | 0.020988 | 59 |
| Top2a | 21973 | 3.109085 | 0.003567 | 60 |
| AI607873 | 226691 | 3.10331 | 0.000001 | 61 |
| Igkv17-121 | 667435 | 3.082932 | 0.00065 | 62 |
| Gm12250 | 631323 | 3.078556 | 0.000001 | 63 |
| Esco2 | 71988 | 3.073002 | 0.002379 | 64 |
| Sell | 20343 | 3.051799 | 0.001332 | 65 |
| Uhrf1 | 18140 | 3.030924 | 0.000001 | 66 |
| Ighv1-52 | 382696 | 3.020935 | 0.031409 | 67 |
| Tnfrsf9 | 21942 | 3.011927 | 0.000313 | 68 |
| Helz2 | 229003 | 2.997481 | 0.000001 | 69 |
| Hk3 | 212032 | 2.985293 | 0.000001 | 70 |
| Apol6 | 71939 | 2.978998 | 1.92E-05 | 71 |
| Gm4955 | 240921 | 2.977326 | 0.000001 | 72 |
| Ighv1-42 | 629906 | 2.957982 | 0.005016 | 73 |
| Col4a6 | 94216 | 2.956875 | 0.000001 | 74 |
| Mmp25 | 240047 | 2.940887 | 0.000001 | 75 |
| Atp8b4 | 241633 | 2.928702 | 0.006558 | 76 |
| Tgtp1 | 21822 | 2.914726 | 0.000224 | 77 |
| Cxcl11 | 56066 | 2.912326 | 0.002033 | 78 |
| Gm12185 | 620913 | 2.892685 | 0.000001 | 79 |
| Plac8 | 231507 | 2.880362 | 4.60E-06 | 80 |
| Aoah | 27052 | 2.847741 | 0.000331 | 81 |
| C1s1 | 50908 | 2.839757 | 0.000001 | 82 |
| Casc5 | 76464 | 2.83168 | 5.59E-05 | 83 |
| F5 | 14067 | 2.820625 | 0.015641 | 84 |
| Il12rb1 | 16161 | 2.810476 | 0.000001 | 85 |
| Ccl19 | 24047 | 2.807809 | 0.001977 | 86 |
| Gimap1 | 16205 | 2.806179 | 0.006076 | 87 |
| A2m | 232345 | 2.801263 | 0.000186 | 88 |
| Itgb2 | 16414 | 2.800696 | 0.000427 | 89 |
| Gbp2 | 14469 | 2.799414 | 0.000362 | 90 |
| Tlr9 | 81897 | 2.797425 | 0.000001 | 91 |
| Col20a1 | 73368 | 2.795094 | 0.000261 | 92 |

| | | | | |
|---|---|---|---|---|
| Slfn5 | 327978 | 2.788264 | 0.000001 | 93 |
| Osmr | 18414 | 2.77381 | 0.000001 | 94 |
| Gbp5 | 229898 | 2.769677 | 0.000504 | 95 |
| Ifi47 | 15953 | 2.760063 | 2.93E-05 | 96 |
| Pbk | 52033 | 2.753057 | 0.002351 | 97 |
| Gm1966 | 434223 | 2.749896 | 0.000001 | 98 |
| Myo1g | 246177 | 2.748887 | 0.000001 | 99 |
| Ttn | 22138 | 2.742402 | 0.003613 | 100 |
| Tlr8 | 170744 | 2.735325 | 0.029283 | 101 |
| Vcam1 | 22329 | 2.733504 | 0.001084 | 102 |
| Vim | 22352 | 2.727555 | 0.001015 | 103 |
| Ccr7 | 12775 | 2.723948 | 0.00312 | 104 |
| Ccl7 | 20306 | 2.716389 | 0.003905 | 105 |
| Nusap1 | 108907 | 2.713556 | 0.000112 | 106 |
| H2-DMb2 | 15000 | 2.702125 | 0.000116 | 107 |
| Pyhin1 | 236312 | 2.697993 | 0.000001 | 108 |
| C4b | 12268 | 2.696941 | 0.004127 | 109 |
| Wdfy4 | 545030 | 2.694782 | 0.000001 | 110 |
| C1ra | 50909 | 2.693037 | 0.000001 | 111 |
| Cdc6 | 23834 | 2.692314 | 0.000957 | 112 |
| BC035044 | 232406 | 2.691075 | 0.005373 | 113 |
| Gzmb | 14939 | 2.686433 | 0.000131 | 114 |
| Oas3 | 246727 | 2.684474 | 0.000001 | 115 |
| Dnase1l3 | 13421 | 2.671092 | 0.000001 | 116 |
| Slfn1 | 20555 | 2.669079 | 0.000001 | 117 |
| Pole | 18973 | 2.637568 | 0.000001 | 118 |
| Mnda | 381308 | 2.624797 | 0.000001 | 119 |
| Ciita | 12265 | 2.622807 | 0.000001 | 120 |
| Ms4a6b | 69774 | 2.617466 | 0.000001 | 121 |
| Irf1 | 16362 | 2.615555 | 0.000667 | 122 |
| Siglec1 | 20612 | 2.609439 | 0.000001 | 123 |
| Tlr6 | 21899 | 2.604556 | 0.003242 | 124 |
| Itgal | 16408 | 2.60454 | 0.000001 | 125 |
| Sh2d2a | 27371 | 2.597935 | 0.000118 | 126 |
| Upp1 | 22271 | 2.597805 | 8.32E-05 | 127 |
| Fcgr4 | 246256 | 2.595484 | 1.20E-06 | 128 |
| Cysltr1 | 58861 | 2.593026 | 0.006038 | 129 |
| Ticrr | 77011 | 2.58053 | 0.017284 | 130 |
| Cdh1 | 12550 | 2.579835 | 0.001064 | 131 |
| C3 | 12266 | 2.576751 | 5.90E-06 | 132 |
| Icam1 | 15894 | 2.571007 | 0.000001 | 133 |
| Ccl2 | 20296 | 2.569438 | 2.49E-05 | 134 |
| Il12b | 16160 | 2.564225 | 0.00733 | 135 |
| Plbd1 | 66857 | 2.563352 | 0.000001 | 136 |
| Trim25 | 217069 | 2.562772 | 0.000704 | 137 |
| Ifi204 | 15951 | 2.562386 | 0.000001 | 138 |
| Alpk1 | 71481 | 2.560889 | 1.09E-05 | 139 |

| | | | | |
|---|---|---|---|---|
| Gata3 | 14462 | 2.557785 | 0.008833 | 140 |
| Ggt5 | 23887 | 2.556518 | 0.002948 | 141 |
| Rhbdf2 | 217344 | 2.554955 | 9.63E-06 | 142 |
| Prf1 | 18646 | 2.554007 | 0.023156 | 143 |
| Il18rap | 16174 | 2.550939 | 0.003906 | 144 |
| Cd244 | 18106 | 2.548421 | 0.023526 | 145 |
| Ubash3a | 328795 | 2.5446 | 0.011424 | 146 |
| Has2 | 15117 | 2.54434 | 0.002658 | 147 |
| Melk | 17279 | 2.543415 | 0.006574 | 148 |
| Ctsw | 13041 | 2.539919 | 0.002364 | 149 |
| Misp | 78906 | 2.538096 | 0.022265 | 150 |
| Irgm2 | 54396 | 2.53136 | 0.000151 | 151 |
| Gm8995 | 668139 | 2.524425 | 0.000001 | 152 |
| Col11a2 | 12815 | 2.521555 | 0.000001 | 153 |
| Serpina3f | 238393 | 2.520455 | 0.000001 | 154 |
| Robo4 | 74144 | 2.519523 | 0.003264 | 155 |
| Rbm47 | 245945 | 2.51527 | 0.000001 | 156 |
| Cp | 12870 | 2.494687 | 0.000001 | 157 |
| Xdh | 22436 | 2.493422 | 0.000001 | 158 |
| Grap2 | 17444 | 2.490029 | 0.000194 | 159 |
| Csf2rb2 | 12984 | 2.489553 | 0.000001 | 160 |
| Tnfrsf14 | 230979 | 2.487656 | 0.002963 | 161 |
| Gvin1 | 74558 | 2.485645 | 0.000001 | 162 |
| Iigp1 | 60440 | 2.485167 | 0.000161 | 163 |
| Irg1 | 16365 | 2.484096 | 4.57E-06 | 164 |
| Scarf1 | 380713 | 2.483407 | 0.02219 | 165 |
| Oasl1 | 231655 | 2.482002 | 0.000001 | 166 |
| Themis2 | 230787 | 2.481765 | 0.000001 | 167 |
| Gbp9 | 236573 | 2.477778 | 0.000001 | 168 |
| Parp3 | 235587 | 2.476096 | 0.000001 | 169 |
| Rnf213 | 672511 | 2.474642 | 0.000001 | 170 |
| Casp12 | 12364 | 2.473805 | 0.001332 | 171 |
| Dock2 | 94176 | 2.468138 | 0.000001 | 172 |
| Kcnj13 | 1E+08 | 2.458651 | 0.001946 | 173 |
| Gm4070 | 1E+08 | 2.450047 | 0.049215 | 174 |
| Clcf1 | 56708 | 2.444972 | 0.019457 | 175 |
| Amica1 | 270152 | 2.436648 | 0.005222 | 176 |
| Pdcd1 | 18566 | 2.435206 | 0.000591 | 177 |
| Phf11a | 219131 | 2.434826 | 0.015894 | 178 |
| Itgb7 | 16421 | 2.43419 | 1.44E-05 | 179 |
| Fn1 | 14268 | 2.433275 | 0.000001 | 180 |
| Il1rn | 16181 | 2.427703 | 0.019505 | 181 |
| Ptx3 | 19288 | 2.424681 | 0.006411 | 182 |
| Gbp10 | 626578 | 2.424376 | 0.0024 | 183 |
| Card11 | 108723 | 2.419546 | 2.65E-06 | 184 |
| Pydc4 | 623121 | 2.419462 | 2.66E-05 | 185 |
| Cd69 | 12515 | 2.418717 | 0.001427 | 186 |

| | | | | |
|---|---|---|---|---|
| Itgb4 | 192897 | 2.417751 | 0.000001 | 187 |
| Tlr12 | 384059 | 2.409508 | 0.010114 | 188 |
| Tnfaip2 | 21928 | 2.407676 | 4.26E-06 | 189 |
| Kif15 | 209737 | 2.407111 | 0.00118 | 190 |
| Col4a5 | 12830 | 2.406942 | 0.000001 | 191 |
| Il2rb | 16185 | 2.40455 | 0.000001 | 192 |
| Mvp | 78388 | 2.400596 | 0.000001 | 193 |
| E2f2 | 242705 | 2.399127 | 0.003337 | 194 |
| Sp100 | 20684 | 2.394319 | 0.000001 | 195 |
| Mndal | 1E+08 | 2.393569 | 0.000001 | 196 |
| Parp9 | 80285 | 2.392967 | 0.000914 | 197 |
| Ly75 | 17076 | 2.389824 | 1.03E-05 | 198 |
| Klk6 | 19144 | 2.387566 | 0.001075 | 199 |
| Ms4a4b | 60361 | 2.379922 | 0.0004 | 200 |
| AU020206 | 1.01E+08 | 2.376658 | 0.000393 | 201 |
| 9930111J2 | 245240 | 2.376252 | 0.000001 | 202 |
| Gsdmd | 69146 | 2.375917 | 1.73E-05 | 203 |
| F830016B0 | 240328 | 2.373733 | 0.000001 | 204 |
| Cd300lf | 246746 | 2.373457 | 0.001253 | 205 |
| Rasal3 | 320484 | 2.372186 | 0.000001 | 206 |
| Rrm2 | 20135 | 2.370441 | 0.000407 | 207 |
| Fam83g | 69640 | 2.369549 | 0.028484 | 208 |
| Gm16340 | 1.01E+08 | 2.368279 | 0.045963 | 209 |
| Bcl3 | 12051 | 2.34975 | 5.19E-05 | 210 |
| Clca3a1 | 12722 | 2.347967 | 0.004563 | 211 |
| Mlkl | 74568 | 2.347368 | 0.00144 | 212 |
| Cd180 | 17079 | 2.345822 | 1.25E-06 | 213 |
| Slfn9 | 237886 | 2.343497 | 0.000001 | 214 |
| Gbp8 | 76074 | 2.334792 | 3.19E-06 | 215 |
| Nid2 | 18074 | 2.325826 | 0.000001 | 216 |
| Bub1 | 12235 | 2.324899 | 0.027401 | 217 |
| Ms4a6c | 73656 | 2.319001 | 2.04E-05 | 218 |
| Cilp | 214425 | 2.318682 | 0.019244 | 219 |
| Cd36 | 12491 | 2.317083 | 0.001005 | 220 |
| Hk2 | 15277 | 2.31495 | 0.000001 | 221 |
| Prdm1 | 12142 | 2.314226 | 7.07E-06 | 222 |
| Trim12c | 319236 | 2.313389 | 1.74E-05 | 223 |
| Fgl2 | 14190 | 2.310378 | 0.000001 | 224 |
| Cd40 | 21939 | 2.3069 | 0.003515 | 225 |
| Tgm2 | 21817 | 2.305332 | 0.000001 | 226 |
| Igtp | 16145 | 2.303486 | 0.000193 | 227 |
| Duox2 | 214593 | 2.302786 | 0.000001 | 228 |
| Gm4841 | 225594 | 2.290306 | 0.000001 | 229 |
| Ttk | 22137 | 2.289674 | 0.012304 | 230 |
| Naip6 | 17952 | 2.28761 | 0.003293 | 231 |
| Mcm3 | 17215 | 2.285698 | 3.61E-06 | 232 |
| Ptprc | 19264 | 2.282079 | 0.000001 | 233 |

| | | | | |
|---|---|---|---|---|
| Rgs1 | 50778 | 2.281863 | 0.031819 | 234 |
| 9930111J2 | 667214 | 2.27971 | 0.000001 | 235 |
| Cdk1 | 12534 | 2.274951 | 0.026686 | 236 |
| Gimap4 | 107526 | 2.272518 | 0.000307 | 237 |
| Tmem106a | 217203 | 2.268601 | 0.000001 | 238 |
| Mpeg1 | 17476 | 2.265938 | 0.002634 | 239 |
| Ighv1-18 | 629871 | 2.265613 | 0.000215 | 240 |
| Acp5 | 11433 | 2.265533 | 0.028717 | 241 |
| Tgtp2 | 1E+08 | 2.264121 | 0.000222 | 242 |
| Flnb | 286940 | 2.263114 | 0.000001 | 243 |
| Lcn2 | 16819 | 2.262775 | 0.00313 | 244 |
| Gm5431 | 432555 | 2.25989 | 1.00E-05 | 245 |
| Socs3 | 12702 | 2.24427 | 0.012884 | 246 |
| Col16a1 | 107581 | 2.241668 | 1.92E-06 | 247 |
| Ccr2 | 12772 | 2.241638 | 0.004357 | 248 |
| Dhx58 | 80861 | 2.228453 | 1.78E-06 | 249 |
| Loxl2 | 94352 | 2.223684 | 0.009536 | 250 |
| Ighg2b | 16016 | 2.221538 | 0.007755 | 251 |
| Ncf2 | 17970 | 2.215537 | 4.81E-06 | 252 |
| Tagap | 72536 | 2.2145 | 2.52E-05 | 253 |
| Brip1 | 237911 | 2.210874 | 0.014719 | 254 |
| Arhgef5 | 54324 | 2.207047 | 0.000114 | 255 |
| Anpep | 16790 | 2.205309 | 0.000312 | 256 |
| Cdca2 | 108912 | 2.20504 | 0.048587 | 257 |
| Procr | 19124 | 2.203451 | 0.035127 | 258 |
| Flna | 192176 | 2.203333 | 6.06E-05 | 259 |
| Acap1 | 216859 | 2.202925 | 0.002715 | 260 |
| Psd4 | 215632 | 2.202173 | 7.72E-05 | 261 |
| Uba7 | 74153 | 2.197638 | 0.000001 | 262 |
| Batf2 | 74481 | 2.197613 | 0.000153 | 263 |
| Dennd1c | 70785 | 2.197443 | 0.002968 | 264 |
| Tigit | 1E+08 | 2.189634 | 0.049104 | 265 |
| Ercc6l | 236930 | 2.187049 | 0.016314 | 266 |
| Ndc80 | 67052 | 2.185718 | 0.023213 | 267 |
| Tgfbi | 21810 | 2.180819 | 0.000001 | 268 |
| Naalad2 | 72560 | 2.180764 | 0.005506 | 269 |
| Asap3 | 230837 | 2.180204 | 2.98E-06 | 270 |
| Napsa | 16541 | 2.179889 | 0.015312 | 271 |
| Ptpn6 | 15170 | 2.178076 | 0.000001 | 272 |
| Gzma | 14938 | 2.172329 | 0.015726 | 273 |
| Ifi203 | 15950 | 2.169895 | 0.000001 | 274 |
| Osgin1 | 71839 | 2.169524 | 0.02816 | 275 |
| Aldh1l1 | 107747 | 2.16745 | 0.013997 | 276 |
| Gbp7 | 229900 | 2.167337 | 0.001004 | 277 |
| Notch2 | 18129 | 2.1619 | 7.04E-05 | 278 |
| Hcar2 | 80885 | 2.161692 | 0.001502 | 279 |
| Sox17 | 20671 | 2.158916 | 0.028228 | 280 |

| | | | | |
|---|---|---|---|---|
| Hist1h3c | 319148 | 2.156877 | 3.18E-05 | 281 |
| Gm4951 | 240327 | 2.156687 | 0.000001 | 282 |
| Cdh5 | 12562 | 2.154938 | 4.26E-05 | 283 |
| Baz1a | 217578 | 2.154529 | 0.000001 | 284 |
| Aspm | 12316 | 2.154067 | 0.014828 | 285 |
| Naip2 | 17948 | 2.152523 | 5.58E-06 | 286 |
| Elf4 | 56501 | 2.145094 | 0.000001 | 287 |
| Was | 22376 | 2.143486 | 0.00108 | 288 |
| Igkv5-43 | 381783 | 2.141072 | 0.004917 | 289 |
| Sp110 | 109032 | 2.136672 | 8.29E-06 | 290 |
| Adgrv1 | 110789 | 2.13434 | 0.002 | 291 |
| Cd74 | 16149 | 2.133144 | 0.000906 | 292 |
| Pik3ap1 | 83490 | 2.131877 | 0.000001 | 293 |
| Tubb6 | 67951 | 2.130019 | 0.001861 | 294 |
| Fanci | 208836 | 2.128487 | 0.039334 | 295 |
| Cd8b1 | 12526 | 2.127862 | 0.005308 | 296 |
| Klra2 | 16633 | 2.127683 | 0.019365 | 297 |
| Cd22 | 12483 | 2.126463 | 0.003863 | 298 |
| Pla2g4a | 18783 | 2.12512 | 3.88E-06 | 299 |
| Cxcl9 | 17329 | 2.12248 | 0.000001 | 300 |
| Tnc | 21923 | 2.122081 | 0.000001 | 301 |
| Ifih1 | 71586 | 2.121483 | 0.000109 | 302 |
| Tap1 | 21354 | 2.121133 | 0.000276 | 303 |
| Plxnb3 | 140571 | 2.114856 | 0.000001 | 304 |
| Cd8a | 12525 | 2.113122 | 7.58E-05 | 305 |
| Klrk1 | 27007 | 2.1105 | 0.008906 | 306 |
| Aox1 | 11761 | 2.110017 | 0.000001 | 307 |
| Trim30a | 20128 | 2.108803 | 0.000001 | 308 |
| Pirb | 18733 | 2.101425 | 0.000138 | 309 |
| H2-Ab1 | 14961 | 2.10057 | 0.001175 | 310 |
| Arhgap4 | 171207 | 2.093576 | 0.000001 | 311 |
| Pcolce | 18542 | 2.092701 | 6.44E-06 | 312 |
| Hcls1 | 15163 | 2.091887 | 1.06E-06 | 313 |
| Plcg2 | 234779 | 2.089278 | 5.40E-06 | 314 |
| Kif2c | 73804 | 2.087098 | 0.035127 | 315 |
| Notch1 | 18128 | 2.081296 | 0.001722 | 316 |
| Msn | 17698 | 2.075753 | 0.000564 | 317 |
| Nlrp1a | 195046 | 2.075632 | 0.04885 | 318 |
| H2-Q4 | 15015 | 2.075295 | 0.000001 | 319 |
| Nlrc5 | 434341 | 2.069405 | 0.000001 | 320 |
| Nfkb2 | 18034 | 2.068928 | 0.000001 | 321 |
| Spn | 20737 | 2.068888 | 0.000153 | 322 |
| Slc43a3 | 58207 | 2.067981 | 0.002364 | 323 |
| Tnfsf10 | 22035 | 2.067446 | 0.000001 | 324 |
| Casp4 | 12363 | 2.065285 | 0.000297 | 325 |
| Lcp1 | 18826 | 2.065105 | 0.00096 | 326 |
| Trim21 | 20821 | 2.063678 | 0.012628 | 327 |

| | | | | | |
|---|---|---|---|---|---|
| Gbp3 | 55932 | 2.061612 | 0.005846 | | 328 |
| Ddx60 | 234311 | 2.061467 | 0.000001 | | 329 |
| Zfp781 | 331188 | 2.058017 | 0.008578 | | 330 |
| Lamb2 | 16779 | 2.056091 | 0.000001 | | 331 |
| Tcaf2 | 232748 | 2.05532 | 0.002207 | | 332 |
| BC023105 | 667597 | 2.054778 | 0.010683 | | 333 |
| Serping1 | 12258 | 2.054422 | 0.000001 | | 334 |
| Mocos | 68591 | 2.048329 | 0.002908 | | 335 |
| Col4a4 | 12829 | 2.046499 | 0.00733 | | 336 |
| Sp140 | 434484 | 2.045845 | 0.001595 | | 337 |
| Csf1r | 12978 | 2.045074 | 0.035497 | | 338 |
| E2f8 | 108961 | 2.044368 | 0.010042 | | 339 |
| Cybb | 13058 | 2.04417 | 0.000001 | | 340 |
| Vav1 | 22324 | 2.043735 | 2.19E-06 | | 341 |
| Tbx21 | 57765 | 2.041748 | 0.02343 | | 342 |
| Gbp4 | 17472 | 2.041475 | 0.000001 | | 343 |
| Gm20045 | 1.05E+08 | 2.040629 | 0.012887 | | 344 |
| Ltbp1 | 268977 | 2.034447 | 0.000001 | | 345 |
| Ly6c2 | 1E+08 | 2.031663 | 0.000001 | | 346 |
| Inpp5d | 16331 | 2.02741 | 0.0042 | | 347 |
| AB124611 | 382062 | 2.026624 | 0.000129 | | 348 |
| Apobec3 | 80287 | 2.022564 | 0.000001 | | 349 |
| Kif22 | 110033 | 2.017506 | 0.013906 | | 350 |
| Cd86 | 12524 | 2.015735 | 4.40E-05 | | 351 |
| Spag17 | 74362 | 2.014245 | 0.043365 | | 352 |
| Cnr2 | 12802 | 2.009336 | 0.018119 | | 353 |
| Sash3 | 74131 | 2.008654 | 0.000364 | | 354 |
| Frem2 | 242022 | 2.007762 | 3.27E-05 | | 355 |
| Fermt3 | 108101 | 2.003365 | 1.65E-06 | | 356 |
| Bcl2a1a | 12044 | 2.002494 | 0.013505 | | 357 |
| Trf | 22041 | 2.002218 | 1.36E-06 | | 358 |
| Tnf | 21926 | 1.997027 | 0.013952 | | 359 |
| H2-T24 | 15042 | 1.993963 | 1.85E-05 | | 360 |
| Irf8 | 15900 | 1.990792 | 0.000001 | | 361 |
| Tbxas1 | 21391 | 1.987428 | 0.006801 | | 362 |
| Evi2b | 216984 | 1.985631 | 0.034713 | | 363 |
| Runx3 | 12399 | 1.983034 | 0.003255 | | 364 |
| Tlr2 | 24088 | 1.982082 | 8.56E-06 | | 365 |
| Fcgr2b | 14130 | 1.981704 | 0.000001 | | 366 |
| Casp8 | 12370 | 1.97758 | 8.74E-05 | | 367 |
| Cd2 | 12481 | 1.973922 | 0.028904 | | 368 |
| Slamf7 | 75345 | 1.969611 | 3.16E-06 | | 369 |
| Parp12 | 243771 | 1.967547 | 0.002485 | | 370 |
| Tmem173 | 72512 | 1.96701 | 3.14E-05 | | 371 |
| Fblim1 | 74202 | 1.962719 | 0.000417 | | 372 |
| Pear1 | 73182 | 1.962631 | 0.006232 | | 373 |
| Plekha4 | 69217 | 1.961408 | 0.000117 | | 374 |

| | | | | |
|---|---|---|---|---|
| Rhoh | 74734 | 1.960254 | 0.007783 | 375 |
| Ccrl2 | 54199 | 1.96019 | 0.013494 | 376 |
| Clspn | 269582 | 1.958722 | 0.001128 | 377 |
| Il1a | 16175 | 1.956443 | 0.016995 | 378 |
| Col3a1 | 12825 | 1.955836 | 0.001799 | 379 |
| Cspg4 | 121021 | 1.952288 | 0.034165 | 380 |
| Il2rg | 16186 | 1.951356 | 2.07E-06 | 381 |
| Cxcr6 | 80901 | 1.950231 | 0.014089 | 382 |
| Adgre5 | 26364 | 1.949824 | 0.000001 | 383 |
| Cd6 | 12511 | 1.949749 | 0.001332 | 384 |
| Nod1 | 107607 | 1.94778 | 0.000001 | 385 |
| Fosb | 14282 | 1.945677 | 0.036117 | 386 |
| Anxa4 | 11746 | 1.943705 | 2.02E-06 | 387 |
| H2-K1 | 14972 | 1.943116 | 0.001721 | 388 |
| Trim15 | 69097 | 1.941134 | 0.025464 | 389 |
| Samsn1 | 67742 | 1.940368 | 0.001012 | 390 |
| Dtx3l | 209200 | 1.937528 | 0.001962 | 391 |
| Gpnmb | 93695 | 1.928951 | 1.31E-05 | 392 |
| Vwa5a | 67776 | 1.927898 | 0.000663 | 393 |
| Cfh | 12628 | 1.926893 | 0.013004 | 394 |
| Cd274 | 60533 | 1.925192 | 0.000001 | 395 |
| Dock8 | 76088 | 1.921641 | 0.000001 | 396 |
| Irgm1 | 15944 | 1.920399 | 0.000555 | 397 |
| H2-Aa | 14960 | 1.919313 | 0.000001 | 398 |
| Mx1 | 17857 | 1.918132 | 0.000001 | 399 |
| Lilrb4 | 14728 | 1.918073 | 0.000558 | 400 |
| Ssc5d | 269855 | 1.916092 | 0.002367 | 401 |
| Gbp11 | 634650 | 1.915175 | 0.000669 | 402 |
| Mfrp | 259172 | 1.908912 | 0.0042 | 403 |
| H2-Q5 | 15016 | 1.907094 | 0.000736 | 404 |
| Ect2 | 13605 | 1.906655 | 0.018479 | 405 |
| Trim5 | 667823 | 1.904181 | 0.008688 | 406 |
| C2 | 12263 | 1.897682 | 0.000269 | 407 |
| Parp4 | 328417 | 1.897017 | 0.000001 | 408 |
| Rsad2 | 58185 | 1.89344 | 0.000001 | 409 |
| C1rl | 232371 | 1.893277 | 0.009528 | 410 |
| Cst7 | 13011 | 1.892003 | 0.000448 | 411 |
| Ly9 | 17085 | 1.891363 | 0.000442 | 412 |
| 2810417H1 | 68026 | 1.887381 | 0.036906 | 413 |
| Ifit3 | 15959 | 1.88683 | 0.008179 | 414 |
| Lag3 | 16768 | 1.886323 | 0.024981 | 415 |
| Zbp1 | 58203 | 1.884216 | 1.14E-06 | 416 |
| Plekhg2 | 101497 | 1.882361 | 4.92E-05 | 417 |
| Abcc3 | 76408 | 1.879534 | 0.004676 | 418 |
| Slfn8 | 276950 | 1.875121 | 0.000001 | 419 |
| Naip5 | 17951 | 1.87364 | 0.003634 | 420 |
| Slfn2 | 20556 | 1.873366 | 4.09E-05 | 421 |

| | | | | |
|---|---|---|---|---|
| Slfn10-ps | 237887 | 1.87317 | 0.026946 | 422 |
| Trac | 1E+08 | 1.871972 | 0.000427 | 423 |
| Ikzf3 | 22780 | 1.871374 | 0.000992 | 424 |
| Atf3 | 11910 | 1.869537 | 0.000001 | 425 |
| St14 | 19143 | 1.868889 | 0.000698 | 426 |
| Axl | 26362 | 1.867455 | 0.000504 | 427 |
| H2-Q7 | 15018 | 1.866626 | 0.000001 | 428 |
| Cdk6 | 12571 | 1.866406 | 0.001797 | 429 |
| Il18bp | 16068 | 1.862901 | 0.000001 | 430 |
| Zc3hav1 | 78781 | 1.862117 | 0.000959 | 431 |
| Tmem140 | 68487 | 1.861947 | 2.31E-05 | 432 |
| Hck | 15162 | 1.861856 | 6.39E-05 | 433 |
| Cd3e | 12501 | 1.859072 | 0.004795 | 434 |
| Trim56 | 384309 | 1.858604 | 0.000001 | 435 |
| Adgre1 | 13733 | 1.854831 | 0.000143 | 436 |
| H2-Eb1 | 14969 | 1.85392 | 0.001064 | 437 |
| Hmha1 | 70719 | 1.851596 | 3.58E-05 | 438 |
| Tlr13 | 279572 | 1.850842 | 0.000292 | 439 |
| Pros1 | 19128 | 1.850703 | 5.40E-05 | 440 |
| Traf5 | 22033 | 1.850057 | 0.020945 | 441 |
| Neu4 | 241159 | 1.848825 | 0.000345 | 442 |
| F13a1 | 74145 | 1.848762 | 0.0048 | 443 |
| Ckap2 | 80986 | 1.846874 | 0.005494 | 444 |
| Oas2 | 246728 | 1.8431 | 0.00118 | 445 |
| Gpr65 | 14744 | 1.839517 | 0.01896 | 446 |
| Uox | 22262 | 1.837112 | 0.017832 | 447 |
| Ly6i | 57248 | 1.836995 | 0.000706 | 448 |
| Psmb9 | 16912 | 1.834131 | 7.73E-06 | 449 |
| Cd72 | 12517 | 1.831377 | 0.006855 | 450 |
| Plscr2 | 18828 | 1.831017 | 0.001845 | 451 |
| H2-Q6 | 110557 | 1.829999 | 0.000001 | 452 |
| Lpin3 | 64899 | 1.829056 | 0.028616 | 453 |
| Plekhg1 | 213783 | 1.828509 | 0.000001 | 454 |
| Lum | 17022 | 1.828189 | 0.009474 | 455 |
| Ifi44 | 99899 | 1.82172 | 0.017614 | 456 |
| Clec7a | 56644 | 1.817099 | 9.74E-05 | 457 |
| Cd84 | 12523 | 1.813473 | 0.00075 | 458 |
| Ehd2 | 259300 | 1.812908 | 0.000001 | 459 |
| Fes | 14159 | 1.812204 | 0.000961 | 460 |
| Gfap | 14580 | 1.81032 | 0.009343 | 461 |
| Itpr2 | 16439 | 1.808523 | 1.32E-06 | 462 |
| Col4a1 | 12826 | 1.800855 | 0.000001 | 463 |
| Ifi30 | 65972 | 1.800631 | 1.12E-05 | 464 |
| Cd48 | 12506 | 1.800425 | 0.015545 | 465 |
| Fgr | 14191 | 1.794691 | 0.01368 | 466 |
| Trpv4 | 63873 | 1.793179 | 0.02206 | 467 |
| Folr1 | 14275 | 1.7929 | 0.047897 | 468 |

| | | | | |
|---|---|---|---|---|
| Nckap1l | 105855 | 1.789437 | 1.21E-05 | 469 |
| Lrrk1 | 233328 | 1.789359 | 0.000001 | 470 |
| Apol9b | 71898 | 1.789168 | 0.000271 | 471 |
| Apod | 11815 | 1.787072 | 0.000345 | 472 |
| Fam26f | 215900 | 1.786328 | 0.007871 | 473 |
| H2-Oa | 15001 | 1.783807 | 0.015071 | 474 |
| Gm11131 | 1.03E+08 | 1.78109 | 0.016327 | 475 |
| C3ar1 | 12267 | 1.780134 | 3.32E-05 | 476 |
| Cntrl | 26920 | 1.776449 | 0.000001 | 477 |
| Lamc3 | 23928 | 1.77311 | 0.000953 | 478 |
| Postn | 50706 | 1.770448 | 0.000001 | 479 |
| Uaca | 72565 | 1.7693 | 0.000596 | 480 |
| Fam129a | 63913 | 1.767707 | 0.000001 | 481 |
| Bub1b | 12236 | 1.767663 | 0.001595 | 482 |
| Tap2 | 21355 | 1.763695 | 0.005419 | 483 |
| Frrs1 | 20321 | 1.762899 | 0.049587 | 484 |
| Zfp36 | 22695 | 1.762859 | 5.80E-06 | 485 |
| Trim34a | 94094 | 1.762738 | 0.00011 | 486 |
| Phf11b | 236451 | 1.762688 | 0.001032 | 487 |
| Ctss | 13040 | 1.762055 | 0.032682 | 488 |
| Rtkn2 | 170799 | 1.75943 | 0.000001 | 489 |
| Phldb2 | 208177 | 1.758625 | 0.00013 | 490 |
| Arpc1b | 11867 | 1.758415 | 0.021603 | 491 |
| Cxcl10 | 15945 | 1.756056 | 0.014597 | 492 |
| Serpina3n | 20716 | 1.75289 | 0.00329 | 493 |
| Hpse | 15442 | 1.750767 | 7.46E-05 | 494 |
| Ikzf1 | 22778 | 1.749813 | 0.000001 | 495 |
| Myo1f | 17916 | 1.749366 | 8.55E-06 | 496 |
| Cyp4f18 | 72054 | 1.748263 | 0.022468 | 497 |
| Col1a2 | 12843 | 1.747201 | 1.55E-06 | 498 |
| Lef1 | 16842 | 1.747033 | 0.026183 | 499 |
| Tcirg1 | 27060 | 1.746611 | 0.000001 | 500 |
| Psmb8 | 16913 | 1.745973 | 0.000001 | 501 |
| Fcgr1 | 14129 | 1.742807 | 0.000001 | 502 |
| Lyn | 17096 | 1.741587 | 0.000001 | 503 |
| Csf2rb | 12983 | 1.73895 | 1.67E-05 | 504 |
| Prr11 | 270906 | 1.737596 | 0.018034 | 505 |
| Igha | 238447 | 1.737203 | 7.24E-05 | 506 |
| Adora3 | 11542 | 1.736197 | 0.011101 | 507 |
| Mmrn2 | 105450 | 1.736005 | 0.035274 | 508 |
| Ddx58 | 230073 | 1.727304 | 0.000989 | 509 |
| Cytip | 227929 | 1.726438 | 0.002096 | 510 |
| Trim30d | 209387 | 1.72577 | 0.000001 | 511 |
| Irf5 | 27056 | 1.725048 | 0.000359 | 512 |
| Espl1 | 105988 | 1.72391 | 0.000898 | 513 |
| Nkg7 | 72310 | 1.722244 | 0.008761 | 514 |
| Nfe2l2 | 18024 | 1.722113 | 4.49E-06 | 515 |

| | | | | |
|---|---|---|---|---|
| Gpr179 | 217143 | 1.719777 | 0.007858 | 516 |
| Myrf | 225908 | 1.718602 | 0.000001 | 517 |
| Mag | 17136 | 1.716042 | 0.000001 | 518 |
| Frmd4b | 232288 | 1.71452 | 0.000001 | 519 |
| Adamts1 | 11504 | 1.714505 | 0.000001 | 520 |
| Kif20a | 19348 | 1.713931 | 0.007624 | 521 |
| Stard9 | 668880 | 1.711994 | 9.15E-06 | 522 |
| Iglc2 | 110786 | 1.710744 | 0.033693 | 523 |
| Gpr84 | 80910 | 1.710556 | 0.004422 | 524 |
| Lsp1 | 16985 | 1.709028 | 1.92E-05 | 525 |
| Ptafr | 19204 | 1.703589 | 7.42E-05 | 526 |
| Trim47 | 217333 | 1.702495 | 0.003619 | 527 |
| Oas1a | 246730 | 1.701688 | 3.20E-05 | 528 |
| Gbgt1 | 227671 | 1.698902 | 0.024052 | 529 |
| Emp1 | 13730 | 1.698262 | 0.006218 | 530 |
| Tspo | 12257 | 1.697029 | 4.21E-05 | 531 |
| Slamf8 | 74748 | 1.693453 | 0.001375 | 532 |
| Tjp3 | 27375 | 1.692727 | 0.011509 | 533 |
| Plin2 | 11520 | 1.687275 | 0.000149 | 534 |
| Hmmr | 15366 | 1.687076 | 0.00414 | 535 |
| Fxyd5 | 18301 | 1.685352 | 0.002884 | 536 |
| Trim12a | 76681 | 1.679982 | 9.52E-05 | 537 |
| Clec2d | 93694 | 1.678338 | 4.10E-05 | 538 |
| Ms4a6d | 68774 | 1.676428 | 0.013018 | 539 |
| Stab1 | 192187 | 1.676419 | 0.000001 | 540 |
| Neurl3 | 214854 | 1.67626 | 0.000425 | 541 |
| Adamtsl4 | 229595 | 1.675482 | 0.000001 | 542 |
| Npr1 | 18160 | 1.675339 | 0.040618 | 543 |
| Enpp2 | 18606 | 1.674771 | 0.000001 | 544 |
| Grn | 14824 | 1.673889 | 0.041068 | 545 |
| Lpcat2 | 270084 | 1.673384 | 0.000527 | 546 |
| Cd93 | 17064 | 1.673028 | 0.000001 | 547 |
| Lpxn | 107321 | 1.671243 | 0.005163 | 548 |
| Tjp2 | 21873 | 1.670717 | 0.000001 | 549 |
| Ccnb1 | 268697 | 1.670278 | 0.018668 | 550 |
| Tor4a | 227612 | 1.669493 | 0.001426 | 551 |
| Herc6 | 67138 | 1.669158 | 0.000001 | 552 |
| Ap1g2 | 11766 | 1.667788 | 0.011244 | 553 |
| Col5a1 | 12831 | 1.665025 | 0.008578 | 554 |
| Pdgfra | 18595 | 1.664794 | 0.006558 | 555 |
| Ly6a | 110454 | 1.664301 | 0.000001 | 556 |
| Nes | 18008 | 1.663463 | 0.000458 | 557 |
| Gata2 | 14461 | 1.662373 | 0.040502 | 558 |
| Gimap8 | 243374 | 1.659409 | 0.011889 | 559 |
| Btk | 12229 | 1.658736 | 0.022007 | 560 |
| Pou2af1 | 18985 | 1.65861 | 0.027933 | 561 |
| Emilin1 | 100952 | 1.658424 | 0.000001 | 562 |

| | | | | |
|---|---|---|---|---|
| Isg20 | 57444 | 1.65717 | 0.004839 | 563 |
| Ebf3 | 13593 | 1.65692 | 0.000001 | 564 |
| Loxl3 | 16950 | 1.656807 | 0.00503 | 565 |
| Palld | 72333 | 1.654465 | 4.51E-05 | 566 |
| Glipr2 | 384009 | 1.653909 | 4.54E-05 | 567 |
| Serpina3g | 20715 | 1.653799 | 8.46E-05 | 568 |
| Dok1 | 13448 | 1.65373 | 0.006823 | 569 |
| Ppp1r3b | 244416 | 1.6527 | 0.004641 | 570 |
| Fam46c | 74645 | 1.652191 | 0.000001 | 571 |
| Capg | 12332 | 1.650816 | 0.00089 | 572 |
| Mmp14 | 17387 | 1.649314 | 0.000001 | 573 |
| Nlrp3 | 216799 | 1.643231 | 0.016529 | 574 |
| Gsn | 227753 | 1.64255 | 0.000001 | 575 |
| Traf3ip3 | 215243 | 1.642547 | 0.008074 | 576 |
| Galnt6 | 207839 | 1.641793 | 0.000001 | 577 |
| Hist1h3b | 319150 | 1.640806 | 0.003393 | 578 |
| Trim14 | 74735 | 1.637886 | 0.003329 | 579 |
| Ifit3b | 667370 | 1.637322 | 0.025682 | 580 |
| Aspg | 104816 | 1.635848 | 0.010274 | 581 |
| Samd9l | 209086 | 1.632915 | 0.000001 | 582 |
| Ikbke | 56489 | 1.632695 | 0.030152 | 583 |
| 4632428N( | 74048 | 1.629855 | 2.13E-05 | 584 |
| Efemp1 | 216616 | 1.629267 | 0.000134 | 585 |
| Ncf4 | 17972 | 1.628721 | 0.015598 | 586 |
| Afap1l2 | 226250 | 1.625565 | 3.14E-05 | 587 |
| Ccr5 | 12774 | 1.625516 | 5.99E-05 | 588 |
| Dna2 | 327762 | 1.624604 | 0.009359 | 589 |
| Eng | 13805 | 1.624494 | 0.000001 | 590 |
| G530011O( | 654820 | 1.620936 | 2.23E-05 | 591 |
| Arhgap9 | 216445 | 1.619185 | 0.047292 | 592 |
| Slc1a5 | 20514 | 1.619072 | 0.007288 | 593 |
| Smtn | 29856 | 1.61675 | 1.09E-06 | 594 |
| Tmem63a | 208795 | 1.614421 | 0.000001 | 595 |
| Otx1 | 18423 | 1.61409 | 0.000338 | 596 |
| Lect1 | 16840 | 1.612537 | 0.00437 | 597 |
| Psme2b | 621823 | 1.612055 | 0.023177 | 598 |
| Parp10 | 671535 | 1.609339 | 0.000001 | 599 |
| Itga5 | 16402 | 1.609247 | 0.001004 | 600 |
| Lgals3 | 16854 | 1.609052 | 0.001297 | 601 |
| Unc45b | 217012 | 1.608937 | 0.010515 | 602 |
| Vsig10 | 231668 | 1.607887 | 0.032922 | 603 |
| Txnip | 56338 | 1.605975 | 0.000001 | 604 |
| Mcam | 84004 | 1.605464 | 1.35E-05 | 605 |
| Renbp | 19703 | 1.604576 | 0.032383 | 606 |
| Erbb3 | 13867 | 1.604546 | 0.000001 | 607 |
| Abcb1a | 18671 | 1.599708 | 1.20E-05 | 608 |
| Eomes | 13813 | 1.597474 | 0.000001 | 609 |

| | | | | |
|---|---|---|---|---|
| Cyr61 | 16007 | 1.596639 | 0.000478 | 610 |
| Pdpn | 14726 | 1.596592 | 8.58E-06 | 611 |
| Epha2 | 13836 | 1.592963 | 0.002691 | 612 |
| Ncaph | 215387 | 1.591878 | 0.047514 | 613 |
| Il10ra | 16154 | 1.589388 | 4.47E-05 | 614 |
| Stat2 | 20847 | 1.589034 | 0.000515 | 615 |
| Slc11a1 | 18173 | 1.588112 | 3.47E-05 | 616 |
| Spef2 | 320277 | 1.583575 | 0.032166 | 617 |
| Ighv5-6 | 777780 | 1.58271 | 4.21E-06 | 618 |
| Stat4 | 20849 | 1.581935 | 0.002104 | 619 |
| Prkd2 | 101540 | 1.580945 | 0.002619 | 620 |
| Mlph | 171531 | 1.580035 | 0.002699 | 621 |
| Cebpd | 12609 | 1.577809 | 0.000223 | 622 |
| Plau | 18792 | 1.576495 | 0.004277 | 623 |
| Ifi35 | 70110 | 1.575363 | 6.26E-05 | 624 |
| Ets1 | 23871 | 1.573952 | 0.000001 | 625 |
| Ccdc170 | 1.01E+08 | 1.572974 | 9.54E-06 | 626 |
| Stat1 | 20846 | 1.567761 | 0.004063 | 627 |
| Car5b | 56078 | 1.567415 | 0.00564 | 628 |
| Erbb2 | 13866 | 1.562952 | 0.036104 | 629 |
| Fhod1 | 234686 | 1.560594 | 0.002478 | 630 |
| Cxcl13 | 55985 | 1.559923 | 5.54E-05 | 631 |
| Ncf1 | 17969 | 1.559518 | 2.13E-05 | 632 |
| Gpr35 | 64095 | 1.559116 | 0.004733 | 633 |
| Cd37 | 12493 | 1.555365 | 0.001294 | 634 |
| Runx1 | 12394 | 1.55141 | 0.000662 | 635 |
| Enpp6 | 320981 | 1.550877 | 1.43E-06 | 636 |
| Slc15a3 | 65221 | 1.550418 | 0.003564 | 637 |
| Pld4 | 104759 | 1.548809 | 0.000807 | 638 |
| Cmah | 12763 | 1.547297 | 0.031841 | 639 |
| Plekhh1 | 211945 | 1.546127 | 0.000001 | 640 |
| Srgn | 19073 | 1.544592 | 0.010319 | 641 |
| Ahnak | 66395 | 1.543758 | 0.000001 | 642 |
| Havcr2 | 171285 | 1.540805 | 0.00255 | 643 |
| Ifit2 | 15958 | 1.540038 | 0.000496 | 644 |
| Col11a1 | 12814 | 1.537836 | 0.00769 | 645 |
| Sphk1 | 20698 | 1.537797 | 0.043795 | 646 |
| Plscr1 | 22038 | 1.533155 | 0.00853 | 647 |
| Esyt1 | 23943 | 1.532335 | 0.000162 | 648 |
| Cnn3 | 71994 | 1.532305 | 0.008819 | 649 |
| Rnf135 | 71956 | 1.527712 | 0.00316 | 650 |
| H2-T23 | 15040 | 1.524818 | 0.0048 | 651 |
| Igsf6 | 80719 | 1.523412 | 0.037702 | 652 |
| Laptm5 | 16792 | 1.522681 | 2.07E-06 | 653 |
| Plod1 | 18822 | 1.522658 | 0.000001 | 654 |
| Tmem72 | 319776 | 1.521556 | 0.019825 | 655 |
| Slc7a2 | 11988 | 1.520138 | 1.64E-06 | 656 |

| | | | | |
|---|---|---|---|---|
| Adamts4 | 240913 | 1.519908 | 0.000001 | 657 |
| Sned1 | 208777 | 1.519011 | 9.88E-05 | 658 |
| Cpq | 54381 | 1.518437 | 0.000227 | 659 |
| Cpt1a | 12894 | 1.518292 | 2.76E-05 | 660 |
| Aim2 | 383619 | 1.518222 | 0.002634 | 661 |
| Fmo5 | 14263 | 1.518055 | 0.005836 | 662 |
| Kif20b | 240641 | 1.517333 | 0.036104 | 663 |
| Kdr | 16542 | 1.514463 | 0.00097 | 664 |
| Lama5 | 16776 | 1.514009 | 0.000001 | 665 |
| Pik3cg | 30955 | 1.511558 | 0.000258 | 666 |
| Megf10 | 70417 | 1.511491 | 0.000001 | 667 |
| Sox10 | 20665 | 1.510809 | 0.000001 | 668 |
| H2-D1 | 14964 | 1.510629 | 0.005124 | 669 |
| Prc1 | 233406 | 1.506404 | 0.020412 | 670 |
| AW112010 | 107350 | 1.505874 | 0.000559 | 671 |
| Pim1 | 18712 | 1.504729 | 0.000001 | 672 |
| 9530082P2 | 638247 | 1.504716 | 0.013105 | 673 |
| Ptpn22 | 19260 | 1.503425 | 0.006462 | 674 |
| Myof | 226101 | 1.503172 | 0.000001 | 675 |
| Fli1 | 14247 | 1.502609 | 0.000158 | 676 |
| Ggta1 | 14594 | 1.50189 | 0.001566 | 677 |
| Nipal4 | 214112 | 1.500822 | 0.000423 | 678 |
| Anxa3 | 11745 | 1.499183 | 0.003822 | 679 |
| Zfp217 | 228913 | 1.497413 | 0.000107 | 680 |
| Atp7b | 11979 | 1.497018 | 0.000434 | 681 |
| Cd44 | 12505 | 1.495792 | 1.61E-06 | 682 |
| Snx33 | 235406 | 1.495395 | 0.000001 | 683 |
| Fam111a | 107373 | 1.494237 | 0.000952 | 684 |
| Cep192 | 70799 | 1.492381 | 1.62E-05 | 685 |
| Col5a2 | 12832 | 1.489752 | 0.02677 | 686 |
| Lhx1 | 16869 | 1.489181 | 0.000572 | 687 |
| Wipf1 | 215280 | 1.488823 | 0.000001 | 688 |
| C5ar1 | 12273 | 1.488561 | 0.040907 | 689 |
| Rasa4 | 54153 | 1.484748 | 5.24E-06 | 690 |
| C1qa | 12259 | 1.483598 | 0.034915 | 691 |
| Syne3 | 212073 | 1.483156 | 0.000786 | 692 |
| Irf7 | 54123 | 1.482763 | 0.000001 | 693 |
| Gna15 | 14676 | 1.481831 | 0.044917 | 694 |
| Traf1 | 22029 | 1.481074 | 0.00134 | 695 |
| Susd5 | 382111 | 1.48107 | 0.001895 | 696 |
| Dnase2a | 13423 | 1.475562 | 0.02237 | 697 |
| Kif11 | 16551 | 1.472754 | 0.001657 | 698 |
| Stat5a | 20850 | 1.47229 | 0.001809 | 699 |
| Lcp2 | 16822 | 1.470674 | 0.000253 | 700 |
| 5730559C1 | 67313 | 1.468358 | 0.019086 | 701 |
| Klf4 | 16600 | 1.467769 | 0.008705 | 702 |
| Dsp | 109620 | 1.466977 | 2.39E-06 | 703 |

| | | | | |
|---|---|---|---|---|
| Rhbdf1 | 13650 | 1.466479 | 0.002364 | 704 |
| Tfpi | 21788 | 1.462428 | 0.029842 | 705 |
| Vcan | 13003 | 1.461457 | 2.96E-06 | 706 |
| Creb5 | 231991 | 1.459839 | 0.000122 | 707 |
| Itih5 | 209378 | 1.457639 | 0.000001 | 708 |
| Spi1 | 20375 | 1.454884 | 0.00036 | 709 |
| 5031414D1 | 271221 | 1.454766 | 0.046127 | 710 |
| Piezo1 | 234839 | 1.454586 | 0.000001 | 711 |
| Syk | 20963 | 1.451389 | 8.54E-05 | 712 |
| Mcm6 | 17219 | 1.451171 | 1.34E-05 | 713 |
| Ckap2l | 70466 | 1.450428 | 0.049104 | 714 |
| Prr5l | 72446 | 1.449754 | 0.000001 | 715 |
| Irak4 | 266632 | 1.447975 | 0.019111 | 716 |
| Emid1 | 140703 | 1.447082 | 0.012947 | 717 |
| Pnpla7 | 241274 | 1.447048 | 1.29E-05 | 718 |
| Pml | 18854 | 1.445869 | 0.000001 | 719 |
| Ccdc88b | 78317 | 1.444501 | 9.53E-05 | 720 |
| P2ry13 | 74191 | 1.442734 | 0.002114 | 721 |
| Thbd | 21824 | 1.441881 | 0.03456 | 722 |
| Prkcq | 18761 | 1.440457 | 0.000001 | 723 |
| Itga1 | 109700 | 1.437019 | 0.02206 | 724 |
| Gm6548 | 625054 | 1.436894 | 0.001178 | 725 |
| Lyl1 | 17095 | 1.434728 | 0.017529 | 726 |
| Maml2 | 270118 | 1.430722 | 0.000001 | 727 |
| Col8a2 | 329941 | 1.42634 | 0.005598 | 728 |
| Gab1 | 14388 | 1.425066 | 0.000001 | 729 |
| Nkx6-2 | 14912 | 1.42505 | 1.04E-05 | 730 |
| Unc93b1 | 54445 | 1.424229 | 0.000143 | 731 |
| Ebf2 | 13592 | 1.424022 | 0.000001 | 732 |
| Cxcl16 | 66102 | 1.42148 | 0.004818 | 733 |
| Prex2 | 109294 | 1.421376 | 0.014433 | 734 |
| Caskin2 | 140721 | 1.421156 | 1.38E-05 | 735 |
| Angptl2 | 26360 | 1.421127 | 0.00251 | 736 |
| Thbs1 | 21825 | 1.420679 | 0.000001 | 737 |
| Hs3st3b1 | 54710 | 1.420524 | 0.007883 | 738 |
| S1pr3 | 13610 | 1.419981 | 1.38E-05 | 739 |
| Eif2ak2 | 19106 | 1.418942 | 0.000001 | 740 |
| Svep1 | 64817 | 1.418921 | 0.043754 | 741 |
| Ctsh | 13036 | 1.418043 | 0.000119 | 742 |
| Icosl | 50723 | 1.417978 | 0.002798 | 743 |
| Apol9a | 223672 | 1.413753 | 0.000349 | 744 |
| Cd9 | 12527 | 1.412776 | 3.82E-05 | 745 |
| Itpr3 | 16440 | 1.412706 | 0.00041 | 746 |
| Slc29a3 | 71279 | 1.40711 | 3.99E-05 | 747 |
| Tlr3 | 142980 | 1.406873 | 0.000147 | 748 |
| Znfx1 | 98999 | 1.406559 | 0.001906 | 749 |
| Aldh1a1 | 11668 | 1.403451 | 3.05E-05 | 750 |

| | | | | |
|---|---|---|---|---|
| Tmco4 | 77056 | 1.403108 | 0.000133 | 751 |
| Hspa1a | 193740 | 1.401919 | 0.000001 | 752 |
| Klf2 | 16598 | 1.399034 | 0.002902 | 753 |
| Tnfsf8 | 21949 | 1.397176 | 0.044582 | 754 |
| Heph | 15203 | 1.395011 | 0.000905 | 755 |
| Il21r | 60504 | 1.394846 | 0.013082 | 756 |
| Tns3 | 319939 | 1.393342 | 0.035656 | 757 |
| Svil | 225115 | 1.392621 | 3.22E-06 | 758 |
| Pax5 | 18507 | 1.391194 | 0.047576 | 759 |
| Ptprz1 | 19283 | 1.39094 | 0.014662 | 760 |
| Cd33 | 12489 | 1.390936 | 0.004127 | 761 |
| Arhgap17 | 70497 | 1.389416 | 6.19E-05 | 762 |
| Bin2 | 668218 | 1.38927 | 0.000304 | 763 |
| Csf1 | 12977 | 1.38848 | 0.000001 | 764 |
| Oas1g | 23960 | 1.384972 | 0.026747 | 765 |
| Pnp | 18950 | 1.38495 | 3.50E-05 | 766 |
| Fzd7 | 14369 | 1.384105 | 0.000001 | 767 |
| Ttf2 | 74044 | 1.379417 | 0.005592 | 768 |
| Ripk1 | 19766 | 1.379049 | 0.000236 | 769 |
| Corin | 53419 | 1.378554 | 3.82E-05 | 770 |
| Oasl2 | 23962 | 1.376867 | 0.012628 | 771 |
| Pask | 269224 | 1.375571 | 0.010661 | 772 |
| Phldb1 | 102693 | 1.374782 | 0.002713 | 773 |
| Mx2 | 17858 | 1.374768 | 0.000001 | 774 |
| Tpx2 | 72119 | 1.371879 | 0.003535 | 775 |
| Kank2 | 235041 | 1.369987 | 0.0001 | 776 |
| Atp13a5 | 268878 | 1.369856 | 0.031174 | 777 |
| Rbpms | 19663 | 1.369833 | 0.016683 | 778 |
| Tnfaip3 | 21929 | 1.369685 | 1.92E-05 | 779 |
| Stk10 | 20868 | 1.369356 | 0.000599 | 780 |
| Ldlrap1 | 100017 | 1.369318 | 0.025116 | 781 |
| Slco2a1 | 24059 | 1.368622 | 0.042142 | 782 |
| Hspa1b | 15511 | 1.368304 | 0.000001 | 783 |
| Tlr7 | 170743 | 1.367537 | 0.010843 | 784 |
| H2-DMb1 | 14999 | 1.364313 | 0.000352 | 785 |
| Rin3 | 217835 | 1.363576 | 0.005263 | 786 |
| Cnn2 | 12798 | 1.36342 | 0.000325 | 787 |
| Nfam1 | 74039 | 1.362657 | 0.004635 | 788 |
| Tcn2 | 21452 | 1.361171 | 0.000642 | 789 |
| Sema3b | 20347 | 1.360977 | 0.003016 | 790 |
| Aqp4 | 11829 | 1.358135 | 0.010128 | 791 |
| Chst3 | 53374 | 1.356158 | 0.001316 | 792 |
| Mrvi1 | 17540 | 1.35544 | 0.001698 | 793 |
| Stxbp3 | 20912 | 1.353522 | 2.04E-05 | 794 |
| Iqgap1 | 29875 | 1.353063 | 0.000001 | 795 |
| C1qc | 12262 | 1.352306 | 0.03477 | 796 |
| Rac2 | 19354 | 1.351528 | 0.00048 | 797 |

| | | | | |
|---|---|---|---|---|
| Micall2 | 231830 | 1.348701 | 0.005681 | 798 |
| Cldn11 | 18417 | 1.347339 | 0.000001 | 799 |
| Iqgap3 | 404710 | 1.34549 | 0.014706 | 800 |
| Fcer1g | 14127 | 1.345189 | 0.00022 | 801 |
| Mmp2 | 17390 | 1.345186 | 0.009183 | 802 |
| Arap3 | 106952 | 1.343657 | 0.015509 | 803 |
| Csf3r | 12986 | 1.342605 | 0.001502 | 804 |
| Slco1c1 | 58807 | 1.340856 | 0.000001 | 805 |
| B4galt1 | 14595 | 1.33956 | 7.75E-06 | 806 |
| Prox1 | 19130 | 1.333464 | 0.000001 | 807 |
| Col12a1 | 12816 | 1.333226 | 1.25E-05 | 808 |
| Bmp2k | 140780 | 1.33313 | 0.000001 | 809 |
| Sema3d | 108151 | 1.331799 | 0.001111 | 810 |
| Grap | 71520 | 1.331352 | 0.04447 | 811 |
| Tgfb1 | 21803 | 1.331265 | 0.002089 | 812 |
| Ebf1 | 13591 | 1.327122 | 0.005858 | 813 |
| Aass | 30956 | 1.325236 | 0.017531 | 814 |
| Cdh19 | 227485 | 1.323931 | 3.16E-05 | 815 |
| Mboat1 | 218121 | 1.323097 | 0.024761 | 816 |
| Ly86 | 17084 | 1.322713 | 0.003373 | 817 |
| Abca1 | 11303 | 1.322351 | 1.16E-06 | 818 |
| Ctnna1 | 12385 | 1.321771 | 0.000001 | 819 |
| Mov10 | 17454 | 1.321644 | 0.000001 | 820 |
| Ifitm3 | 66141 | 1.321326 | 5.97E-05 | 821 |
| Slc12a4 | 20498 | 1.319176 | 1.27E-06 | 822 |
| Sh3bp2 | 24055 | 1.3185 | 0.00012 | 823 |
| Plekhg3 | 263406 | 1.317356 | 0.000001 | 824 |
| Slc38a3 | 76257 | 1.31697 | 0.043582 | 825 |
| Prkd3 | 75292 | 1.314619 | 0.000001 | 826 |
| Pycard | 66824 | 1.312726 | 0.017179 | 827 |
| Myo6 | 17920 | 1.311771 | 1.33E-06 | 828 |
| Cand2 | 67088 | 1.311065 | 0.000554 | 829 |
| Sh3tc2 | 225608 | 1.310734 | 0.02686 | 830 |
| Phactr4 | 100169 | 1.310631 | 0.000115 | 831 |
| Tgif1 | 21815 | 1.309969 | 0.020733 | 832 |
| Tnfrsf1a | 21937 | 1.309065 | 3.95E-05 | 833 |
| Col1a1 | 12842 | 1.308815 | 0.001053 | 834 |
| Map4k1 | 26411 | 1.306348 | 0.012675 | 835 |
| Ptges | 64292 | 1.305793 | 0.012796 | 836 |
| Tfcp2l1 | 81879 | 1.304053 | 0.000231 | 837 |
| Cyba | 13057 | 1.302657 | 0.006364 | 838 |
| Glis3 | 226075 | 1.302002 | 0.000001 | 839 |
| Tep1 | 21745 | 1.301699 | 8.52E-05 | 840 |
| Tns2 | 209039 | 1.301672 | 0.000001 | 841 |
| Usp18 | 24110 | 1.301078 | 0.000239 | 842 |
| Csrnp1 | 215418 | 1.300981 | 2.70E-06 | 843 |
| Ermn | 77767 | 1.295781 | 0.000001 | 844 |

| | | | | |
|---|---|---|---|---|
| Pbxip1 | 229534 | 1.294923 | 5.43E-05 | 845 |
| Abca4 | 11304 | 1.294766 | 0.000734 | 846 |
| Rbl1 | 19650 | 1.294611 | 0.013314 | 847 |
| Anxa2 | 12306 | 1.291997 | 2.89E-06 | 848 |
| Nfkbie | 18037 | 1.290396 | 0.003909 | 849 |
| Cybrd1 | 73649 | 1.288647 | 0.006264 | 850 |
| Timeless | 21853 | 1.285824 | 1.41E-06 | 851 |
| Cd53 | 12508 | 1.284297 | 0.001349 | 852 |
| Rreb1 | 68750 | 1.282625 | 0.000001 | 853 |
| Slc8b1 | 170756 | 1.281332 | 0.002107 | 854 |
| Plcb3 | 18797 | 1.280749 | 0.006393 | 855 |
| Dock1 | 330662 | 1.279865 | 0.000001 | 856 |
| Fzd6 | 14368 | 1.278706 | 0.00805 | 857 |
| Slc12a7 | 20499 | 1.278704 | 0.000161 | 858 |
| Lpp | 210126 | 1.277391 | 3.58E-05 | 859 |
| P2rx7 | 18439 | 1.2772 | 0.00041 | 860 |
| Padi2 | 18600 | 1.276015 | 0.000001 | 861 |
| Cav1 | 12389 | 1.275998 | 0.001595 | 862 |
| Irf4 | 16364 | 1.274118 | 0.013055 | 863 |
| Cd52 | 23833 | 1.273836 | 0.002794 | 864 |
| Itk | 16428 | 1.272912 | 0.017284 | 865 |
| Synpo2 | 118449 | 1.272884 | 0.002724 | 866 |
| Olfml3 | 99543 | 1.269201 | 0.00041 | 867 |
| Ugt8a | 22239 | 1.265641 | 0.000001 | 868 |
| Arid5a | 214855 | 1.263686 | 4.10E-05 | 869 |
| Birc3 | 11796 | 1.261842 | 0.00144 | 870 |
| Entpd1 | 12495 | 1.258262 | 1.91E-06 | 871 |
| Zfp36l2 | 12193 | 1.256698 | 0.000001 | 872 |
| Mtbp | 105837 | 1.256316 | 0.03104 | 873 |
| Myh9 | 17886 | 1.256023 | 0.019257 | 874 |
| Gsap | 212167 | 1.254766 | 0.000391 | 875 |
| Clic1 | 114584 | 1.253328 | 0.003264 | 876 |
| Plekhf1 | 72287 | 1.252477 | 0.038607 | 877 |
| Stk17b | 98267 | 1.252441 | 0.001269 | 878 |
| Zfp36l1 | 12192 | 1.251988 | 0.000001 | 879 |
| 1700017B0 | 74211 | 1.250092 | 1.23E-05 | 880 |
| Syngr2 | 20973 | 1.247229 | 2.60E-05 | 881 |
| Abcc9 | 20928 | 1.246207 | 0.012526 | 882 |
| Uncx | 22255 | 1.244821 | 0.027852 | 883 |
| Tifa | 211550 | 1.243101 | 0.009354 | 884 |
| Hvcn1 | 74096 | 1.241779 | 0.001552 | 885 |
| Amot | 27494 | 1.239825 | 0.000001 | 886 |
| Acsf2 | 264895 | 1.239761 | 0.006336 | 887 |
| Itpripl2 | 319622 | 1.236663 | 0.000105 | 888 |
| Mab21l1 | 17116 | 1.236044 | 0.038133 | 889 |
| Selplg | 20345 | 1.235564 | 0.002092 | 890 |
| Socs1 | 12703 | 1.235097 | 0.000118 | 891 |

| | | | | |
|---|---|---|---|---|
| Vwf | 22371 | 1.233479 | 0.023428 | 892 |
| Plin3 | 66905 | 1.2312 | 4.55E-05 | 893 |
| Aldh1l2 | 216188 | 1.230219 | 0.002358 | 894 |
| Ighv1-61 | 674018 | 1.229409 | 0.000569 | 895 |
| Daam2 | 76441 | 1.229174 | 0.000001 | 896 |
| Tgfbr2 | 21813 | 1.228785 | 7.73E-06 | 897 |
| Akap13 | 75547 | 1.226411 | 0.000001 | 898 |
| Tmem119 | 231633 | 1.225331 | 0.001269 | 899 |
| Pdgfrb | 18596 | 1.225247 | 0.000377 | 900 |
| Anln | 68743 | 1.225099 | 0.000001 | 901 |
| Sbno2 | 216161 | 1.223491 | 0.000001 | 902 |
| Adamtsl1 | 77739 | 1.220502 | 0.000001 | 903 |
| Rrbp1 | 81910 | 1.22004 | 6.79E-05 | 904 |
| Aif1 | 11629 | 1.21986 | 0.042289 | 905 |
| Gli2 | 14633 | 1.218599 | 0.018027 | 906 |
| Slco2b1 | 101488 | 1.218209 | 0.004718 | 907 |
| Crispld2 | 78892 | 1.21795 | 0.023399 | 908 |
| Epsti1 | 108670 | 1.217776 | 0.001816 | 909 |
| Cyp1b1 | 13078 | 1.217347 | 0.000284 | 910 |
| Kank1 | 107351 | 1.216606 | 3.46E-05 | 911 |
| Slc39a12 | 277468 | 1.213845 | 0.001651 | 912 |
| Sox9 | 20682 | 1.210857 | 1.07E-05 | 913 |
| Ezr | 22350 | 1.210524 | 0.000001 | 914 |
| Myo1d | 338367 | 1.210165 | 0.000001 | 915 |
| Efemp2 | 58859 | 1.209747 | 0.00692 | 916 |
| Ptrf | 19285 | 1.208956 | 1.89E-06 | 917 |
| Rest | 19712 | 1.208222 | 0.000001 | 918 |
| Stat3 | 20848 | 1.208023 | 0.000001 | 919 |
| Scarna10 | 1E+08 | 1.207005 | 0.006225 | 920 |
| Gli3 | 14634 | 1.206612 | 0.000961 | 921 |
| Frmd8 | 67457 | 1.206011 | 4.17E-06 | 922 |
| Maff | 17133 | 1.204428 | 0.007545 | 923 |
| Cpt2 | 12896 | 1.20413 | 0.000706 | 924 |
| Shc1 | 20416 | 1.203651 | 0.000342 | 925 |
| Cd82 | 12521 | 1.203451 | 1.91E-05 | 926 |
| Ifit1 | 15957 | 1.202573 | 0.02277 | 927 |
| Arap1 | 69710 | 1.201784 | 6.42E-05 | 928 |
| Wdr62 | 233064 | 1.199184 | 0.011196 | 929 |
| Hmox1 | 15368 | 1.197713 | 7.57E-05 | 930 |
| Bst2 | 69550 | 1.197497 | 0.00088 | 931 |
| Matn4 | 17183 | 1.194397 | 0.030082 | 932 |
| Acvrl1 | 11482 | 1.189322 | 0.032486 | 933 |
| Gem | 14579 | 1.189272 | 0.025647 | 934 |
| Itpripl1 | 73338 | 1.187388 | 0.011155 | 935 |
| Cnp | 12799 | 1.186802 | 0.000001 | 936 |
| Vtn | 22370 | 1.186372 | 8.75E-05 | 937 |
| Wasf2 | 242687 | 1.185542 | 0.000001 | 938 |

| | | | | |
|---|---|---|---|---|
| Arhgef10 | 234094 | 1.182159 | 0.000001 | 939 |
| Tagln2 | 21346 | 1.180613 | 1.43E-06 | 940 |
| Mapkapk3 | 102626 | 1.180309 | 0.01682 | 941 |
| Bag3 | 29810 | 1.179452 | 1.41E-06 | 942 |
| Erap1 | 80898 | 1.17917 | 0.000245 | 943 |
| Ctsc | 13032 | 1.17908 | 4.94E-05 | 944 |
| Myo1e | 71602 | 1.178849 | 0.000001 | 945 |
| Elf1 | 13709 | 1.175923 | 0.000318 | 946 |
| Il6ra | 16194 | 1.175791 | 0.00012 | 947 |
| Prom1 | 19126 | 1.175756 | 0.028822 | 948 |
| Insc | 233752 | 1.175691 | 0.013505 | 949 |
| Slco1a4 | 28250 | 1.175375 | 0.005797 | 950 |
| Mcm5 | 17218 | 1.175083 | 0.001059 | 951 |
| Serpine1 | 18787 | 1.174788 | 0.002788 | 952 |
| Neat1 | 66961 | 1.173922 | 0.000404 | 953 |
| Athl1 | 212974 | 1.173832 | 0.048382 | 954 |
| Spsb1 | 74646 | 1.173434 | 0.000001 | 955 |
| Brca1 | 12189 | 1.172502 | 0.000816 | 956 |
| Cdkn1c | 12577 | 1.171148 | 0.016114 | 957 |
| Ighg2c | 404711 | 1.170945 | 0.010411 | 958 |
| Cpne3 | 70568 | 1.167962 | 8.83E-05 | 959 |
| Slc14a1 | 108052 | 1.167894 | 0.010522 | 960 |
| Med12 | 59024 | 1.165019 | 6.05E-06 | 961 |
| Gm9581 | 672884 | 1.164852 | 0.038595 | 962 |
| Sec14l5 | 665119 | 1.163651 | 0.005379 | 963 |
| Capn3 | 12335 | 1.162665 | 0.004122 | 964 |
| Aff1 | 17355 | 1.162532 | 0.000001 | 965 |
| Fasn | 14104 | 1.162347 | 0.001466 | 966 |
| Entpd2 | 12496 | 1.161876 | 0.049104 | 967 |
| Crybg3 | 224273 | 1.1616 | 0.003634 | 968 |
| Cmklr1 | 14747 | 1.16032 | 0.023574 | 969 |
| Sod3 | 20657 | 1.159707 | 0.010695 | 970 |
| Abi3 | 66610 | 1.159029 | 0.032804 | 971 |
| Mavs | 228607 | 1.15838 | 0.004794 | 972 |
| Ttyh2 | 117160 | 1.156742 | 0.000001 | 973 |
| Atg7 | 74244 | 1.154518 | 0.000001 | 974 |
| Notch3 | 18131 | 1.154486 | 0.001988 | 975 |
| Epas1 | 13819 | 1.149946 | 0.000001 | 976 |
| Adap2 | 216991 | 1.149081 | 0.00757 | 977 |
| Cad | 69719 | 1.146366 | 0.000001 | 978 |
| Rab32 | 67844 | 1.145646 | 0.01974 | 979 |
| Tjp1 | 21872 | 1.145521 | 0.000001 | 980 |
| Ifi27l2a | 76933 | 1.144821 | 0.001075 | 981 |
| Fbln1 | 14114 | 1.144747 | 3.45E-05 | 982 |
| Pld1 | 18805 | 1.144602 | 0.001809 | 983 |
| Col8a1 | 12837 | 1.144573 | 0.002957 | 984 |
| Cldn19 | 242653 | 1.143877 | 0.049021 | 985 |

| | | | | |
|---|---|---|---|---|
| Pon3 | 269823 | 1.140594 | 0.018865 | 986 |
| Hhip | 15245 | 1.140175 | 0.000188 | 987 |
| Hpgds | 54486 | 1.138943 | 0.005117 | 988 |
| Aldh3b1 | 67689 | 1.13759 | 0.006708 | 989 |
| Hacl1 | 56794 | 1.137136 | 0.028702 | 990 |
| Tram2 | 170829 | 1.137016 | 0.008208 | 991 |
| Il33 | 77125 | 1.136936 | 0.000001 | 992 |
| Exoc3l4 | 74190 | 1.136744 | 0.014112 | 993 |
| Igfbp2 | 16008 | 1.134586 | 0.000362 | 994 |
| Gpsm3 | 106512 | 1.133444 | 0.024786 | 995 |
| Stom | 13830 | 1.13145 | 0.010041 | 996 |
| Mical1 | 171580 | 1.130891 | 0.006669 | 997 |
| Kifc1 | 1.01E+08 | 1.130867 | 0.031065 | 998 |
| Steap3 | 68428 | 1.129523 | 0.006675 | 999 |
| Rtp4 | 67775 | 1.1294 | 0.000372 | 1000 |
| Col6a3 | 12835 | 1.127447 | 0.000001 | 1001 |
| Fkbp10 | 14230 | 1.124607 | 0.026826 | 1002 |
| Chd7 | 320790 | 1.122027 | 0.000001 | 1003 |
| Kif13b | 16554 | 1.121946 | 0.000001 | 1004 |
| Tifab | 212937 | 1.120387 | 0.01659 | 1005 |
| Sorbs3 | 20410 | 1.11915 | 4.34E-06 | 1006 |
| Ldlr | 16835 | 1.115497 | 0.000001 | 1007 |
| Jak3 | 16453 | 1.115126 | 0.021444 | 1008 |
| Magt1 | 67075 | 1.114586 | 6.05E-06 | 1009 |
| Ndrg1 | 17988 | 1.114368 | 0.000001 | 1010 |
| Plcd1 | 18799 | 1.114049 | 0.002215 | 1011 |
| Bmf | 171543 | 1.112328 | 0.002575 | 1012 |
| Samhd1 | 56045 | 1.110409 | 1.18E-05 | 1013 |
| Cyp4v3 | 102294 | 1.109606 | 0.016262 | 1014 |
| Pdlim2 | 213019 | 1.108264 | 0.007162 | 1015 |
| Nfatc1 | 18018 | 1.104879 | 0.001139 | 1016 |
| Aspa | 11484 | 1.103497 | 0.000738 | 1017 |
| Mrc2 | 17534 | 1.103045 | 0.013769 | 1018 |
| Nek8 | 140859 | 1.098997 | 0.048974 | 1019 |
| Fyb | 23880 | 1.098902 | 0.002265 | 1020 |
| Lrig3 | 320398 | 1.097331 | 0.01724 | 1021 |
| Epb4.1l2 | 13822 | 1.096801 | 1.51E-05 | 1022 |
| Ltbr | 17000 | 1.096434 | 0.005627 | 1023 |
| Rnf43 | 207742 | 1.093469 | 0.042815 | 1024 |
| Fa2h | 338521 | 1.092464 | 0.000001 | 1025 |
| Gadd45b | 17873 | 1.090588 | 0.007046 | 1026 |
| Ppfibp2 | 19024 | 1.09051 | 0.001551 | 1027 |
| P2ry6 | 233571 | 1.089067 | 0.03382 | 1028 |
| Hr | 15460 | 1.088427 | 0.000001 | 1029 |
| Akna | 100182 | 1.087055 | 0.00014 | 1030 |
| Gjc2 | 118454 | 1.085962 | 1.48E-05 | 1031 |
| Stard13 | 243362 | 1.0857 | 0.000001 | 1032 |

| | | | | |
|---|---|---|---|---|
| Dab2 | 13132 | 1.08497 | 8.85E-05 | 1033 |
| Samd3 | 268288 | 1.080676 | 0.000642 | 1034 |
| Tmbim1 | 69660 | 1.080281 | 2.42E-06 | 1035 |
| Col18a1 | 12822 | 1.079644 | 0.005557 | 1036 |
| Bche | 12038 | 1.079494 | 0.003849 | 1037 |
| Dock5 | 68813 | 1.079431 | 0.000001 | 1038 |
| Olfml1 | 244198 | 1.079135 | 0.006718 | 1039 |
| Flt1 | 14254 | 1.076115 | 0.000816 | 1040 |
| Plp1 | 18823 | 1.074779 | 0.000001 | 1041 |
| Cenpe | 229841 | 1.074459 | 0.03654 | 1042 |
| Fcgrt | 14132 | 1.071994 | 0.003965 | 1043 |
| Cecr2 | 330409 | 1.071382 | 0.005264 | 1044 |
| Sash1 | 70097 | 1.069277 | 0.000001 | 1045 |
| Nupr1 | 56312 | 1.06667 | 0.048382 | 1046 |
| Tmem88b | 320587 | 1.064187 | 0.000001 | 1047 |
| Rhoj | 80837 | 1.063783 | 0.011509 | 1048 |
| Crot | 74114 | 1.062566 | 0.002984 | 1049 |
| Pik3r5 | 320207 | 1.062254 | 0.049938 | 1050 |
| B2m | 12010 | 1.061349 | 0.013815 | 1051 |
| Pygl | 110095 | 1.061031 | 0.017133 | 1052 |
| Ston2 | 108800 | 1.060753 | 0.005681 | 1053 |
| Nuak2 | 74137 | 1.057172 | 0.007767 | 1054 |
| Dcn | 13179 | 1.056827 | 0.008325 | 1055 |
| Fcgr3 | 14131 | 1.056682 | 0.006724 | 1056 |
| Xaf1 | 327959 | 1.056173 | 0.000805 | 1057 |
| Malt1 | 240354 | 1.053006 | 0.017692 | 1058 |
| Lyz2 | 17105 | 1.05004 | 0.001445 | 1059 |
| Lbp | 16803 | 1.049891 | 0.00651 | 1060 |
| Prrx1 | 18933 | 1.048421 | 0.026817 | 1061 |
| Zbtb7b | 22724 | 1.046428 | 0.000001 | 1062 |
| Syde1 | 71709 | 1.046079 | 0.001417 | 1063 |
| Dmd | 13405 | 1.045666 | 3.65E-06 | 1064 |
| Adam17 | 11491 | 1.045382 | 0.000865 | 1065 |
| Mcm2 | 17216 | 1.044677 | 0.002638 | 1066 |
| Triobp | 110253 | 1.044536 | 0.000679 | 1067 |
| Tirap | 117149 | 1.044528 | 0.045511 | 1068 |
| Il17ra | 16172 | 1.043262 | 0.000202 | 1069 |
| Tor3a | 30935 | 1.043246 | 0.001104 | 1070 |
| Gjc3 | 118446 | 1.043238 | 0.000001 | 1071 |
| Loxl1 | 16949 | 1.038485 | 0.02285 | 1072 |
| Fgfrl1 | 116701 | 1.037814 | 0.016011 | 1073 |
| Mertk | 17289 | 1.037178 | 0.014503 | 1074 |
| Adhfe1 | 76187 | 1.03609 | 0.010939 | 1075 |
| Tfeb | 21425 | 1.031962 | 0.008915 | 1076 |
| Pde8a | 18584 | 1.029466 | 0.000191 | 1077 |
| Slc9a3r1 | 26941 | 1.028448 | 0.000271 | 1078 |
| Dennd3 | 105841 | 1.027475 | 0.008191 | 1079 |

| | | | | |
|---|---|---|---|---|
| Amotl2 | 56332 | 1.026573 | 0.000001 | 1080 |
| Tapbp | 21356 | 1.026459 | 8.69E-06 | 1081 |
| Lrp4 | 228357 | 1.023176 | 0.003009 | 1082 |
| Dnah6 | 330355 | 1.022226 | 0.031444 | 1083 |
| Plcd4 | 18802 | 1.020683 | 0.004215 | 1084 |
| Hells | 15201 | 1.02027 | 0.021225 | 1085 |
| Emilin2 | 246707 | 1.020058 | 9.39E-06 | 1086 |
| Smc2 | 14211 | 1.018576 | 0.003066 | 1087 |
| Ccnyl1 | 227210 | 1.018343 | 9.38E-06 | 1088 |
| Cnksr3 | 215748 | 1.017375 | 0.005103 | 1089 |
| Ccdc114 | 211535 | 1.015561 | 0.042697 | 1090 |
| Cdc42ep1 | 104445 | 1.015147 | 3.16E-06 | 1091 |
| Ctnna3 | 216033 | 1.015069 | 0.002948 | 1092 |
| Gpr37 | 14763 | 1.014449 | 0.000001 | 1093 |
| Nid1 | 18073 | 1.014222 | 0.00118 | 1094 |
| Fbn1 | 14118 | 1.010473 | 6.57E-06 | 1095 |
| Lap3 | 66988 | 1.010342 | 0.000787 | 1096 |
| Npnt | 114249 | 1.009058 | 0.012874 | 1097 |
| Pdlim4 | 30794 | 1.0079 | 0.015217 | 1098 |
| Trex1 | 22040 | 1.007089 | 0.019139 | 1099 |
| Lima1 | 65970 | 1.006156 | 0.003847 | 1100 |
| Gpsm2 | 76123 | 1.004139 | 0.017832 | 1101 |
| Tln1 | 21894 | 1.002561 | 0.045926 | 1102 |
| Mdfic | 16543 | 1.000799 | 0.000426 | 1103 |
| Jup | 16480 | 0.999516 | 0.000001 | 1104 |
| Ucp2 | 22228 | 0.999234 | 5.61E-06 | 1105 |
| Litaf | 56722 | 0.998612 | 0.00012 | 1106 |
| Casp1 | 12362 | 0.998278 | 0.005234 | 1107 |
| Grid2ip | 170935 | 0.996972 | 0.000001 | 1108 |
| Gm4790 | 215208 | 0.996762 | 0.000721 | 1109 |
| Slc4a2 | 20535 | 0.996635 | 0.000001 | 1110 |
| Sdc4 | 20971 | 0.996544 | 4.20E-05 | 1111 |
| Erbb2ip | 59079 | 0.995995 | 0.000001 | 1112 |
| Thbs3 | 21827 | 0.995254 | 0.047576 | 1113 |
| Elovl7 | 74559 | 0.991863 | 0.000001 | 1114 |
| Atp2a3 | 53313 | 0.991265 | 0.004105 | 1115 |
| Ccbl2 | 229905 | 0.989741 | 0.009202 | 1116 |
| Tnfaip8 | 106869 | 0.988711 | 0.004662 | 1117 |
| Lrp5 | 16973 | 0.985971 | 0.011992 | 1118 |
| Met | 17295 | 0.984391 | 1.73E-05 | 1119 |
| Tgfbr1 | 21812 | 0.984312 | 0.000402 | 1120 |
| Pstpip2 | 19201 | 0.983981 | 0.0086 | 1121 |
| Pdk4 | 27273 | 0.98055 | 0.014149 | 1122 |
| Slc4a4 | 54403 | 0.98034 | 0.037308 | 1123 |
| Casp7 | 12369 | 0.978823 | 0.006387 | 1124 |
| Acadl | 11363 | 0.976753 | 0.001033 | 1125 |
| Tnfrsf1b | 21938 | 0.976023 | 0.001091 | 1126 |

| | | | | |
|---|---|---|---|---|
| Tyrobp | 22177 | 0.975824 | 0.026509 | 1127 |
| Aldh1a2 | 19378 | 0.975797 | 0.00757 | 1128 |
| Galnt4 | 14426 | 0.975549 | 0.037414 | 1129 |
| Fzd5 | 14367 | 0.971394 | 0.01886 | 1130 |
| Dennd4c | 329877 | 0.970246 | 1.87E-06 | 1131 |
| Pygm | 19309 | 0.969553 | 0.002606 | 1132 |
| Gadd45g | 23882 | 0.969527 | 0.012711 | 1133 |
| Serpinh1 | 12406 | 0.968867 | 0.00067 | 1134 |
| Lepr | 16847 | 0.968759 | 0.029577 | 1135 |
| Gldc | 104174 | 0.968754 | 0.020153 | 1136 |
| Gpt | 76282 | 0.968422 | 0.039229 | 1137 |
| Lamc1 | 226519 | 0.967937 | 2.13E-05 | 1138 |
| Kcnk6 | 52150 | 0.967501 | 0.020754 | 1139 |
| Sh3pxd2b | 268396 | 0.965676 | 0.001596 | 1140 |
| Slc37a2 | 56857 | 0.96327 | 0.026056 | 1141 |
| Kcnj10 | 16513 | 0.963014 | 0.000115 | 1142 |
| Ptbp1 | 19205 | 0.960454 | 1.67E-05 | 1143 |
| Nfkbiz | 80859 | 0.959365 | 0.017619 | 1144 |
| Itgb5 | 16419 | 0.958488 | 0.003399 | 1145 |
| Phgdh | 236539 | 0.956148 | 0.000612 | 1146 |
| Phka1 | 18679 | 0.955452 | 0.000134 | 1147 |
| Swap70 | 20947 | 0.955432 | 0.001487 | 1148 |
| Vasp | 22323 | 0.953781 | 4.07E-05 | 1149 |
| Pxn | 19303 | 0.952483 | 7.67E-05 | 1150 |
| Rhog | 56212 | 0.952235 | 0.000001 | 1151 |
| Nckap5 | 210356 | 0.951818 | 0.001057 | 1152 |
| Gcnt1 | 14537 | 0.948873 | 0.012761 | 1153 |
| Tprn | 97031 | 0.947305 | 0.000458 | 1154 |
| Sulf1 | 240725 | 0.946918 | 0.000001 | 1155 |
| Pltp | 18830 | 0.944878 | 4.02E-06 | 1156 |
| Mcm4 | 17217 | 0.944868 | 0.000122 | 1157 |
| Ptpn14 | 19250 | 0.944546 | 0.01359 | 1158 |
| Rassf4 | 213391 | 0.944365 | 0.001182 | 1159 |
| Plekho2 | 102595 | 0.943736 | 0.000529 | 1160 |
| Oplah | 75475 | 0.940835 | 0.001761 | 1161 |
| Zeb2 | 24136 | 0.939002 | 0.000001 | 1162 |
| Bard1 | 12021 | 0.938815 | 0.009797 | 1163 |
| Myd88 | 17874 | 0.934278 | 0.014149 | 1164 |
| D16Ertd47 | 67102 | 0.933274 | 0.000623 | 1165 |
| Gtse1 | 29870 | 0.932043 | 0.037116 | 1166 |
| Sgk1 | 20393 | 0.929514 | 0.000001 | 1167 |
| Ptpn13 | 19249 | 0.928611 | 1.88E-05 | 1168 |
| Olig2 | 50913 | 0.928408 | 0.001312 | 1169 |
| Adam12 | 11489 | 0.927673 | 0.016598 | 1170 |
| Tst | 22117 | 0.925604 | 0.014275 | 1171 |
| Filip1l | 78749 | 0.92531 | 0.031556 | 1172 |
| Dock6 | 319899 | 0.922609 | 0.006489 | 1173 |

| | | | | |
|---|---|---|---|---|
| Kl | 16591 | 0.9224 | 0.0163 | 1174 |
| Fgfr2 | 14183 | 0.921604 | 0.000001 | 1175 |
| Plod3 | 26433 | 0.921339 | 7.46E-05 | 1176 |
| Atp7a | 11977 | 0.919904 | 0.000502 | 1177 |
| Hip1 | 215114 | 0.918498 | 0.000704 | 1178 |
| Zfp110 | 65020 | 0.916884 | 7.17E-06 | 1179 |
| Lama2 | 16773 | 0.914353 | 0.001034 | 1180 |
| 1700084C0 | 78465 | 0.913655 | 0.010739 | 1181 |
| Rp2h | 19889 | 0.913436 | 0.000001 | 1182 |
| Rgs16 | 19734 | 0.912032 | 0.001984 | 1183 |
| Map3k1 | 26401 | 0.911717 | 0.000001 | 1184 |
| Hadh | 15107 | 0.911196 | 0.004994 | 1185 |
| Lfng | 16848 | 0.909591 | 0.003527 | 1186 |
| Sgol2a | 68549 | 0.909291 | 0.040823 | 1187 |
| Serpinb9 | 20723 | 0.907472 | 0.012252 | 1188 |
| Fgd3 | 30938 | 0.904797 | 0.000103 | 1189 |
| Etv6 | 14011 | 0.904069 | 0.000001 | 1190 |
| Cbs | 12411 | 0.903483 | 0.030596 | 1191 |
| Itga6 | 16403 | 0.900741 | 0.0005 | 1192 |
| Slc7a11 | 26570 | 0.900502 | 0.000001 | 1193 |
| Pvrl4 | 71740 | 0.89677 | 0.01936 | 1194 |
| Soat1 | 20652 | 0.896496 | 0.000001 | 1195 |
| Lair1 | 52855 | 0.896005 | 0.045618 | 1196 |
| Pard3b | 72823 | 0.895118 | 5.05E-05 | 1197 |
| Gpam | 14732 | 0.893855 | 0.003613 | 1198 |
| Hapln2 | 73940 | 0.893471 | 0.005033 | 1199 |
| Drc7 | 330830 | 0.893164 | 0.016432 | 1200 |
| Bcas1 | 76960 | 0.892807 | 0.000492 | 1201 |
| Pgm5 | 226041 | 0.892131 | 0.002167 | 1202 |
| Cdt1 | 67177 | 0.891122 | 0.018406 | 1203 |
| Ampd3 | 11717 | 0.886846 | 0.001419 | 1204 |
| Cfap43 | 1E+08 | 0.885897 | 0.043675 | 1205 |
| Mog | 17441 | 0.883005 | 0.000001 | 1206 |
| Arhgap31 | 12549 | 0.881505 | 3.44E-06 | 1207 |
| Heatr5a | 320487 | 0.880006 | 0.001337 | 1208 |
| Csrp1 | 13007 | 0.87907 | 4.63E-05 | 1209 |
| Nde1 | 67203 | 0.877077 | 0.00433 | 1210 |
| Bgn | 12111 | 0.876445 | 0.022252 | 1211 |
| Shroom3 | 27428 | 0.876023 | 0.004853 | 1212 |
| Chdh | 218865 | 0.87489 | 0.014998 | 1213 |
| Ak7 | 78801 | 0.874364 | 0.031473 | 1214 |
| Thbs2 | 21826 | 0.873105 | 4.21E-05 | 1215 |
| Gltp | 56356 | 0.872647 | 0.000001 | 1216 |
| Sipa1 | 20469 | 0.871797 | 0.002948 | 1217 |
| Bcar3 | 29815 | 0.871698 | 0.002551 | 1218 |
| Phf11d | 219132 | 0.870759 | 0.001492 | 1219 |
| Timp3 | 21859 | 0.870066 | 0.000001 | 1220 |

| | | | | |
|---|---|---|---|---|
| Aebp1 | 11568 | 0.869336 | 0.013906 | 1221 |
| Celsr1 | 12614 | 0.869141 | 0.019329 | 1222 |
| Trim59 | 66949 | 0.867221 | 2.33E-06 | 1223 |
| Sgpl1 | 20397 | 0.865149 | 0.000229 | 1224 |
| Zbtb20 | 56490 | 0.863927 | 0.000001 | 1225 |
| Pcsk6 | 18553 | 0.86345 | 0.005874 | 1226 |
| Olig1 | 50914 | 0.861155 | 0.000313 | 1227 |
| Rab11fip1 | 75767 | 0.860897 | 0.016306 | 1228 |
| Prrc2c | 226562 | 0.860621 | 0.000001 | 1229 |
| Kif19a | 286942 | 0.860334 | 0.038213 | 1230 |
| Fyco1 | 17281 | 0.858065 | 0.000001 | 1231 |
| Adgrf5 | 224792 | 0.857938 | 0.01232 | 1232 |
| Nedd9 | 18003 | 0.850403 | 0.000279 | 1233 |
| Inppl1 | 16332 | 0.848525 | 0.004063 | 1234 |
| Carhsp1 | 52502 | 0.848319 | 0.001305 | 1235 |
| Cntn2 | 21367 | 0.847951 | 0.000001 | 1236 |
| 4931406CC | 70984 | 0.847726 | 0.008145 | 1237 |
| Appl2 | 216190 | 0.847098 | 0.00143 | 1238 |
| Gm3764 | 1E+08 | 0.846973 | 0.046042 | 1239 |
| Thbs4 | 21828 | 0.846606 | 0.024381 | 1240 |
| Il4ra | 16190 | 0.84628 | 0.002984 | 1241 |
| Fgd6 | 13998 | 0.846045 | 0.007147 | 1242 |
| Nkx2-2 | 18088 | 0.844297 | 0.006449 | 1243 |
| Pla2g16 | 225845 | 0.843828 | 0.000166 | 1244 |
| Plek | 56193 | 0.842793 | 0.000133 | 1245 |
| Ccl5 | 20304 | 0.838113 | 0.014978 | 1246 |
| Rin2 | 74030 | 0.835582 | 0.001782 | 1247 |
| Elovl1 | 54325 | 0.834536 | 0.021504 | 1248 |
| Cat | 12359 | 0.833812 | 9.49E-05 | 1249 |
| Srebf1 | 20787 | 0.8335 | 0.000001 | 1250 |
| Cd68 | 12514 | 0.830388 | 0.038595 | 1251 |
| Vamp8 | 22320 | 0.829146 | 0.016824 | 1252 |
| Oas1b | 23961 | 0.828753 | 0.002313 | 1253 |
| Plce1 | 74055 | 0.828221 | 0.002047 | 1254 |
| Sp1 | 20683 | 0.827391 | 0.000001 | 1255 |
| Wrn | 22427 | 0.827104 | 3.63E-05 | 1256 |
| H2-DMa | 14998 | 0.826857 | 0.007918 | 1257 |
| Gareml | 242915 | 0.826241 | 0.010237 | 1258 |
| Phyhd1 | 227696 | 0.825115 | 0.036067 | 1259 |
| Tob2 | 57259 | 0.824488 | 8.80E-05 | 1260 |
| Ncoa3 | 17979 | 0.82382 | 0.000001 | 1261 |
| Pld2 | 18806 | 0.823653 | 0.017284 | 1262 |
| Gpr37l1 | 171469 | 0.822378 | 0.010188 | 1263 |
| Acss1 | 68738 | 0.821576 | 0.019308 | 1264 |
| Sfrp1 | 20377 | 0.821505 | 0.013888 | 1265 |
| Wdr76 | 241627 | 0.82087 | 0.011441 | 1266 |
| Ppp1r18 | 76448 | 0.820102 | 0.005021 | 1267 |

| | | | | |
|---|---|---|---|---|
| Tgfa | 21802 | 0.818109 | 0.014171 | 1268 |
| Cdhr1 | 170677 | 0.817931 | 0.00082 | 1269 |
| Rasgrp3 | 240168 | 0.817257 | 9.34E-05 | 1270 |
| Smchd1 | 74355 | 0.816694 | 5.40E-06 | 1271 |
| Yeats2 | 208146 | 0.815488 | 0.000001 | 1272 |
| Cebpa | 12606 | 0.814759 | 0.008418 | 1273 |
| Cflar | 12633 | 0.814729 | 0.000181 | 1274 |
| Rcsd1 | 226594 | 0.814444 | 0.007538 | 1275 |
| Hadha | 97212 | 0.814382 | 0.000558 | 1276 |
| Rel | 19696 | 0.813253 | 0.001363 | 1277 |
| Irak2 | 108960 | 0.813061 | 0.01134 | 1278 |
| Meg3 | 17263 | 0.811924 | 0.000258 | 1279 |
| Hgf | 15234 | 0.811384 | 0.024789 | 1280 |
| Lgi4 | 243914 | 0.809299 | 0.017133 | 1281 |
| Ctnnal1 | 54366 | 0.809227 | 0.002364 | 1282 |
| Lck | 16818 | 0.809119 | 0.00917 | 1283 |
| Nhsl1 | 215819 | 0.808541 | 0.006837 | 1284 |
| Abcc4 | 239273 | 0.808325 | 0.036544 | 1285 |
| Hepacam | 72927 | 0.807021 | 0.000826 | 1286 |
| Dnm2 | 13430 | 0.806895 | 7.17E-06 | 1287 |
| Epb4.1l5 | 226352 | 0.806394 | 0.00171 | 1288 |
| Eps8 | 13860 | 0.805353 | 0.000001 | 1289 |
| Dchs1 | 233651 | 0.804549 | 0.001606 | 1290 |
| 2410004PC | 73667 | 0.803072 | 0.049498 | 1291 |
| Ubc | 22190 | 0.801349 | 0.000001 | 1292 |
| Pla2g7 | 27226 | 0.800155 | 0.00014 | 1293 |
| Crb2 | 241324 | 0.79984 | 0.010218 | 1294 |
| Klhdc7a | 242721 | 0.797014 | 0.010937 | 1295 |
| Gatm | 67092 | 0.796675 | 0.000001 | 1296 |
| Lrrc9 | 78257 | 0.796348 | 0.035088 | 1297 |
| Evi2a | 14017 | 0.795618 | 0.005427 | 1298 |
| Sgk3 | 170755 | 0.792151 | 0.001179 | 1299 |
| Gjb1 | 14618 | 0.789925 | 0.001994 | 1300 |
| Itgav | 16410 | 0.788635 | 0.000556 | 1301 |
| Slc12a2 | 20496 | 0.786415 | 0.000001 | 1302 |
| Agl | 77559 | 0.784783 | 0.000521 | 1303 |
| Myo9b | 17925 | 0.784766 | 2.15E-05 | 1304 |
| Abcd1 | 11666 | 0.784763 | 0.003807 | 1305 |
| Aldh2 | 11669 | 0.783898 | 0.003517 | 1306 |
| Prelp | 116847 | 0.782558 | 0.000706 | 1307 |
| Rab3il1 | 74760 | 0.78221 | 0.032706 | 1308 |
| Pmp22 | 18858 | 0.781814 | 0.00183 | 1309 |
| Ednrb | 13618 | 0.780309 | 0.026184 | 1310 |
| Mpdz | 17475 | 0.779179 | 0.000001 | 1311 |
| Fbxo32 | 67731 | 0.778289 | 0.000235 | 1312 |
| Antxr1 | 69538 | 0.777928 | 0.000145 | 1313 |
| Add3 | 27360 | 0.776569 | 8.36E-05 | 1314 |

| | | | | |
|---|---|---|---|---|
| Rhpn2 | 52428 | 0.776267 | 0.002813 | 1315 |
| Cmtm6 | 67213 | 0.776211 | 0.002287 | 1316 |
| Irf9 | 16391 | 0.7758 | 0.002313 | 1317 |
| Itga9 | 104099 | 0.774682 | 0.023452 | 1318 |
| Tril | 66873 | 0.774243 | 0.032018 | 1319 |
| Snap23 | 20619 | 0.772334 | 0.017804 | 1320 |
| Ube2l6 | 56791 | 0.771013 | 0.004419 | 1321 |
| Stat6 | 20852 | 0.767488 | 0.001881 | 1322 |
| Anxa5 | 11747 | 0.766671 | 0.000423 | 1323 |
| Cdc42ep2 | 104252 | 0.765807 | 0.00784 | 1324 |
| Kat2b | 18519 | 0.76347 | 0.000318 | 1325 |
| Trim67 | 330863 | 0.761152 | 0.001259 | 1326 |
| Lacc1 | 210808 | 0.76105 | 0.047952 | 1327 |
| Clic5 | 224796 | 0.760957 | 0.018562 | 1328 |
| S1pr5 | 94226 | 0.759912 | 0.000941 | 1329 |
| Tnxb | 81877 | 0.759645 | 0.001943 | 1330 |
| Kdm3a | 104263 | 0.757227 | 0.000188 | 1331 |
| Mal | 17153 | 0.756892 | 0.000001 | 1332 |
| Jmjd1c | 108829 | 0.756576 | 0.000001 | 1333 |
| Limd1 | 29806 | 0.753672 | 0.00052 | 1334 |
| Mtr | 238505 | 0.753201 | 4.21E-06 | 1335 |
| Acot11 | 329910 | 0.75313 | 0.017885 | 1336 |
| Miat | 330166 | 0.752284 | 0.006264 | 1337 |
| Lrrc16a | 68732 | 0.75214 | 0.000001 | 1338 |
| Vav3 | 57257 | 0.751443 | 0.03095 | 1339 |
| F3 | 14066 | 0.75095 | 0.021188 | 1340 |
| 2700049A0 | 76967 | 0.750038 | 0.000318 | 1341 |
| Kndc1 | 76484 | 0.749682 | 0.000313 | 1342 |
| Reep3 | 28193 | 0.749503 | 7.34E-05 | 1343 |
| Scamp2 | 24044 | 0.749031 | 0.017374 | 1344 |
| Atm | 11920 | 0.748646 | 0.000001 | 1345 |
| Hn1l | 52009 | 0.747767 | 0.010319 | 1346 |
| Pnpla2 | 66853 | 0.74758 | 0.004887 | 1347 |
| Erdr1 | 170942 | 0.747407 | 0.033906 | 1348 |
| Prob1 | 381148 | 0.746383 | 0.047282 | 1349 |
| Rab31 | 106572 | 0.746284 | 4.17E-05 | 1350 |
| Lrguk | 74354 | 0.745481 | 0.00137 | 1351 |
| Btd | 26363 | 0.74476 | 0.025651 | 1352 |
| Itga7 | 16404 | 0.744443 | 0.03778 | 1353 |
| Ctdsp1 | 227292 | 0.743665 | 0.000236 | 1354 |
| Fam46a | 212943 | 0.743359 | 4.26E-05 | 1355 |
| Dpyd | 99586 | 0.74203 | 0.027381 | 1356 |
| Serinc5 | 218442 | 0.740354 | 8.50E-05 | 1357 |
| Adgrd1 | 243277 | 0.738437 | 0.009354 | 1358 |
| Mast4 | 328329 | 0.736256 | 0.000001 | 1359 |
| Tyk2 | 54721 | 0.736196 | 0.029903 | 1360 |
| Npc1 | 18145 | 0.736159 | 3.10E-06 | 1361 |

| | | | | |
|---|---|---|---|---|
| Eml3 | 225898 | 0.735487 | 0.044509 | 1362 |
| Ift140 | 106633 | 0.734618 | 0.005978 | 1363 |
| Inf2 | 70435 | 0.734175 | 0.002634 | 1364 |
| Sertad1 | 55942 | 0.733977 | 0.026903 | 1365 |
| Ikzf2 | 22779 | 0.732996 | 0.00099 | 1366 |
| Col4a2 | 12827 | 0.730154 | 5.15E-05 | 1367 |
| Adamts20 | 223838 | 0.729971 | 0.01744 | 1368 |
| Tuba1c | 22146 | 0.729437 | 0.006343 | 1369 |
| Wsb1 | 78889 | 0.729376 | 0.002045 | 1370 |
| Ptprd | 19266 | 0.728687 | 0.000001 | 1371 |
| Zbtb21 | 114565 | 0.727623 | 0.001512 | 1372 |
| Itpkc | 233011 | 0.726357 | 0.002538 | 1373 |
| Efhd1 | 98363 | 0.72593 | 0.000749 | 1374 |
| Trib1 | 211770 | 0.724268 | 0.035578 | 1375 |
| Hspa2 | 15512 | 0.723926 | 7.47E-05 | 1376 |
| Mob1a | 232157 | 0.723061 | 2.29E-05 | 1377 |
| Cmpk2 | 22169 | 0.722406 | 0.002887 | 1378 |
| Id3 | 15903 | 0.71878 | 0.035897 | 1379 |
| Pold1 | 18971 | 0.718532 | 0.011874 | 1380 |
| Sox8 | 20681 | 0.718499 | 7.34E-05 | 1381 |
| Zhx2 | 387609 | 0.718498 | 1.06E-05 | 1382 |
| Mobp | 17433 | 0.718321 | 5.40E-06 | 1383 |
| Irs2 | 384783 | 0.718307 | 1.16E-06 | 1384 |
| Apbb1ip | 54519 | 0.716613 | 0.017735 | 1385 |
| Wdhd1 | 218973 | 0.716028 | 0.021154 | 1386 |
| Aco1 | 11428 | 0.714342 | 7.69E-05 | 1387 |
| Acap2 | 78618 | 0.714141 | 0.000001 | 1388 |
| Fryl | 72313 | 0.713349 | 0.000001 | 1389 |
| Mfn1 | 67414 | 0.712929 | 0.014377 | 1390 |
| 2810459M | 72792 | 0.711353 | 0.006506 | 1391 |
| Psme2 | 19188 | 0.710168 | 0.001379 | 1392 |
| Ltbp3 | 16998 | 0.707887 | 0.002216 | 1393 |
| St18 | 240690 | 0.706783 | 0.004898 | 1394 |
| Mga | 29808 | 0.705088 | 0.000001 | 1395 |
| Ranbp2 | 19386 | 0.704516 | 0.000001 | 1396 |
| Rrm1 | 20133 | 0.704463 | 0.001169 | 1397 |
| Abca8a | 217258 | 0.700224 | 0.008292 | 1398 |
| Soga1 | 320706 | 0.699467 | 0.000001 | 1399 |
| Myh14 | 71960 | 0.699065 | 0.005442 | 1400 |
| D8Ertd82e | 244418 | 0.697524 | 6.48E-06 | 1401 |
| Npc2 | 67963 | 0.697115 | 0.000701 | 1402 |
| Ctxn3 | 629147 | 0.695368 | 0.024042 | 1403 |
| Adcy7 | 11513 | 0.693524 | 0.019441 | 1404 |
| Rela | 19697 | 0.693324 | 0.001474 | 1405 |
| Unc5b | 107449 | 0.693202 | 1.62E-05 | 1406 |
| Sft2d2 | 108735 | 0.690979 | 0.000726 | 1407 |
| Abcg2 | 26357 | 0.689669 | 0.025837 | 1408 |

| | | | | |
|---|---|---|---|---|
| Il1rap | 16180 | 0.68897 | 1.52E-05 | 1409 |
| Pdlim5 | 56376 | 0.688719 | 0.000868 | 1410 |
| Trafd1 | 231712 | 0.688181 | 0.000618 | 1411 |
| Rps6ka1 | 20111 | 0.687964 | 0.001718 | 1412 |
| Bach1 | 12013 | 0.687675 | 0.000559 | 1413 |
| Ctnnd1 | 12388 | 0.687476 | 0.001336 | 1414 |
| Ptpn21 | 24000 | 0.686766 | 0.002908 | 1415 |
| Sall1 | 58198 | 0.686586 | 0.001296 | 1416 |
| 2-Sep | 18000 | 0.685466 | 1.43E-06 | 1417 |
| Ccdc141 | 545428 | 0.685168 | 0.00543 | 1418 |
| Zfp462 | 242466 | 0.685157 | 0.000001 | 1419 |
| Ankhd1 | 108857 | 0.683939 | 7.59E-06 | 1420 |
| Bcl2l11 | 12125 | 0.683491 | 0.012637 | 1421 |
| Casz1 | 69743 | 0.68334 | 0.022341 | 1422 |
| Ank2 | 109676 | 0.683339 | 0.000001 | 1423 |
| Npas3 | 27386 | 0.681095 | 0.004424 | 1424 |
| Aldh16a1 | 69748 | 0.680194 | 0.011846 | 1425 |
| Heg1 | 77446 | 0.678887 | 0.00019 | 1426 |
| Ide | 15925 | 0.678056 | 0.000887 | 1427 |
| Fads2 | 56473 | 0.67757 | 0.001508 | 1428 |
| Wwtr1 | 97064 | 0.677567 | 0.013574 | 1429 |
| Yap1 | 22601 | 0.674937 | 0.002364 | 1430 |
| Tnpo1 | 238799 | 0.674517 | 2.19E-06 | 1431 |
| Il6st | 16195 | 0.674272 | 0.000001 | 1432 |
| Eprs | 107508 | 0.673509 | 5.49E-05 | 1433 |
| Vcl | 22330 | 0.672482 | 0.001063 | 1434 |
| Nup98 | 269966 | 0.670982 | 0.000001 | 1435 |
| Hspb8 | 80888 | 0.669074 | 0.011183 | 1436 |
| Cers2 | 76893 | 0.668156 | 6.83E-05 | 1437 |
| Fmnl3 | 22379 | 0.666614 | 0.017284 | 1438 |
| Car2 | 12349 | 0.666535 | 6.00E-06 | 1439 |
| Tmem176b | 65963 | 0.665781 | 0.040954 | 1440 |
| Fermt2 | 218952 | 0.664789 | 8.23E-06 | 1441 |
| Ccnd3 | 12445 | 0.664568 | 0.001203 | 1442 |
| Tmod3 | 50875 | 0.664128 | 0.01152 | 1443 |
| Mid1 | 17318 | 0.663338 | 0.004873 | 1444 |
| Dusp16 | 70686 | 0.658832 | 0.000001 | 1445 |
| Vwa1 | 246228 | 0.658445 | 0.049252 | 1446 |
| Mthfd1 | 108156 | 0.6575 | 0.000381 | 1447 |
| Tspan2 | 70747 | 0.655251 | 0.000001 | 1448 |
| Psmb10 | 19171 | 0.655233 | 0.021453 | 1449 |
| Macf1 | 11426 | 0.653051 | 1.13E-06 | 1450 |
| Wnk1 | 232341 | 0.652676 | 0.000001 | 1451 |
| Polr2a | 20020 | 0.65167 | 0.000001 | 1452 |
| Rock1 | 19877 | 0.65062 | 0.001083 | 1453 |
| Pgm1 | 72157 | 0.649559 | 0.003881 | 1454 |
| Lpar1 | 14745 | 0.649481 | 0.015358 | 1455 |

| | | | | |
|---|---|---|---|---|
| Cryab | 12955 | 0.649389 | 0.000167 | 1456 |
| Phlpp1 | 98432 | 0.648076 | 0.000001 | 1457 |
| Scd1 | 20249 | 0.646431 | 0.001816 | 1458 |
| Opalin | 226115 | 0.646425 | 0.002543 | 1459 |
| Col6a2 | 12834 | 0.646117 | 0.032065 | 1460 |
| Cltc | 67300 | 0.646013 | 0.000001 | 1461 |
| Suclg2 | 20917 | 0.645972 | 0.028904 | 1462 |
| Fras1 | 231470 | 0.644196 | 0.000001 | 1463 |
| Nfkb1 | 18033 | 0.644193 | 0.009218 | 1464 |
| Fnbp1 | 14269 | 0.643722 | 0.000001 | 1465 |
| Igsf10 | 242050 | 0.642496 | 0.017463 | 1466 |
| Itgb1 | 16412 | 0.642113 | 0.000497 | 1467 |
| Isg15 | 1E+08 | 0.641339 | 0.032205 | 1468 |
| N4bp2 | 333789 | 0.641078 | 0.001046 | 1469 |
| Tubb2b | 73710 | 0.640463 | 0.010078 | 1470 |
| Rassf2 | 215653 | 0.639522 | 3.16E-05 | 1471 |
| Tspan15 | 70423 | 0.638735 | 0.01268 | 1472 |
| Cgnl1 | 68178 | 0.63866 | 0.014681 | 1473 |
| Gna13 | 14674 | 0.638198 | 0.000425 | 1474 |
| Sall3 | 20689 | 0.634947 | 0.049215 | 1475 |
| Car12 | 76459 | 0.633672 | 0.001977 | 1476 |
| Epb4.1l4a | 13824 | 0.633171 | 0.002 | 1477 |
| Sec16a | 227648 | 0.632115 | 0.000001 | 1478 |
| Tead1 | 21676 | 0.631242 | 0.000233 | 1479 |
| Lhfpl2 | 218454 | 0.629453 | 0.001328 | 1480 |
| Clk1 | 12747 | 0.628467 | 0.017768 | 1481 |
| Calb2 | 12308 | 0.62784 | 0.000202 | 1482 |
| Sh3d19 | 27059 | 0.627831 | 0.000188 | 1483 |
| Atad2 | 70472 | 0.626386 | 0.044037 | 1484 |
| Gpc5 | 103978 | 0.625633 | 0.035019 | 1485 |
| Kdm1b | 218214 | 0.625196 | 0.00819 | 1486 |
| Itgb8 | 320910 | 0.624631 | 9.00E-05 | 1487 |
| Fndc3b | 72007 | 0.624466 | 0.000119 | 1488 |
| Uhrf1bp1 | 224648 | 0.62421 | 9.35E-05 | 1489 |
| 2810474O1 | 67246 | 0.623307 | 0.001988 | 1490 |
| Pgd | 110208 | 0.622378 | 0.001656 | 1491 |
| Nhlrc2 | 66866 | 0.621303 | 0.01202 | 1492 |
| A330023F2 | 320977 | 0.620283 | 0.030765 | 1493 |
| Smc4 | 70099 | 0.618963 | 0.024028 | 1494 |
| Jun | 16476 | 0.617738 | 0.00088 | 1495 |
| Hsd17b4 | 15488 | 0.614972 | 0.003194 | 1496 |
| Luzp1 | 269593 | 0.614913 | 0.000001 | 1497 |
| Nav1 | 215690 | 0.614502 | 5.47E-06 | 1498 |
| Mdc1 | 240087 | 0.612798 | 0.014954 | 1499 |
| Kmt2c | 231051 | 0.610862 | 0.000001 | 1500 |
| Cemip | 80982 | 0.610549 | 0.000718 | 1501 |
| Hipk2 | 15258 | 0.609628 | 0.000001 | 1502 |

| | | | | |
|---|---|---|---|---|
| Rnf111 | 93836 | 0.60953 | 0.000186 | 1503 |
| Prkdc | 19090 | 0.608297 | 0.000699 | 1504 |
| Prdx6 | 11758 | 0.607689 | 0.017971 | 1505 |
| Kdm6b | 216850 | 0.607311 | 0.027972 | 1506 |
| Nav2 | 78286 | 0.606062 | 0.002177 | 1507 |
| Ptbp3 | 230257 | 0.606023 | 5.24E-06 | 1508 |
| Llgl1 | 16897 | 0.605598 | 0.001739 | 1509 |
| Prpf8 | 192159 | 0.603971 | 0.000001 | 1510 |
| Qk | 19317 | 0.603476 | 0.000001 | 1511 |
| Dync1h1 | 13424 | 0.601684 | 0.000001 | 1512 |
| Nwd1 | 319555 | 0.600878 | 0.013159 | 1513 |
| Spty2d1 | 101685 | 0.600594 | 0.010188 | 1514 |
| Aldh6a1 | 104776 | 0.600539 | 0.027461 | 1515 |
| Nbeal1 | 269198 | 0.600135 | 5.99E-06 | 1516 |
| Hps5 | 246694 | 0.599897 | 0.017618 | 1517 |
| Bicc1 | 83675 | 0.599264 | 0.007732 | 1518 |
| Lipe | 16890 | 0.598865 | 0.044753 | 1519 |
| Idh1 | 15926 | 0.597014 | 0.000411 | 1520 |
| Nr4a2 | 18227 | 0.595621 | 0.02193 | 1521 |
| Cep350 | 74081 | 0.594972 | 0.000001 | 1522 |
| Dip2b | 239667 | 0.594366 | 0.000001 | 1523 |
| Nup153 | 218210 | 0.594272 | 6.90E-06 | 1524 |
| BC034090 | 207792 | 0.593785 | 0.014538 | 1525 |
| Ddr2 | 18214 | 0.593563 | 2.20E-06 | 1526 |
| Pllp | 67801 | 0.592621 | 0.001657 | 1527 |
| Erich3 | 209601 | 0.592016 | 0.00179 | 1528 |
| Lnx2 | 140887 | 0.591689 | 0.026615 | 1529 |
| Cald1 | 109624 | 0.588604 | 0.004818 | 1530 |
| Zcchc24 | 71918 | 0.588375 | 0.003481 | 1531 |
| Shprh | 268281 | 0.588041 | 7.46E-06 | 1532 |
| Ssfa2 | 70599 | 0.586921 | 0.006632 | 1533 |
| Kalrn | 545156 | 0.585229 | 0.000001 | 1534 |
| Nbeal2 | 235627 | 0.584058 | 0.008233 | 1535 |
| Dusp10 | 63953 | 0.583877 | 0.027676 | 1536 |
| Scd2 | 20250 | 0.581158 | 5.60E-05 | 1537 |
| Map4k4 | 26921 | 0.580992 | 0.000001 | 1538 |
| Arsg | 74008 | 0.578327 | 0.009767 | 1539 |
| Birc6 | 12211 | 0.577803 | 0.000001 | 1540 |
| Shisa5 | 66940 | 0.577517 | 0.006088 | 1541 |
| Nup160 | 59015 | 0.577434 | 0.00068 | 1542 |
| Cd2ap | 12488 | 0.576275 | 0.003739 | 1543 |
| Cep295 | 319675 | 0.576007 | 1.04E-05 | 1544 |
| Lrrc3 | 237387 | 0.5755 | 0.02951 | 1545 |
| Man2b1 | 17159 | 0.574267 | 0.02354 | 1546 |
| Cd63 | 12512 | 0.573716 | 0.034804 | 1547 |
| Nr2f2 | 11819 | 0.573634 | 0.039462 | 1548 |
| Zbtb1 | 268564 | 0.573178 | 0.010177 | 1549 |

| | | | | |
|---|---|---|---|---|
| Stk38 | 106504 | 0.572613 | 0.023715 | 1550 |
| Gpr146 | 80290 | 0.572324 | 0.017974 | 1551 |
| Spata13 | 219140 | 0.570961 | 0.00045 | 1552 |
| Afap1 | 70292 | 0.570731 | 6.93E-06 | 1553 |
| Ep300 | 328572 | 0.568787 | 0.000001 | 1554 |
| Klhl4 | 237010 | 0.56852 | 0.001761 | 1555 |
| Mccc1 | 72039 | 0.567324 | 0.003122 | 1556 |
| Kif13a | 16553 | 0.566699 | 3.48E-05 | 1557 |
| Ubr3 | 68795 | 0.566382 | 0.000001 | 1558 |
| Plekhb1 | 27276 | 0.565375 | 0.000189 | 1559 |
| Tmcc3 | 319880 | 0.565095 | 2.44E-05 | 1560 |
| Gna12 | 14673 | 0.564361 | 2.13E-05 | 1561 |
| Rbms2 | 56516 | 0.563668 | 0.027989 | 1562 |
| Edem1 | 192193 | 0.563015 | 7.66E-05 | 1563 |
| Itfg3 | 106581 | 0.562792 | 0.037874 | 1564 |
| Ubr5 | 70790 | 0.562444 | 0.000001 | 1565 |
| Zfp638 | 18139 | 0.562396 | 0.000001 | 1566 |
| Rbpj | 19664 | 0.561813 | 0.000589 | 1567 |
| Son | 20658 | 0.559017 | 1.26E-05 | 1568 |
| Mbp | 17196 | 0.558649 | 0.000001 | 1569 |
| Acad11 | 102632 | 0.558258 | 0.016498 | 1570 |
| Sowahc | 268301 | 0.557561 | 0.000874 | 1571 |
| Pcdhgc3 | 93706 | 0.557423 | 0.006607 | 1572 |
| Lrpprc | 72416 | 0.55727 | 0.000001 | 1573 |
| Cyfip1 | 20430 | 0.556532 | 0.001228 | 1574 |
| Pikfyve | 18711 | 0.555855 | 0.000001 | 1575 |
| Shroom4 | 208431 | 0.555497 | 0.017956 | 1576 |
| H2-T22 | 15039 | 0.554797 | 0.011204 | 1577 |
| Cttnbp2nl | 80281 | 0.55279 | 0.001003 | 1578 |
| Mitf | 17342 | 0.552032 | 0.019825 | 1579 |
| Scpep1 | 74617 | 0.551261 | 0.003095 | 1580 |
| Tns1 | 21961 | 0.550629 | 5.67E-05 | 1581 |
| Arid1b | 239985 | 0.548982 | 0.016264 | 1582 |
| Capn2 | 12334 | 0.547945 | 0.005815 | 1583 |
| Mob3a | 208228 | 0.543073 | 0.010519 | 1584 |
| Aldh4a1 | 212647 | 0.542635 | 0.019086 | 1585 |
| Nbr1 | 17966 | 0.540783 | 0.000704 | 1586 |
| Zscan29 | 99334 | 0.540731 | 0.035019 | 1587 |
| Nfat5 | 54446 | 0.53939 | 0.000533 | 1588 |
| Dhx9 | 13211 | 0.539343 | 0.000001 | 1589 |
| Gca | 227960 | 0.539228 | 0.026579 | 1590 |
| Cep97 | 74201 | 0.537501 | 0.003039 | 1591 |
| Midn | 59090 | 0.537177 | 0.002048 | 1592 |
| Foxn3 | 71375 | 0.534096 | 7.51E-05 | 1593 |
| Slc44a1 | 100434 | 0.533256 | 3.16E-06 | 1594 |
| Sf3b3 | 101943 | 0.532858 | 0.000303 | 1595 |
| Ascc3 | 77987 | 0.531684 | 4.73E-05 | 1596 |

| | | | | |
|---|---|---|---|---|
| Rbm15 | 229700 | 0.530928 | 2.54E-05 | 1597 |
| Bcorl1 | 320376 | 0.530413 | 0.001039 | 1598 |
| Pabpc1 | 18458 | 0.529516 | 0.000606 | 1599 |
| Atl3 | 109168 | 0.528768 | 0.001501 | 1600 |
| Plagl2 | 54711 | 0.528585 | 0.025429 | 1601 |
| Bak1 | 12018 | 0.528279 | 0.004847 | 1602 |
| Zfp536 | 243937 | 0.527827 | 3.43E-06 | 1603 |
| Lrp1 | 16971 | 0.527466 | 1.92E-05 | 1604 |
| Zswim6 | 67263 | 0.527058 | 3.08E-05 | 1605 |
| Nampt | 59027 | 0.52597 | 0.00047 | 1606 |
| Slc17a6 | 140919 | 0.525934 | 0.047113 | 1607 |
| Slit1 | 20562 | 0.524929 | 7.92E-05 | 1608 |
| Ptpn23 | 104831 | 0.524657 | 0.001939 | 1609 |
| Ap3b1 | 11774 | 0.522187 | 0.002509 | 1610 |
| Ddr1 | 12305 | 0.521613 | 5.02E-05 | 1611 |
| Creb3l2 | 208647 | 0.52133 | 0.004794 | 1612 |
| 4931406P1 | 233103 | 0.520782 | 0.007684 | 1613 |
| Cblb | 208650 | 0.52078 | 0.000306 | 1614 |
| Sytl2 | 83671 | 0.520303 | 0.005427 | 1615 |
| Pfas | 237823 | 0.519226 | 0.014567 | 1616 |
| Pamr1 | 210622 | 0.518649 | 0.019555 | 1617 |
| Col9a3 | 12841 | 0.518407 | 0.026729 | 1618 |
| Pcx | 18563 | 0.518097 | 0.008743 | 1619 |
| Scara3 | 219151 | 0.517566 | 0.040183 | 1620 |
| Ddb1 | 13194 | 0.517563 | 1.10E-06 | 1621 |
| Mcm7 | 17220 | 0.516258 | 0.005517 | 1622 |
| Arid2 | 77044 | 0.5159 | 0.000001 | 1623 |
| Sox4 | 20677 | 0.515699 | 0.004432 | 1624 |
| Tdrd7 | 100121 | 0.515284 | 0.004491 | 1625 |
| Morc3 | 338467 | 0.513954 | 0.007545 | 1626 |
| Msh6 | 17688 | 0.512741 | 0.000596 | 1627 |
| S100a16 | 67860 | 0.512461 | 0.026372 | 1628 |
| Igsf9b | 235086 | 0.512448 | 0.001826 | 1629 |
| Ttc37 | 218343 | 0.512238 | 0.000663 | 1630 |
| Anxa6 | 11749 | 0.511606 | 0.001035 | 1631 |
| Ocln | 18260 | 0.510369 | 0.023807 | 1632 |
| Glud1 | 14661 | 0.509086 | 0.007625 | 1633 |
| Rorb | 225998 | 0.509028 | 0.010177 | 1634 |
| Alms1 | 236266 | 0.508805 | 9.00E-05 | 1635 |
| Nptx2 | 53324 | 0.50873 | 0.004054 | 1636 |
| Hnrnpf | 98758 | 0.508053 | 0.000535 | 1637 |
| Atp11c | 320940 | 0.507387 | 0.024295 | 1638 |
| Cdkn1a | 12575 | 0.506894 | 0.010519 | 1639 |
| Myo1c | 17913 | 0.506857 | 0.034804 | 1640 |
| Traf6 | 22034 | 0.506268 | 0.026056 | 1641 |
| Cc2d2a | 231214 | 0.505261 | 0.023101 | 1642 |
| Man2b2 | 17160 | 0.504981 | 0.027029 | 1643 |

| | | | | |
|---|---|---|---|---|
| Kcnk13 | 217826 | 0.50481 | 0.022893 | 1644 |
| Prr14l | 215476 | 0.504535 | 0.000001 | 1645 |
| Rictor | 78757 | 0.504293 | 0.000001 | 1646 |
| Epb4.1 | 269587 | 0.502869 | 0.02686 | 1647 |
| Pik3c2b | 240752 | 0.502597 | 0.000936 | 1648 |
| Mybbp1a | 18432 | 0.500861 | 0.000331 | 1649 |
| Nfia | 18027 | 0.500647 | 0.001895 | 1650 |
| Ncapd2 | 68298 | 0.500433 | 0.042289 | 1651 |
| Tbc1d5 | 72238 | 0.500318 | 7.67E-05 | 1652 |
| Trim36 | 28105 | 0.500098 | 0.029693 | 1653 |
