## Supplementary Table 2 for "Lateral Septal Circuits Govern Schizophrenia-Like Effects of Ketamine on Social Behavior"

**Supplementary Table 2 - Downregulated Translatome**

| symbol | entrez | logfc | adjpv | |
|---|---|---|---|---|
| Mrpl17 | 27397 | -0.50001 | 9.58E-05 | 1 |
| Rps14 | 20044 | -0.50022 | 0.000001 | 2 |
| Fau | 14109 | -0.50027 | 0.000559 | 3 |
| Med30 | 69790 | -0.50035 | 0.025429 | 4 |
| Sstr3 | 20607 | -0.50075 | 0.001172 | 5 |
| B9d1 | 27078 | -0.5008 | 0.04005 | 6 |
| Resp18 | 19711 | -0.50123 | 0.000777 | 7 |
| Mrpl52 | 68836 | -0.50143 | 0.007567 | 8 |
| Ift20 | 55978 | -0.50182 | 0.000718 | 9 |
| 1500009L16Rik | 69784 | -0.50184 | 0.002084 | 10 |
| Dpy30 | 66310 | -0.50188 | 0.017131 | 11 |
| Sstr5 | 20609 | -0.50194 | 0.03456 | 12 |
| Ndufb5 | 66046 | -0.50298 | 1.20E-05 | 13 |
| 1810043G02Rik | 67884 | -0.50341 | 0.017603 | 14 |
| Mrps14 | 64659 | -0.50427 | 0.000457 | 15 |
| Tmem17 | 103765 | -0.50476 | 0.028166 | 16 |
| Trappc2l | 59005 | -0.50476 | 0.001512 | 17 |
| Fam213b | 66469 | -0.50478 | 0.003016 | 18 |
| Mrps21 | 66292 | -0.50508 | 0.000407 | 19 |
| Rpl14 | 67115 | -0.50524 | 0.001645 | 20 |
| Selm | 114679 | -0.50528 | 0.000991 | 21 |
| Cript | 56724 | -0.50669 | 3.84E-05 | 22 |
| Rps12 | 20042 | -0.50821 | 0.032506 | 23 |
| Arl16 | 70317 | -0.5085 | 0.005162 | 24 |
| Bex2 | 12069 | -0.50895 | 0.002043 | 25 |
| Eef1b2 | 55949 | -0.50908 | 0.030454 | 26 |
| 1500011B03Rik | 66236 | -0.50909 | 9.74E-05 | 27 |
| Arhgdig | 14570 | -0.51016 | 0.000723 | 28 |
| Srrd | 70118 | -0.51073 | 0.007232 | 29 |
| Dtd2 | 328092 | -0.51073 | 0.044092 | 30 |
| Cwc15 | 66070 | -0.5118 | 0.006022 | 31 |
| Vstm2l | 277432 | -0.5119 | 0.000001 | 32 |
| Ppih | 66101 | -0.51213 | 0.007605 | 33 |
| Cgref1 | 68567 | -0.5127 | 0.00028 | 34 |
| Pigp | 56176 | -0.51314 | 0.001367 | 35 |
| Mkks | 59030 | -0.51338 | 0.001718 | 36 |
| Adck3 | 67426 | -0.51344 | 0.000105 | 37 |
| Mettl23 | 74319 | -0.51345 | 0.038607 | 38 |
| Fam96b | 68523 | -0.5148 | 0.029985 | 39 |
| Rps8 | 20116 | -0.51563 | 0.000718 | 40 |
| 1700052K11Rik | 73431 | -0.51568 | 0.04388 | 41 |
| Rpl26 | 19941 | -0.51753 | 1.31E-05 | 42 |
| Cox7a2 | 12866 | -0.51782 | 0.000001 | 43 |
| Avpi1 | 69534 | -0.51801 | 0.002076 | 44 |
| Fabp3 | 14077 | -0.51908 | 5.05E-05 | 45 |

| | | | | |
|---|---|---|---|---|
| Ndufs7 | 75406 | -0.52013 | 1.65E-06 | 46 |
| Hspe1 | 15528 | -0.52044 | 3.84E-05 | 47 |
| Ict1 | 68572 | -0.52068 | 1.70E-05 | 48 |
| Lsm2 | 27756 | -0.52074 | 0.021645 | 49 |
| Myeov2 | 66915 | -0.52133 | 9.97E-05 | 50 |
| Mt3 | 17751 | -0.52226 | 0.000001 | 51 |
| Ndufb3 | 66495 | -0.5225 | 0.000304 | 52 |
| Rps27a | 78294 | -0.52305 | 0.004794 | 53 |
| Acyp2 | 75572 | -0.52314 | 0.010847 | 54 |
| Mrpl23 | 19935 | -0.52323 | 0.009758 | 55 |
| Mrpl54 | 66047 | -0.52355 | 0.003028 | 56 |
| Rps25 | 75617 | -0.52393 | 3.61E-06 | 57 |
| Uba52 | 22186 | -0.52413 | 0.000001 | 58 |
| Rpl22l1 | 68028 | -0.52415 | 0.000218 | 59 |
| Mrpl12 | 56282 | -0.52468 | 1.13E-06 | 60 |
| Alkbh7 | 66400 | -0.52677 | 0.042991 | 61 |
| Swi5 | 72931 | -0.52678 | 3.73E-05 | 62 |
| Atp5h | 71679 | -0.52715 | 0.000001 | 63 |
| Mesdc2 | 67943 | -0.52795 | 0.00332 | 64 |
| Ensa | 56205 | -0.52869 | 1.97E-05 | 65 |
| Snrpf | 69878 | -0.52967 | 0.008743 | 66 |
| Hras | 15461 | -0.52984 | 0.000001 | 67 |
| Higd2a | 67044 | -0.53021 | 1.56E-06 | 68 |
| Rps17 | 20068 | -0.53057 | 7.92E-05 | 69 |
| Tceb2 | 67673 | -0.53075 | 0.000001 | 70 |
| Nme1 | 18102 | -0.53094 | 7.41E-06 | 71 |
| Ppapdc1b | 71910 | -0.5312 | 0.008963 | 72 |
| Ndufa12 | 66414 | -0.53195 | 0.000001 | 73 |
| Trpc7 | 26946 | -0.53199 | 0.012056 | 74 |
| 1110059G10Rik | 66202 | -0.53251 | 0.003208 | 75 |
| N6amt1 | 67768 | -0.53381 | 0.000127 | 76 |
| Dnajc15 | 66148 | -0.53637 | 0.000135 | 77 |
| Dnajc17 | 69408 | -0.53649 | 0.034896 | 78 |
| Rpl34 | 68436 | -0.53741 | 0.000816 | 79 |
| Mtfp1 | 67900 | -0.53806 | 7.72E-05 | 80 |
| Ndufv2 | 72900 | -0.53926 | 0.000001 | 81 |
| Rpl36al | 66483 | -0.53986 | 0.000294 | 82 |
| Mrpl20 | 66448 | -0.54022 | 0.00046 | 83 |
| Hist3h2ba | 78303 | -0.54105 | 0.004183 | 84 |
| Gtf2a2 | 235459 | -0.5416 | 0.001596 | 85 |
| Ly6h | 23934 | -0.54237 | 1.16E-06 | 86 |
| Atp6v0b | 114143 | -0.54301 | 0.000001 | 87 |
| Cox6a1 | 12861 | -0.54345 | 0.000001 | 88 |
| 1110032A03Rik | 68721 | -0.54396 | 0.000433 | 89 |
| Cox14 | 66379 | -0.54405 | 1.67E-05 | 90 |
| Tmsb4x | 19241 | -0.54503 | 0.000001 | 91 |
| 1700001L19Rik | 69315 | -0.54853 | 0.000556 | 92 |

| | | | | |
|---|---|---|---|---|
| Ndufb9 | 66218 | -0.54877 | 6.00E-06 | 93 |
| Mien1 | 103742 | -0.54902 | 5.70E-05 | 94 |
| Ndufb11 | 104130 | -0.55198 | 7.28E-06 | 95 |
| Nnat | 18111 | -0.55216 | 1.20E-06 | 96 |
| Nhp2 | 52530 | -0.55366 | 0.01093 | 97 |
| Mrpl27 | 94064 | -0.55434 | 8.84E-06 | 98 |
| Coa3 | 52469 | -0.55442 | 7.13E-06 | 99 |
| Gm14295 | 1E+08 | -0.55518 | 0.00828 | 100 |
| Hcfc1r1 | 353502 | -0.55655 | 6.05E-06 | 101 |
| Cox4i1 | 12857 | -0.55868 | 0.000001 | 102 |
| Mgst3 | 66447 | -0.55885 | 0.000163 | 103 |
| Elof1 | 66126 | -0.5594 | 4.10E-06 | 104 |
| Dpcd | 226162 | -0.55983 | 5.53E-05 | 105 |
| Tmem147 | 69804 | -0.56038 | 2.13E-05 | 106 |
| Emc4 | 68032 | -0.56114 | 0.000001 | 107 |
| Ndufaf5 | 69487 | -0.5629 | 0.001263 | 108 |
| Mrpl36 | 94066 | -0.56376 | 0.001997 | 109 |
| Pop7 | 74097 | -0.56445 | 0.002369 | 110 |
| Atp6v1f | 66144 | -0.56586 | 0.000001 | 111 |
| Rpl31 | 114641 | -0.56716 | 7.10E-06 | 112 |
| Mea1 | 17256 | -0.56737 | 7.05E-05 | 113 |
| Rpl36 | 54217 | -0.56738 | 1.43E-06 | 114 |
| Timm10 | 30059 | -0.56783 | 0.00024 | 115 |
| Deb1 | 26901 | -0.56899 | 0.001712 | 116 |
| Cycs | 13063 | -0.57029 | 0.000001 | 117 |
| Dynlrb1 | 67068 | -0.57062 | 1.00E-06 | 118 |
| Jagn1 | 67767 | -0.57082 | 0.000217 | 119 |
| Tma7 | 66167 | -0.57171 | 1.16E-06 | 120 |
| Snrnp27 | 66618 | -0.57187 | 6.42E-05 | 121 |
| Ubl5 | 66177 | -0.57251 | 0.000001 | 122 |
| Cox7c | 12867 | -0.57319 | 0.000001 | 123 |
| Znhit3 | 448850 | -0.57384 | 0.009901 | 124 |
| Atp5g1 | 11951 | -0.57464 | 0.000001 | 125 |
| Tomm5 | 68512 | -0.57542 | 0.001908 | 126 |
| Mrps28 | 66230 | -0.57584 | 0.021686 | 127 |
| 1110001J03Rik | 66117 | -0.57613 | 0.004347 | 128 |
| Nme5 | 75533 | -0.57618 | 0.005575 | 129 |
| Zcrb1 | 67197 | -0.57642 | 0.003275 | 130 |
| Ppp1r11 | 76497 | -0.57668 | 0.000001 | 131 |
| Gm10076 | 1E+08 | -0.57693 | 0.00014 | 132 |
| Ndufs5 | 595136 | -0.57815 | 1.07E-05 | 133 |
| Tmem256 | 69186 | -0.57855 | 0.016616 | 134 |
| Tmem258 | 69038 | -0.57949 | 0.009345 | 135 |
| Mrps33 | 14548 | -0.57961 | 0.000407 | 136 |
| Ndufc2 | 68197 | -0.57993 | 0.000001 | 137 |
| Mrps36 | 66128 | -0.58106 | 7.66E-05 | 138 |
| Pet100 | 1.01E+08 | -0.58168 | 0.001057 | 139 |

| | | | | |
|---|---|---|---|---|
| Lamtor2 | 83409 | -0.58441 | 0.001495 | 140 |
| Ndufv3 | 78330 | -0.58467 | 0.000001 | 141 |
| Cetn3 | 12626 | -0.5856 | 0.032922 | 142 |
| Ccdc124 | 234388 | -0.58872 | 1.16E-06 | 143 |
| Atox1 | 11927 | -0.58911 | 0.006331 | 144 |
| Coprs | 66423 | -0.58964 | 9.54E-05 | 145 |
| Amn | 93835 | -0.59053 | 0.045937 | 146 |
| Gpx4 | 625249 | -0.59091 | 0.000001 | 147 |
| Gpx3 | 14778 | -0.59141 | 0.00298 | 148 |
| Bola3 | 78653 | -0.59247 | 0.010583 | 149 |
| 1700021F05Rik | 67851 | -0.59374 | 0.019643 | 150 |
| Prkrip1 | 66801 | -0.59392 | 0.035282 | 151 |
| Praf2 | 54637 | -0.59434 | 0.000001 | 152 |
| Snrpe | 20643 | -0.59462 | 0.001331 | 153 |
| Nol12 | 97961 | -0.59623 | 0.019692 | 154 |
| Mrpl21 | 353242 | -0.59638 | 0.000662 | 155 |
| Saysd1 | 67509 | -0.59671 | 0.033693 | 156 |
| Polr2i | 69920 | -0.59818 | 0.006616 | 157 |
| Ndufc1 | 66377 | -0.59861 | 1.28E-06 | 158 |
| Serp2 | 72661 | -0.59903 | 0.000001 | 159 |
| Rpl41 | 67945 | -0.59938 | 0.000001 | 160 |
| Ndufb8 | 67264 | -0.60027 | 0.000001 | 161 |
| N6amt2 | 68043 | -0.6017 | 0.002223 | 162 |
| Ndufa11 | 69875 | -0.60224 | 2.33E-06 | 163 |
| Ndufa1 | 54405 | -0.6025 | 0.000107 | 164 |
| Lsm7 | 66094 | -0.6028 | 8.77E-05 | 165 |
| Esyt3 | 272636 | -0.60301 | 0.000425 | 166 |
| Atp5e | 67126 | -0.60354 | 1.67E-05 | 167 |
| Ndufb6 | 230075 | -0.60429 | 3.75E-06 | 168 |
| Fkbp3 | 30795 | -0.60435 | 0.001595 | 169 |
| Fam162a | 70186 | -0.6047 | 0.000746 | 170 |
| Cox6b1 | 110323 | -0.60558 | 0.000001 | 171 |
| Rps21 | 66481 | -0.60568 | 0.000449 | 172 |
| Ramp1 | 51801 | -0.60573 | 0.006865 | 173 |
| Nop10 | 66181 | -0.60615 | 3.89E-05 | 174 |
| 1500011K16Rik | 67885 | -0.60817 | 0.006154 | 175 |
| Denr | 68184 | -0.61046 | 0.001034 | 176 |
| Atp5l | 27425 | -0.61289 | 0.000001 | 177 |
| Ndufs6 | 407785 | -0.61789 | 5.71E-05 | 178 |
| Rpl36a | 19982 | -0.62235 | 0.032321 | 179 |
| Nrgn | 64011 | -0.62354 | 3.11E-06 | 180 |
| Cxcl14 | 57266 | -0.62744 | 3.29E-05 | 181 |
| Uqcrb | 67530 | -0.62794 | 0.000001 | 182 |
| Lin7b | 22342 | -0.62813 | 3.45E-05 | 183 |
| Cox8a | 12868 | -0.62835 | 0.000001 | 184 |
| Cck | 12424 | -0.62896 | 0.026298 | 185 |
| Cox16 | 66272 | -0.62958 | 0.002223 | 186 |

| | | | | |
|---|---|---|---|---|
| Ost4 | 67695 | -0.62958 | 0.000322 | 187 |
| Rpp21 | 67676 | -0.63022 | 0.012815 | 188 |
| Ndufa4 | 17992 | -0.63355 | 0.000001 | 189 |
| Morn2 | 378462 | -0.63455 | 0.003327 | 190 |
| Gm29514 | 1.03E+08 | -0.6346 | 0.046786 | 191 |
| Ndufa2 | 17991 | -0.6346 | 0.000001 | 192 |
| Mtx1 | 17827 | -0.63576 | 0.0027 | 193 |
| Phtf2 | 68770 | -0.63733 | 0.018 | 194 |
| Ndufa7 | 66416 | -0.64118 | 0.000001 | 195 |
| Ccdc74a | 72315 | -0.64148 | 0.035127 | 196 |
| Nenf | 66208 | -0.64154 | 1.38E-05 | 197 |
| Tmem91 | 320208 | -0.64196 | 0.000001 | 198 |
| Pam16 | 66449 | -0.64316 | 4.79E-06 | 199 |
| Dnajc4 | 57431 | -0.64436 | 0.011951 | 200 |
| Timm10b | 14356 | -0.64601 | 0.0005 | 201 |
| Snrpd2 | 107686 | -0.64627 | 2.58E-06 | 202 |
| Sst | 20604 | -0.64632 | 0.000001 | 203 |
| Lamb3 | 16780 | -0.64668 | 0.03104 | 204 |
| 1110008P14Rik | 73737 | -0.64732 | 1.17E-05 | 205 |
| 1810022K09Rik | 69126 | -0.64878 | 0.000185 | 206 |
| Grcc10 | 14790 | -0.64908 | 0.000001 | 207 |
| Cox19 | 68033 | -0.64951 | 0.04985 | 208 |
| Shfm1 | 20422 | -0.65586 | 0.007888 | 209 |
| Tmem261 | 66928 | -0.65637 | 4.85E-05 | 210 |
| Gm1673 | 381633 | -0.65756 | 0.008368 | 211 |
| Mrpl42 | 67270 | -0.65882 | 7.41E-06 | 212 |
| Ndufa6 | 67130 | -0.65911 | 0.000001 | 213 |
| 1810043H04Rik | 208501 | -0.66008 | 0.002028 | 214 |
| Uqcr10 | 66152 | -0.66415 | 0.000001 | 215 |
| Tmsb10 | 19240 | -0.6651 | 1.31E-05 | 216 |
| Pcp4 | 18546 | -0.66923 | 0.002187 | 217 |
| Pop5 | 117109 | -0.66929 | 0.002844 | 218 |
| Timm8b | 30057 | -0.66991 | 0.000001 | 219 |
| Ndufs8 | 225887 | -0.67088 | 1.30E-05 | 220 |
| Ccl27a | 20301 | -0.67753 | 0.000217 | 221 |
| Naa38 | 78304 | -0.67804 | 3.25E-05 | 222 |
| Ndufab1 | 70316 | -0.67867 | 0.000001 | 223 |
| Uqcc2 | 67267 | -0.68762 | 0.000001 | 224 |
| Zfp560 | 434377 | -0.69022 | 0.003206 | 225 |
| Cebpzos | 68554 | -0.69038 | 0.000901 | 226 |
| Uqcrh | 66576 | -0.69283 | 0.000001 | 227 |
| Slirp | 380773 | -0.69431 | 0.000137 | 228 |
| Romo1 | 67067 | -0.69789 | 0.000849 | 229 |
| 2010204K13Rik | 68355 | -0.69849 | 3.58E-06 | 230 |
| Polr2k | 17749 | -0.69932 | 0.000382 | 231 |
| Npy | 109648 | -0.69977 | 5.24E-06 | 232 |
| Rpl38 | 67671 | -0.69998 | 0.002484 | 233 |

| | | | | |
|---|---|---|---|---|
| Ndufa3 | 66091 | -0.7001 | 5.17E-05 | 234 |
| Polr2f | 69833 | -0.70437 | 2.60E-05 | 235 |
| Ccdc171 | 320226 | -0.70697 | 0.035094 | 236 |
| Smim18 | 72632 | -0.71103 | 0.000243 | 237 |
| Rimklb | 108653 | -0.71179 | 0.000001 | 238 |
| Coa6 | 67892 | -0.71399 | 0.000738 | 239 |
| Yjefn3 | 234365 | -0.72021 | 0.000551 | 240 |
| Cox7b | 66142 | -0.72105 | 0.000001 | 241 |
| Gtf2h3 | 209357 | -0.72699 | 9.74E-05 | 242 |
| Uqcr11 | 66594 | -0.72745 | 0.000001 | 243 |
| Rac3 | 170758 | -0.72792 | 0.003008 | 244 |
| Ndufa13 | 67184 | -0.72887 | 0.012388 | 245 |
| Uqcrq | 22272 | -0.72946 | 0.000001 | 246 |
| 1810009A15Rik | 66276 | -0.73222 | 0.023753 | 247 |
| 2510002D24Rik | 72307 | -0.73334 | 0.010522 | 248 |
| Mrpl34 | 94065 | -0.73799 | 4.90E-05 | 249 |
| Mrpl53 | 68499 | -0.74178 | 0.000138 | 250 |
| Txndc17 | 52700 | -0.74441 | 0.032922 | 251 |
| Usmg5 | 66477 | -0.75388 | 0.000001 | 252 |
| Atp5j | 11957 | -0.75725 | 2.02E-06 | 253 |
| Ndufb7 | 66916 | -0.76478 | 6.23E-05 | 254 |
| 8430429K09Rik | 71523 | -0.76933 | 0.007871 | 255 |
| Cox5b | 12859 | -0.76939 | 0.000001 | 256 |
| Pfdn1 | 67199 | -0.77147 | 2.74E-05 | 257 |
| Polr2j | 20022 | -0.7753 | 9.57E-06 | 258 |
| Ngfrap1 | 12070 | -0.77613 | 0.000304 | 259 |
| Dnajc19 | 67713 | -0.78268 | 0.022962 | 260 |
| Atpif1 | 11983 | -0.7903 | 2.06E-06 | 261 |
| Atp5j2 | 57423 | -0.79106 | 1.91E-06 | 262 |
| Aldh3b2 | 621603 | -0.79546 | 0.001353 | 263 |
| 2010107E04Rik | 70257 | -0.79974 | 2.93E-05 | 264 |
| Gm6741 | 627235 | -0.80269 | 0.016107 | 265 |
| Nans | 94181 | -0.80417 | 1.12E-05 | 266 |
| Fkbp2 | 14227 | -0.80456 | 0.008212 | 267 |
| Tomm7 | 66169 | -0.81366 | 0.000001 | 268 |
| Pgam2 | 56012 | -0.82302 | 0.006719 | 269 |
| Cox20 | 66359 | -0.83837 | 0.03382 | 270 |
| Ggct | 110175 | -0.84694 | 0.041823 | 271 |
| Cort | 12854 | -0.8517 | 0.044239 | 272 |
| Vip | 22353 | -0.85262 | 0.024373 | 273 |
| Rpl10a-ps1 | 1E+08 | -0.85364 | 0.000001 | 274 |
| Rpl10a | 19896 | -0.86055 | 0.000001 | 275 |
| Atp5k | 11958 | -0.86231 | 0.000811 | 276 |
| 2310009A05Rik | 66364 | -0.8947 | 0.02839 | 277 |
| Ndufa5 | 68202 | -0.90044 | 3.45E-05 | 278 |
| Anapc13 | 69010 | -0.90534 | 0.008103 | 279 |
| Cox17 | 12856 | -0.90572 | 0.00234 | 280 |

| | | | | |
|---|---|---|---|---|
| 1110017D15Rik | 73721 | -0.91158 | 0.034011 | 281 |
| 1700007K13Rik | 69327 | -0.92278 | 0.026644 | 282 |
| Cox6c | 12864 | -0.94107 | 9.18E-05 | 283 |
| Rps27a-ps2 | 619900 | -0.94599 | 0.005575 | 284 |
| Mrps18c | 68735 | -0.95459 | 2.63E-06 | 285 |
| Ndufb4 | 68194 | -0.9548 | 0.000001 | 286 |
| Xlr3b | 574437 | -0.99847 | 0.015067 | 287 |
| Dpm1 | 13480 | -1.02352 | 0.000394 | 288 |
| Gm3696 | 1E+08 | -1.02664 | 0.008219 | 289 |
| Pdcd5 | 56330 | -1.02863 | 0.002968 | 290 |
| Nms | 433292 | -1.04067 | 0.014662 | 291 |
| Pin4 | 69713 | -1.06095 | 0.014224 | 292 |
| Gm2464 | 1E+08 | -1.10824 | 0.020435 | 293 |
| Gng13 | 64337 | -1.115 | 0.001997 | 294 |
| Prss57 | 73106 | -1.35814 | 0.039602 | 295 |
| Fbxl12os | 66662 | -1.39059 | 2.44E-05 | 296 |
| Cdh23 | 22295 | -1.41124 | 0.000983 | 297 |
| Gm11346 | 76024 | -1.57816 | 0.00076 | 298 |
| Papl | 101744 | -1.60508 | 0.03049 | 299 |
| Rnaseh2c | 68209 | -1.68899 | 0.000001 | 300 |
| Ppp1r3fos | 78185 | -1.73064 | 0.008614 | 301 |
| Rpl15-ps3 | 1E+08 | -1.84538 | 0.00458 | 302 |
| Gm13292 | 1E+08 | -2.32608 | 0.028884 | 303 |
| Gm3383 | 1E+08 | -3.00405 | 0.000145 | 304 |
| Vdac3-ps1 | 22336 | -3.01517 | 0.000001 | 305 |
| Gm15927 | 1.03E+08 | -3.86199 | 0.007251 | 306 |
| Igkv2-109 | 628268 | -4.53396 | 0.002329 | 307 |
| Gm10409 | 1E+08 | -9.25452 | 4.73E-06 | 308 |
| Zbed6 | 667118 | -10 | 1.91E-05 | 309 |
