## Supplementary Table 3 for "Lateral Septal Circuits Govern Schizophrenia-Like Effects of Ketamine on Social Behavior"

### Supplementary Table 3 - Notable Dysregulated Genes By Category

| Category | Symbol | Entrez | Log2fc | Adjpv | Dysregulation |
|---|---|---|---|---|---|
| Apoptotic Processes | Anxa2 | 12306 | 1.2919967 | 2.89E-06 | Upregulated |
| | Anxa4 | 11746 | 1.9437055 | 2.02E-06 | Upregulated |
| | Birc3 | 11796 | 1.2618424 | 0.00144 | Upregulated |
| | Casp1 | 12362 | 0.9982776 | 0.0052342 | Upregulated |
| | Cd82 | 12521 | 1.2034514 | 1.91E-05 | Upregulated |
| | Hspa1a | 193740 | 1.4019191 | 0.000001 | Upregulated |
| | Ifitm3 | 66141 | 1.3213261 | 5.97E-05 | Upregulated |
| | Meg3 | 17263 | 0.8119243 | 0.00025838 | Upregulated |
| | Parp10 | 671535 | 1.6093393 | 0.000001 | Upregulated |
| | Parp12 | 243771 | 1.9675469 | 0.00248546 | Upregulated |
| | Xaf1 | 327959 | 1.0561731 | 0.00080516 | Upregulated |
| | Zc3hav1 | 78781 | 1.8621174 | 0.00095897 | Upregulated |
| Encoding Potassium Inwardly-Rectifying (Kir) Channels | Kcnj10 | 16513 | 0.9630137 | 0.00011485 | Upregulated |
| | Kcnj13 | 100040591 | 2.4586514 | 0.00194637 | Upregulated |
| | Kcnk13 | 217826 | 0.5048098 | 0.0228932 | Upregulated |
| | Kcnk6 | 52150 | 0.9675011 | 0.02075376 | Upregulated |
| Encoding Solute Transporters | Slc11a1 | 18173 | 1.5881121 | 3.47E-05 | Upregulated |
| | Slc12a2 | 20496 | 0.7864146 | 0.000001 | Upregulated |
| | Slc12a4 | 20498 | 1.3191762 | 1.27E-06 | Upregulated |
| | Slc12a7 | 20499 | 1.2787037 | 0.00016128 | Upregulated |
| | Slc14a1 | 108052 | 1.1678939 | 0.01052187 | Upregulated |
| | Slc15a3 | 65221 | 1.5504177 | 0.00356413 | Upregulated |
| | Slc17a6 | 140919 | 0.5259342 | 0.04711297 | Upregulated |
| | Slc1a5 | 20514 | 1.6190717 | 0.00728799 | Upregulated |
| | Slc29a3 | 71279 | 1.40711 | 3.99E-05 | Upregulated |
| | Slc37a2 | 56857 | 0.9632701 | 0.02605571 | Upregulated |
| | Slc38a3 | 76257 | 1.3169695 | 0.04358236 | Upregulated |
| | Slc39a12 | 277468 | 1.2138449 | 0.00165133 | Upregulated |
| | Slc43a3 | 58207 | 2.0679815 | 0.00236365 | Upregulated |
| | Slc44a1 | 100434 | 0.5332562 | 3.16E-06 | Upregulated |
| | Slc4a2 | 20535 | 0.9966346 | 0.000001 | Upregulated |
| | Slc4a4 | 54403 | 0.9803397 | 0.0373076 | Upregulated |
| | Slc7a11 | 26570 | 0.9005024 | 0.000001 | Upregulated |
| | Slc7a2 | 11988 | 1.520138 | 1.64E-06 | Upregulated |
| | Slc8b1 | 170756 | 1.2813317 | 0.0021072 | Upregulated |
| | Slc9a3r1 | 26941 | 1.0284477 | 0.00027058 | Upregulated |

| | | | | | |
|---|---|---|---|---|---|
| | Slco1a4 | 28250 | 1.1753749 | 0.00579739 | Upregulated |
| | Slco1c1 | 58807 | 1.3408565 | 0.000001 | Upregulated |
| | Slco2a1 | 24059 | 1.3686218 | 0.04214201 | Upregulated |
| | Slco2b1 | 101488 | 1.2182087 | 0.00471782 | Upregulated |
| Epigenetic Reader and Modifier Genes | Jmjd1c | 108829 | 0.7565758 | 0.000001 | Upregulated |
| | Prrc2c | 226562 | 0.8606207 | 0.000001 | Upregulated |
| Extracellular Matrix Remodeling | Col4a1 | 12826 | 1.800855 | 0.000001 | Upregulated |
| | Col6a3 | 12835 | 1.1274468 | 0.000001 | Upregulated |
| | Enpp2 | 18606 | 1.6747707 | 0.000001 | Upregulated |
| | Flnb | 286940 | 2.2631139 | 0.000001 | Upregulated |
| | Hspg2 | 15530 | 3.4216293 | 0.000001 | Upregulated |
| | Itgb4 | 192897 | 2.4177507 | 0.000001 | Upregulated |
| | Lamb2 | 16779 | 2.0560907 | 0.000001 | Upregulated |
| | Mag | 17136 | 1.7160423 | 0.000001 | Upregulated |
| | Sulf1 | 240725 | 0.9469175 | 0.000001 | Upregulated |
| | Zeb2 | 24136 | 0.939002 | 0.000001 | Upregulated |
| Myelination or Oligodendrocyte Function | Cldn11 | 18417 | 1.3473386 | 0.000001 | Upregulated |
| | Enpp2 | 18606 | 1.6747707 | 0.000001 | Upregulated |
| | Ermn | 77767 | 1.2957806 | 0.000001 | Upregulated |
| | Flnb | 286940 | 2.2631139 | 0.000001 | Upregulated |
| | Mag | 17136 | 1.7160423 | 0.000001 | Upregulated |
| | Mast4 | 328329 | 0.7362559 | 0.000001 | Upregulated |
| | Meg3 | 17263 | 0.8119243 | 0.00025838 | Upregulated |
| | Myrf | 225908 | 1.7186017 | 0.000001 | Upregulated |
| | Plp1 | 18823 | 1.0747787 | 0.000001 | Upregulated |
| | Ugt8a | 22239 | 1.2656413 | 0.000001 | Upregulated |
| Neuroinflammation/Immune Response | Anxa2 | 12306 | 1.2919967 | 2.89E-06 | Upregulated |
| | Anxa4 | 11746 | 1.9437055 | 2.02E-06 | Upregulated |
| | Casp1 | 12362 | 0.9982776 | 0.0052342 | Upregulated |
| | Helz2 | 229003 | 2.9974814 | 0.000001 | Upregulated |
| | Ighv14-2 | 668421 | 9.2182288 | 0.02157011 | Upregulated |
| | Ighv5-16 | 633568 | 8.3712974 | 2.66E-05 | Upregulated |
| | Ighv9-3 | 780825 | 8.296644 | 0.00148355 | Upregulated |
| | Igkv3-4 | 626347 | 10 | 0.01516193 | Upregulated |
| | Igkv9-123 | 628144 | 10 | 0.01065665 | Upregulated |
| | Mx1 | 17857 | 1.918132 | 0.000001 | Upregulated |
| | Nlrp3 | 216799 | 1.6432315 | 0.01652913 | Upregulated |
| | Nos2 | 18126 | 3.9928871 | 0.000001 | Upregulated |
| | Nr4a2 | 18227 | 0.5956207 | 0.02192992 | Upregulated |

| | | | | | |
|---|---|---|---|---|---|
| Risk of Developmental Disorders | Ebf3 | 13593 | 1.6569205 | 0.000001 | Upregulated |
| | Hspg2 | 15530 | 3.4216293 | 0.000001 | Upregulated |
| | Itgb4 | 192897 | 2.4177507 | 0.000001 | Upregulated |
| | Rbm47 | 245945 | 2.5152695 | 0.000001 | Upregulated |
| Mitochondrial Function | Atp5e | 67126 | -0.603538 | 1.67E-05 | Downregulated |
| | Atp5g1 | 11951 | -0.574645 | 0.000001 | Downregulated |
| | Atp5h | 71679 | -0.52715 | 0.000001 | Downregulated |
| | Atp5j | 11957 | -0.757252 | 2.02E-06 | Downregulated |
| | Atp5j2 | 57423 | -0.791058 | 1.91E-06 | Downregulated |
| | Atp5k | 11958 | -0.862309 | 0.00081071 | Downregulated |
| | Atp5l | 27425 | -0.612889 | 0.000001 | Downregulated |
| | Atp6v1f | 66144 | -0.565864 | 0.000001 | Downregulated |
| | Atpif1 | 11983 | -0.790302 | 2.06E-06 | Downregulated |
| | Ndufb11 | 104130 | -0.551978 | 7.28E-06 | Downregulated |
| | Ndufb3 | 66495 | -0.522497 | 0.00030391 | Downregulated |
| | Ndufb4 | 68194 | -0.954795 | 0.000001 | Downregulated |
| | Ndufb5 | 66046 | -0.502984 | 1.20E-05 | Downregulated |
| | Ndufb6 | 230075 | -0.604287 | 3.75E-06 | Downregulated |
| | Ndufb7 | 66916 | -0.764779 | 6.23E-05 | Downregulated |
| | Ndufb8 | 67264 | -0.600275 | 0.000001 | Downregulated |
| | Ndufb9 | 66218 | -0.548773 | 6.00E-06 | Downregulated |
| | Vdac3-ps1 | 22336 | -3.015171 | 0.000001 | Downregulated |
