## Supplementary Table 4 for "Lateral Septal Circuits Govern Schizophrenia-Like Effects of Ketamine on Social Behavior"

**Supplementary Table 4 - Overlaps of differentially expressed genes (DEGs) between chronic ketamine-treated mice and schizophrenic individuals (Merikangas et al., 2022)**

| Sympol | Entrez | Log2FC | P-Value | Expression |
|---|---|---|---|---|
| Gbp2 | 14469 | **2.799** | **3.62E-04** | Upregulated |
| Birc3 | 11796 | **1.262** | **0.001** | Upregulated |
| Ifitm1 | 68713 | **1.1** | 0.323 | Upregulated |
| Ifitm2 | 80876 | **0.814** | 0.153 | Upregulated |
| Ifitm3 | 66141 | **1.321** | **5.97E-05** | Upregulated |
| Mov10 | 17454 | **1.322** | **1.00E-06** | Upregulated |
| Anxa4 | 11746 | **1.944** | **2.02E-06** | Upregulated |
| Anxa2 | 12306 | **1.292** | **2.89E-06** | Upregulated |
| Cd82 | 12521 | **1.203** | **1.91E-05** | Upregulated |
| Hspa1a | 193740 | **1.402** | **1.00E-06** | Upregulated |
| Itga5 | 16402 | **1.609** | **0.001** | Upregulated |
| Nuak2 | 74137 | **1.057** | **0.008** | Upregulated |
| Oas2 | 246728 | **1.843** | **0.001** | Upregulated |
| Parp10 | 671535 | **1.609** | **1.00E-06** | Upregulated |
| Parp12 | 243771 | **1.968** | **0.002** | Upregulated |
| Pdgfrb | 18596 | **1.225** | **3.78E-04** | Upregulated |
| Rrbp1 | 81910 | **1.22** | **6.79E-05** | Upregulated |
| Srgn | 19073 | **1.545** | **0.01** | Upregulated |
| Tcn2 | 21452 | **1.361** | **6.42E-04** | Upregulated |
| Tgm2 | 21817 | **2.305** | **1.00E-06** | Upregulated |
| Xaf1 | 327959 | **1.056** | **8.05E-04** | Upregulated |
| Zc3hav1 | 78781 | **1.862** | **9.59E-04** | Upregulated |
| Cd46 | 17221 | **0.977** | 0.343 | Upregulated |
| Fbxo32 | 67731 | **0.778** | **2.35E-04** | Upregulated |
| Meg3 | 17263 | **0.812** | **2.58E-04** | Upregulated |
| Nr4a1 | 15370 | **1.264** | 0.334 | Upregulated |
| Casp1 | 12362 | **0.998** | **0.005** | Upregulated |
| Thoc7 | 66231 | -0.457 | **0.001** | Downregulated |
| Dynlt1a | 100310872 | **-0.688** | 0.198 | Downregulated |
| Med28 | 66999 | -0.468 | **1.32E-05** | Downregulated |
| Ypel3 | 66090 | -0.442 | **1.64E-06** | Downregulated |
| Apopt1 | 68020 | -0.468 | **0.003** | Downregulated |
| Gabra5 | 110886 | -0.365 | **0.002** | Downregulated |
| Ndufb2 | 68198 | -0.476 | **5.08E-05** | Downregulated |
| Nrgn | 64011 | **-0.624** | **3.11E-06** | Downregulated |
| Cabp1 | 29867 | -0.419 | **0.032** | Downregulated |
| Hint1 | 15254 | -0.49 | **1.00E-06** | Downregulated |
| Vdac3 | 22335 | -0.329 | **0.001** | Downregulated |

DEG inclusion criteria: Log2FC > 0.5, or p < 0.05
