## Supplementary Table 5 for "Lateral Septal Circuits Govern Schizophrenia-Like Effects of Ketamine on Social Behavior"

**Supplementary Table 5 - Top 50 Highly Expressed Genes in the Saline Group**

| Ranking | Gene Name |
|---|---|
| 1 | Yam1 |
| 2 | Atp1a3 |
| 3 | Calm1 |
| 4 | Ncdn |
| 5 | Actb |
| 6 | Ywhaz |
| 7 | Hspa8 |
| 8 | Atp1b1 |
| 9 | Gapdh |
| 10 | Gnas |
| 11 | App |
| 12 | Ndrg4 |
| 13 | Eef1a1 |
| 14 | Aldoa |
| 15 | Snap25 |
| 16 | Camk2a |
| 17 | Syt1 |
| 18 | Ywhag |
| 19 | Ubb |
| 20 | Psap |
| 21 | Syp |
| 22 | Map1b |
| 23 | Atp2a2 |
| 24 | Vamp2 |
| 25 | Spock2 |
| 26 | Actg1 |
| 27 | Ttc3 |
| 28 | Map2 |
| 29 | Calm3 |
| 30 | Calm2 |
| 31 | Gad2 |
| 32 | Hsp90aa1 |
| 33 | Arf3 |
| 34 | Aplp1 |
| 35 | Dpysl2 |
| 36 | Kif1a |
| 37 | Zwint |
| 38 | Agap2 |
| 39 | Peg3 |
| 40 | Rtn3 |
| 41 | Hsp90ab1 |
| 42 | Atp5b |
| 43 | Vsnl1 |
| 44 | Eno2 |
| 45 | Dnm1 |

| | |
|---|---|
| 46 | Clstn1 |
| 47 | Rtn1 |
| 48 | Prkar1a |
| 49 | Gad1 |
| 50 | Gabbr1 |
